# Self-organizing human heart assembloids with autologous and developmentally relevant cardiac neural crest-derived tissues

**DOI:** 10.1101/2024.12.11.627627

**Authors:** Aleksandra Kostina, Artem Kiselev, Amanda Huang, Haley Lankerd, Sammantha Caywood, Ariadna Jurado-Fernandez, Brett Volmert, Colin O’Hern, Aniwat Juhong, Yifan Liu, Zhen Qiu, Sangbum Park, Aitor Aguirre

## Abstract

Neural crest cells (NCCs) are a multipotent embryonic cell population of ectodermal origin that extensively migrate during early development and contribute to the formation of multiple tissues. Cardiac NCCs play a critical role in heart development by orchestrating outflow tract septation, valve formation, aortic arch artery patterning, parasympathetic innervation, and maturation of the cardiac conduction system. Abnormal migration, proliferation, or differentiation of cardiac NCCs can lead to severe congenital cardiovascular malformations. However, the complexity and timing of early embryonic heart development pose significant challenges to studying the molecular mechanisms underlying NCC-related cardiac pathologies. Here, we present a sophisticated functional model of human heart assembloids derived from induced pluripotent stem cells, which, for the first time, recapitulates cardiac NCC integration into the human embryonic heart *in vitro*. NCCs successfully integrated at developmentally relevant stages into heart organoids, and followed developmental trajectories known to occur in the human heart. They demonstrated extensive migration, differentiated into cholinergic neurons capable of generating nerve impulses, and formed mature glial cells. Additionally, they contributed to the mesenchymal populations of the developing outflow tract. Through transcriptomic analysis, we revealed that NCCs acquire molecular features of their cardiac derivatives as heart assembloids develop. NCC-derived parasympathetic neurons formed functional connections with cardiomyocytes, promoting the maturation of the cardiac conduction system. Leveraging this model’s cellular complexity and functional maturity, we uncovered that early exposure of NCCs to antidepressants harms the development of NCC derivatives in the context of the developing heart. The commonly prescribed antidepressant Paroxetine disrupted the expression of a critical early neuronal transcription factor, resulting in impaired parasympathetic innervation and functional deficits in cardiac tissue. This advanced heart assembloid model holds great promise for high-throughput drug screening and unraveling the molecular mechanisms underlying NCC-related cardiac formation and congenital heart defects.

**IN BRIEF:** Human neural crest heart assembloids resembling the major directions of neural crest differentiation in the human embryonic heart, including parasympathetic innervation and the mesenchymal component of the outflow tract, provide a human-relevant embryonic platform for studying congenital heart defects and drug safety.

## INTRODUCTION

Cardiovascular conditions are the leading cause of morbidity and mortality in developed countries, with congenital heart defects being the most common birth anomalies in humans^1–3^. The complex and multifaceted interplay of genetic, developmental, environmental, and lifestyle factors underlies cardiovascular disorders, posing significant challenges in unraveling their mechanisms and advancing effective treatments. Transgenic and knockout animal models remain indispensable tools and the gold standard for studying the roles of specific genes and cardiac cell populations in embryonic heart formation^4–8^. However, critical interspecies differences in anatomy, physiology, genetics, and development limit the direct extrapolation of findings to humans^9–14^. This emphasizes the importance of integrating animal models with human-based *in vitro* systems to enhance translational relevance and cross-validation. Recently, we and others reported the creation of three-dimensional, self-organizing, pluripotent stem cell-derived human heart organoids (hHOs)^15–18^.These developmentally accurate “mini-hearts” exhibit human heart-like cellular complexity, anatomy and physiology, effectively recapitulating early stages of human heart development. The field of hHOs has grown rapidly, with significant advancements in maturation strategies^19^, morphological complexity^20^, chamber formation^21^, and integration with other organoid systems^22–24^. These developments aim to replicate cardiogenesis better, creating in vitro heart models suitable for studying embryonic and adult cardiovascular pathologies and exploring novel treatments and preventive strategies for cardiovascular diseases^25^.

While the heart is predominantly of mesodermal origin and is formed predominantly from first and second heart field cardiac progenitors, several extracardiac cell populations are essential for proper heart formation and function^26^. Most recently, embryonic macrophages have been successfully incorporated into hHOs, creating the human heart-macrophage assembloids that remarkably mimic cardiac tissue-resident macrophages in the human embryonic heart^27^. Another extracardiac population is neural crest cells (NCCs), which are a crucial multipotent cell population that exists exclusively during early embryogenesis. These cells delaminate from the neural tube and migrate extensively throughout the embryo, contributing to a wide range of tissues and organs, including craniofacial cartilage and skeleton, melanocytes, the enteric nervous system, and the developing heart^28^. Cardiac NCCs, a subset of vagal NCCs, play critical roles in remodeling pharyngeal arch arteries, septating the outflow tract (OFT), driving valvulogenesis, enabling parasympathetic cardiac innervation, and supporting conduction system maturation^29^. While these structures are not entirely composed of NCCs, their development critically depends on NCC contributions. Disrupted migration and differentiation of cardiac NCCs result in congenital heart defects such as atrial and ventricular septal defects and OFT malformations (e.g., Tetralogy of Fallot, transposition of the great arteries, aortic or pulmonary artery stenosis, bicuspid aortic valve, patent ductus arteriosus, and persistent truncus arteriosus)^30–32^.

Heart development begins early in embryogenesis, with NCCs migrating to the developing heart in humans during the fourth week of gestation. This early timeline contributes to the elusive nature of congenital cardiovascular neurocristopathies. The OFT is formed by second heart field progenitors that contribute to the myocardial part, while the NCC-derived mesenchymal component provides essential patterning^33,34^. Although the neural crest contributes a relatively small number of cells compared to the cardiogenic plate, their presence in the OFT is indispensable for normal development^35^. Studies emphasize the primary role of NCCs in cardiovascular development, but the extent of their contributions varies across model systems and species. This highlights the need for highly human-relevant platforms to investigate NCC roles in heart development and their contributions to cardiovascular diseases. Despite the high complexity of existing 3D heart organoid models, none have incorporated NCC migration or tissues, limiting the ability to study processes and pathologies associated with neural crest contributions. While the myocardial portion of the OFT has been modeled separately^21^, models integrating the NCC-derived mesenchymal component, crucial for OFT formation, still need to be developed.

Treating maternal conditions during early pregnancy is particularly challenging, as many medications can severely impact the developing embryo. Depressive disorders, which are increasingly prevalent among women of reproductive age, are commonly treated with selective serotonin reuptake inhibitors (SSRIs)^36^. SSRIs, which block serotonin reuptake and increase extracellular serotonin concentrations, readily cross the placenta^37^. This results in the modulation of embryonic serotonin levels, disrupting its role as a critical signaling molecule during development^38,39^. Accumulating epidemiological evidence links SSRI use during pregnancy to congenital heart malformations in the fetus^40–43^, but human-relevant models are urgently needed to dissect these potential risks during early heart development.

Here, we describe a human neural crest-heart assembloid (hNCHA) platform that closely mimics the in vivo migration of cardiac NCCs into the embryonic heart and their differentiation into major cardiac NCC derivatives in a highly controlled environment. This innovative platform functionally models NCC integration and uncovers the deleterious effects of early SSRI exposure on the formation of critical cardiac structures in the embryonic heart. The hNCHA platform represents a valuable tool for advancing our understanding of neural crest biology in heart development, and provides a unique opportunity to investigate NCC-related congenital heart defects and the impact of pharmacological interventions during early pregnancy.

## RESULTS

### Human Neural Crest-Heart Assembloids generated by integration of autologous hiPSC-Derived Neural Crest Cells into Human Heart Organoids

To create human heart assembloids containing autologous cardiac derivatives of neural crest cells (NCCs), we first generated NCCs from human induced pluripotent stem cells (iPSCs) using a previously reported method that involves inhibiting TGFβ and GSK3^44^. The same hiPSC line was engineered to stably express mCherry using lentiviral transduction, and was used for independent NCC differentiation (mCherry-NCCs), producing autologous labeled NCCs that are easy to detect and monitor for cell migration. Sustained culture for 7 days resulted in expression of the surface marker *NGFR* and transcriptional factors *SOX10*, *TFAP2A*, and *FOXD3*, all associated with neural crest identity **(Supp. Fig. 1A, B)**. We further tested whether hiPSC-derived NCCs mimic developmental migratory behavior. The NCCs were dissociated into single cells and reaggregated in ultra-low attachment plates by centrifugation. After 24 hr, NCCs formed neurospheres that, when plated onto Matrigel, attached, and SOX10^+^ cells migrated out as if from a neural tube upon development **(Supp. Fig 1C, D)**.

Class 3 semaphorin, specifically the ligand Semaphorin 3C (*SEMA3C*), is expressed in myocardial cells and the mesenchyme of the developing heart, providing attractive cues that direct NCC migration through its interaction with the transmembrane receptor Plexin A2 (*PLXNA2*)^45–47^. We analyzed the dynamics of *SEMA3C* expression in human heart organoids (hHOs) at various differentiation time points (day 0 through day 19) using previously published transcriptomic profiles^15^. *SEMA3C* showed peak expression on Day 17 of hHO development **(Supp. Fig 1E)**. We also confirmed *PLXNA2* expression in iPSC-derived NCCs by qRT-PCR **(Supp. Fig 1F)**. Then, we used Day 17 hHOs to integrate *PLXNA2^+^*NCCs to engineer Human Neural Crest Heart Assembloids (hNCHAs) - an *in vitro* system that fully recapitulates the embryonic migration of NCCs in the developing heart. **Figure 1A** provides a schematic summary of the protocol for hNCHA generation. Early embryonic self-organizing and developmentally relevant hHOs were differentiated and maintained until Day 17 following our previously established protocol^15^. Two days before integration (Day -2), 2,000 mCherry-NCCs were seeded in a V-bottom, low-attachment 96-well plate with NCC maintenance media **(Fig. 1B, top left)**. After 24 hours (Day -1), the NCCs formed mCherry-labeled neurospheres **(Fig. 1B, top right)**. On the integration day (Day 0), Day 17 hHOs were transferred to the NCC-formed neurospheres in the V-bottom plate to create human neural crest heart assembloids (hNCHAs) **(Fig. 1B, bottom left; Suppl. Fig. 1G)**. mCherry- NCCs were observed migrating effectively into the hNCHAs as early as 48 hours post-integration **(Supp. Fig. 1H; Supp. Video 1; Supp. Video 2; Supp. Video 3; Supp. Video 4; Supp. Video 5)**. Twelve days after integration, mCherry-NCCs migrated throughout the hNCHAs, forming neuronal-like projections **(Fig. 1B, bottom right; Supp.** Fig. 1I**; Supp. Video 6; Supp. Video 7; Supp. Video 8; Supp. Video 9)**.

**Figure 1.**
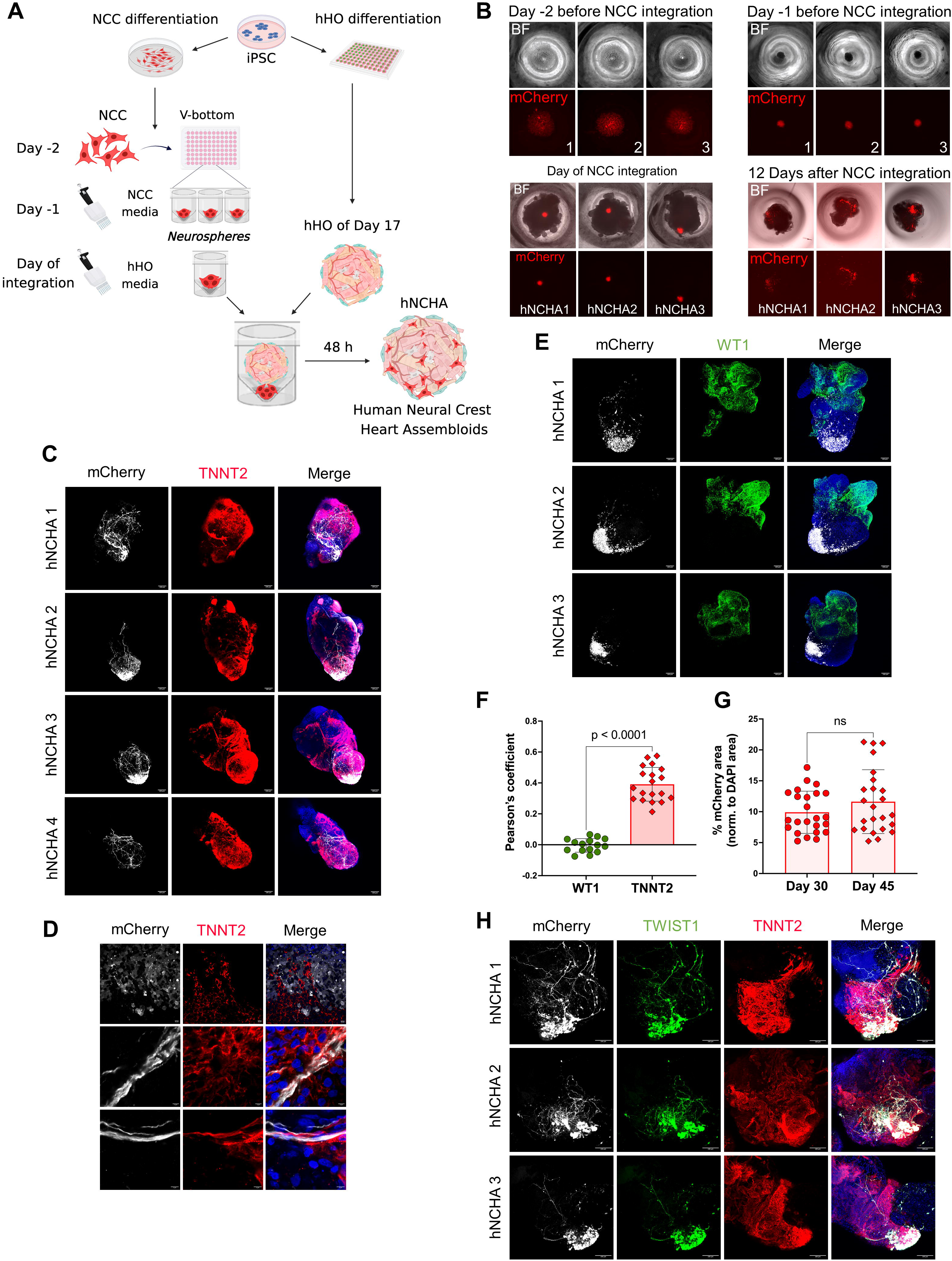
(A) A schematic representation of the protocol for creating human neural crest heart assembloids (hNCHAs). (B) Brightfield (BF) and fluorescent images of neural crest cells with stable mCherry expression (mCherry-NCCs) (top) and human neural crest heart assembloids (hNCHAs) (bottom). Three representative wells are shown. (C) Representative immunofluorescence images of four Day 30 hNCHAs displaying mCherry-NCCs (grey), cardiomyocyte marker TNNT2 (red) and the nuclear marker DAPI (blue); Scale bar = 100 mm. (D) High magnification images of three hNCHAs displaying mCherry-NCCs (grey), cardiomyocyte marker TNNT2 (red) and the nuclear marker DAPI (blue); scale bar = 10 mm. Scale bar = 10 mm (top row) and 5 mm (middle, bottom rows). (E) Representative immunofluorescence images of three Day 30 hNCHAs displaying mCherry-NCCs (grey), epicardial marker WT1 (green) and the nuclear marker DAPI (blue); Scale bar = 100 mm. (F) Quantification of colocalization (Pearson’s coefficient) between NCCs (mCherry) and cardiomyocytes (TNNT2) or epicardial cells (WT1) respectively measured by image analysis; n = 19 for TNNT2, n = 15 for WT1. Values = mean ± SD, unpaired t test. (G) Quantification of mCherry^+^ area in Day 30 and Day 45 hNCHAs measured by image analysis; n = 24. Values are presented as a percentage of DAPI^+^ area. Values = mean ± SD, unpaired t test. (H) Representative immunofluorescence images of three Day 30 hNCHAs displaying mCherry-NCCs (grey), marker of migrating NCCs TWIST1 (green) and the nuclear marker DAPI (blue). Scale bar = 100 mm.

To determine the localization of migrating NCCs in hNCHAs and clarify the spatial relationship between NCCs and other cell types, we performed immunofluorescence staining for the cardiomyocyte-specific marker TNNT2 and the epicardium marker WT1. In Day 30 hNCHAs (13 days after NCC integration), mCherry-NCCs and growing neuronal projections were predominantly colocalized with myocardial tissue **(Fig. 1C)**. High-magnification images of the assembloids showed mCherry-NCCs migrating along cardiomyocytes and neuronal projections closely interacting with them, implying the expression of attractive cues by cardiomyocytes **(Fig. 1D; Supp.** Fig. 1J). In contrast, mCherry-NCCs were found localized on the opposite pole from the area densely covered by WT1^+^ epicardial cells **(Fig. 1E)**. We quantified the area of colocalization between mCherry^+^ cells and either TNNT2^+^ or WT1^+^ regions, confirming that migrating NCCs in hNCHAs are significantly more associated with cardiomyocytes **(Fig. 1F)**. Quantification of the total mCherry^+^ area showed an NCC and derivative presence of 5 to 15% in hNCHAs, with no significant change between assembloids at Day 30 and Day 45 **(Fig. 1G)**. This data demonstrates that integrated NCCs persist in hNCHAs throughout their ongoing development. Immunofluorescence assessment for the key NCC transcription factor *TFAP2A* demonstrated a complete absence of NCCs in non-integrated hHOs **(Supp. Fig. 1K)**. The basic Helix-loop-Helix *TWIST1* is a key transcription factor orchestrating cardiac NCC initial formation and migration as well as their ability to differentiate into diverse cell types^48–50^. Immunofluorescence assessment of hNCHAs revealed abundant *TWIST1* expression in migrating mCherry-NCCs **(Fig. 1H)**. In summary, we created hNCHAs that recapitulate the efficient integration and migration of cardiac NCCs during early heart development and demonstrate active interactions between migrating NCCs and myocardial cells. The migrating NCCs contribute to a specific morphological organization within the heart assembloids and begin differentiating into NCC derivatives, developing neurite-like filaments.

### Single-cell RNA sequencing of hNCHAs reveals developmental paths of cardiac NCCs and transcriptional changes in major cardiac cell types

To broadly characterize the cellular composition and transcriptomic landscape of hNCHAs compared to hHOs without NCCs, we performed single-cell RNA sequencing (scRNA-seq) on hHOs and hNCHAs at Day 29 (12 days after NCC integration) and Day 43 (26 days after NCC integration). Unsupervised dimensionality reduction of the scRNA-seq data using UMAP identified 18 and 19 cell clusters in hHOs and hNCHAs, respectively representing various cell lineages characteristic of the human embryonic heart **(Fig. 2A; Supp.** Fig. 2A). Cell lineages were defined based on the expression of established markers^16,19,51–57^ or through the analysis described below. Representative molecular signatures for each cluster are shown in **Fig. 2B** and **Supplementary Table 3**. Neural crest cells and their derivatives in hNCHAs were identified based on mCherry expression and correspond to cluster 9. Remarkably, the NCC cluster in Day 29 hNCHAs was subdivided into two populations, with partial cell localization at the border of cluster 1 and cluster 3, representing a transitioning mesenchymal population (MESC) and a cell lineage with an OFT signature (OFT), respectively. This finding suggests that integrated NCCs acquire multiple cell fates during migration in hNCHAs. To obtain the transcriptomic profile of NCCs before integration, 10% mCherry-NCCs were added to Day 29 hHO sample during preparation for sequencing. The mCherry signature enabled the separation of NCCs in data analysis. Thus, cluster 9 in Day 29 hHO should be considered as control NCCs before integration. A more detailed description of scRNA-seq sample preparation and data processing is provided in the Methods. We then compared hNCHA transcriptional profile with an scRNA-seq dataset derived from a 9-week post-conception human embryonic heart **(Fig. 2C)**^55^. Integration of the developing human heart and hNCHA datasets revealed a phenomenal overlap across most cell types, highlighting the high level of similarity between engineered hNCHAs and the early embryonic heart. As expected, the NCC population mapped primarily to the neuronal cluster and partially to fibroblast-like cells from the human heart. Interestingly, right ventricular cardiomyocytes (RV CMs) from hNCHAs colocalized with atrial cardiomyocytes (aCMs) from the human heart, reflecting cardiomyocyte specification based on heart field origin: the first heart field gives rise to the left ventricle, while the right ventricle and atria develop from the second heart field^58^. The cluster of cells from hNCHAs labeled ’Others,’ containing a mixture of stromal cells, closely overlapped with white blood cells (WBCs), possibly due to shared transcriptional pathways and potential interactions between stromal and immune cells^59^.

**Figure 2.**
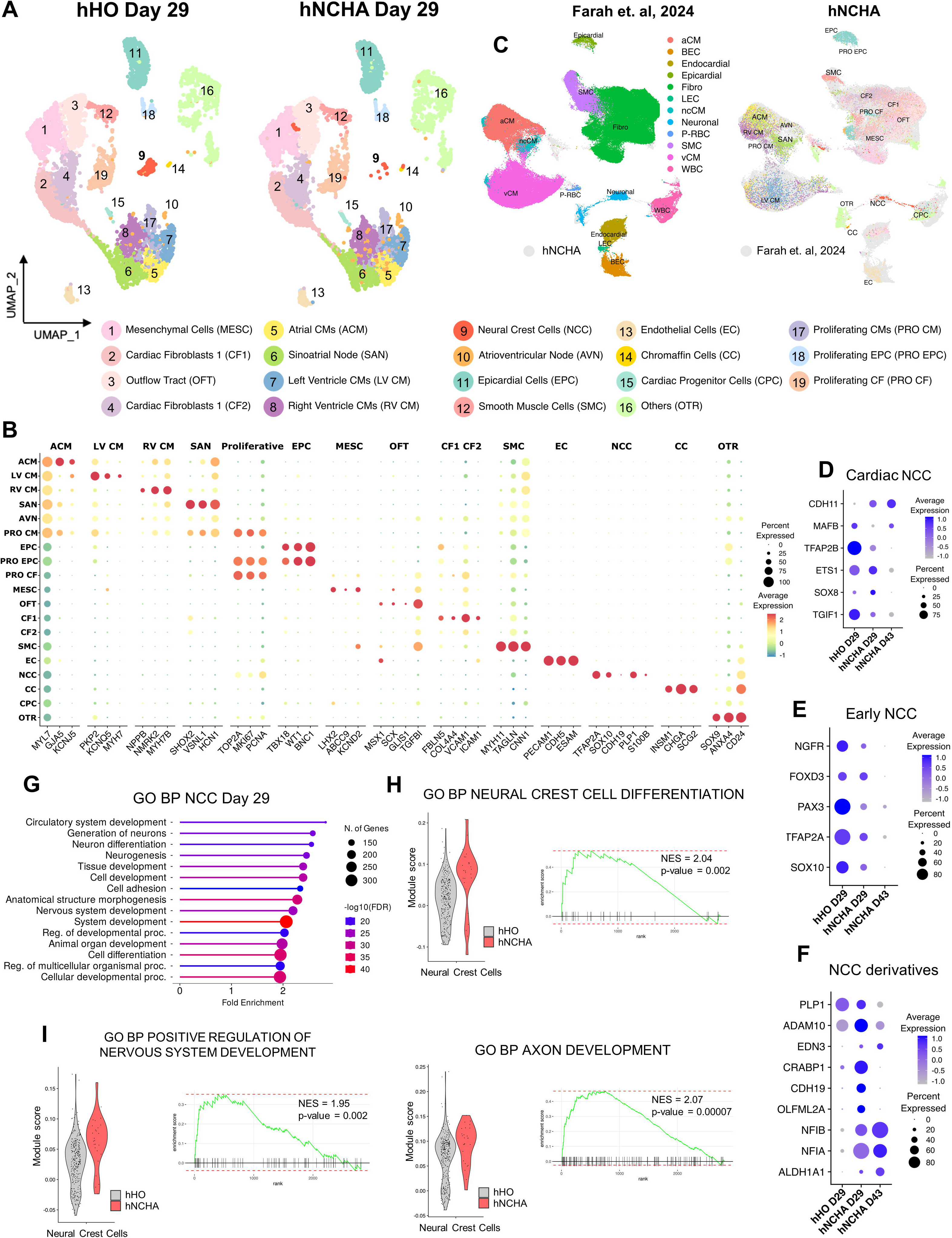
(A) Uniform manifold approximation and projection (UMAP) plots of single-cell RNA sequencing data for Day 29 hHOs and Day 29 hNCHAs. Cluster identities are in the legend below. (B) Dot plot displaying representative marker genes across cell clusters. Dot size is proportional to the percentage of cells in the cluster expressing specific genes. Color intensity indicates average expression. (C) UMAP plots of integrated hNCHA dataset (right) and 9 PCW human embryonic heart dataset (left). The color corresponds to clusters identified for hNCHAs (right UMAP) or taken from the respective publication (left UMAP). (D) (E) (F) Dot plots depicting genes specific to cardiac NCCs (D), early NCC development (E) and NCC differentiation (F) and differentially expressed in NCC cluster of Day 29 hNCHAs and Day 43 hNCHAs versus NCCs before migration. Dot size is proportional to percentage of cells expressing specific genes. Color intensity indicates average expression (p-value<0.05). (G) Lollipop plots displaying top upregulated processes in NCCs of Day 29 hNCHAs versus NCCs before migration. GO – Gene Ontology; BP – Biological Process. (H) (I) Pathway enrichment analysis in NCC cluster of Day 29 hNCHAs versus NCCs before migration. Violin plots depicting cells per sample by gene set module score. Enrichment plots of pathway-related gene sets overrepresented in NCC cluster of Day 29 hNCHAs versus NCCs before migration. NES, enrichment score normalized to mean enrichment of random samples of the same size (value represents the enrichment score after normalization).

Neural crest cells are specified based on the region of the neural tube from which they originate^60^. Cardiac NCCs migrate from the neural tube adjacent to rhombomeres 6, 7, and 8 into the forming pharyngeal arches 3, 4, and 6, where they acquire various cell fates in the developing heart^32^. Previous studies have identified unique genetic signatures specific to cardiac NCCs^61,62^. Genes within a regulatory subcircuit for cardiac NCC specification were found expressed in iPSC-derived mCherry-NCCs, with more expression of *TGIF1* and *ETS1* before migration and *SOX8* expression in migrating NCCs, along with other genes relevant to cardiac NCC development (*MAFB*, *TFAP2B*, *CDH11*) **(Fig. 2D)**. Expression of bona fide NCC markers, such as *SOX10*, *TFAP2A*, *PAX3*, *FOXD3*, and *NGFR*, which are abundant in NCCs prior to migration, was progressively decreasing in NCCs from hNCHAs, clearly demonstrating that NCCs lose their undifferentiated features during migration **(Fig. 2E)**. To determine the primary directions of NCC specification upon migration throughout hNCHAs and identify key cardiac structures formed with NCC contributions, we performed differential gene expression analysis between NCCs from hNCHAs and NCCs before integration, followed by an analysis of molecular pathways enriched in NCCs from assembloids. **Figure 2F** shows a significant upregulation of genes known as key regulators of NCC migration and the early formation of NCC derivatives. These include crucial molecules in retinoic acid (RA) signaling *ALDH1A1* and *CRABP1*^63^, as RA gradients are essential for NCC migration and contribute to the regional specification of NCC derivatives^64^. Also upregulated are the adhesion protein *CDH19*, the ECM modulator *OLFML2A*^65^, transcription factors *NFIA* and *NFIB*, which guide NCCs along specific migratory pathways^66^, *PLP1*, which is involved in the Schwann cell specification program^67^, and the signaling molecule *EDN3*^68,69^.

Pathway enrichment analysis of the NCC cluster from Day 29 hNCHAs revealed that upregulated transcripts were primarily related to circulatory system development processes, cell adhesion, and positive regulation of nervous system development, including generation of neurons, neuron differentiation and axon development (**Fig. 2G, I, J; Supp.** Fig. 2D). Enrichment of these pathways in migrating NCCs is entirely consistent with the main developmental paths previously described for cardiac NCCs^63^. This analysis also confirmed the activity of processes associated with NCC development and differentiation **(Fig. 2H; Supp.** Fig. 2E). The NCC cluster from Day 43 hNCHAs similarly demonstrated predominant activity of neuronal developmental pathways **(Supp. Fig. 2B)**, with these processes showing even more significant upregulation in Day 43 NCCs compared to Day 29 NCCs **(Supp. Fig. 2C)**. The prominent neuronal differentiation of NCCs explains the lower cell count in the NCC cluster within the Day 43 hNCHA dataset. The rarity of intrinsic cardiac neurons, along with their soma size and neuronal projections, poses challenges for scRNA-seq sample preparation, particularly during the filtration step **(Supp. Fig. 2A)**^26,53,63^.

Next, we compared hHOs and hNCHAs by the number of cells in each cluster (**Fig. 3A; Supp.** Fig. 3A). Both Day 29 and Day 43 hNCHAs demonstrated a remarkable increase in one of the cardiomyocyte populations that were defined as atrioventricular node cells in the analysis detailly described below. Notably, no difference was observed in the cell number of this cluster between Day 29 and Day 43 hHOs, suggesting a significant effect of NCC presence **(Supp. Fig. 3B)**. A decrease in the EC cluster was observed in both hHOs and hNCHAs on Day 43, which is expected due to the absence of blood flow, a critical factor for EC viability **(Supp. Fig. 3B; C)**.

**Figure 3.**
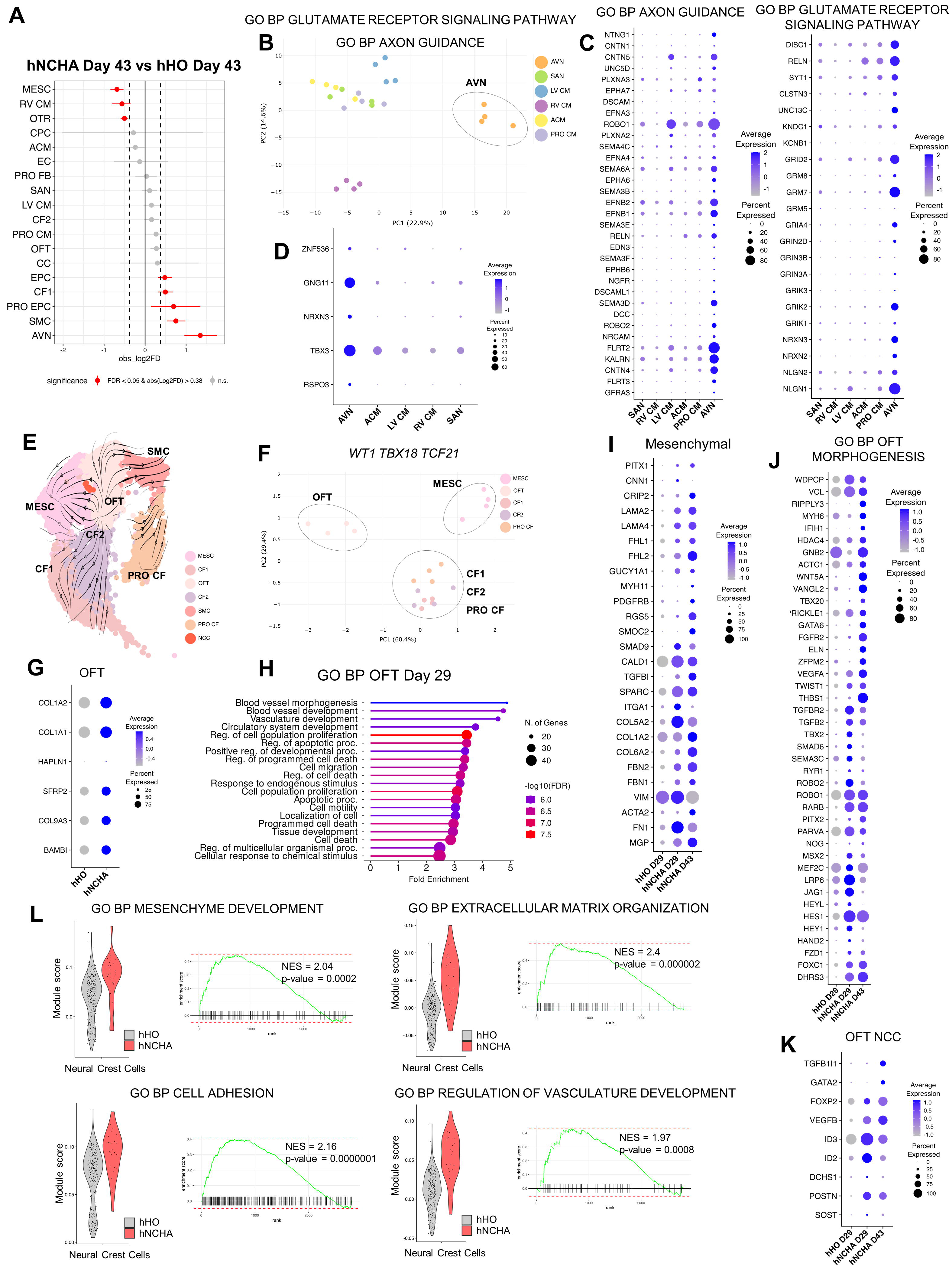
(A) Relative differences in cell proportion for each cluster in Day 43 hNCHAs versus Day 43 hHOs. Clusters colored red have a false discovery rate <0.05 and mean |log2 fold enrichment| > 0.38 (permutation test; n = 10,000). (B) A principal component analysis (PCA) plot illustrates the distinct clustering of the AVN cardiomyocyte population based on the expression of genes linked to axon guidance (GO:0007411; GO:1902667) and the glutamatergic receptor signaling pathway (GO:0007215; GO:0035249). (C) Dot plot depicting genes associated with axon guidance (according to GO:0007411; GO:1902667) and glutamatergic receptor signaling pathway (according to GO:0007215; GO:0035249) and differentially expressed in AVN cluster versus all other cardiomyocyte clusters. Dot size is proportional to percentage of cells expressing specific genes. Color intensity indicates average expression (p-value<0.05). (D) Dot plot depicting genes specific to atrioventricular node and differentially expressed in AVN cluster versus all other cardiomyocyte clusters. Dot size is proportional to percentage of cells expressing specific genes. Color intensity indicates average expression (p-value<0.05). (E) RNA velocity analysis reveals a specification of mesenchymal populations. The direction of the arrow reflects the direction of cellular state changes. (F) PCA plot illustrates the distinct clustering of the mesenchymal populations based on the expression of key epicardial genes *WT1*, *TCF21* and *TBX18*. (G) Dot plots depicting differently upregulated OFT genes in OFT cluster of hNCHAs versus hHOs. Dot size is proportional to the percentage of cells expressing specific genes. Color intensity indicates average expression (p-value<0.05). (H) Lollipop plots displaying top upregulated processes in OFT cluster of Day 29 hNCHAs versus Day 29 hHOs. GO – Gene Ontology; BP – Biological Process. (I) Dot plot depicting genes specific to mesenchymal cell differentiation and differentially expressed in NCC cluster of Day 29 hNCHAs and Day 43 hNCHAs versus NCCs before migration. Dot size is proportional to the percentage of cells expressing specific genes. Color intensity indicates average expression (p-value<0.05). (J) Dot plot depicting genes associated with OFT morphogenesis (according to GO:0003151) and differentially expressed in NCC cluster of Day 29 hNCHAs and Day 43 hNCHAs versus NCCs before migration. Dot size is proportional to the percentage of cells expressing specific genes. Color intensity indicates average expression (p-value<0.05). (K) Dot plot depicting additional genes associated with OFT development and differentially expressed in NCC cluster of Day 29 hNCHAs and Day 43 hNCHAs versus NCCs before migration. Dot size is proportional to the percentage of cells expressing specific genes. Color intensity indicates average expression (p-value<0.05). (L) Pathway enrichment analysis in NCC cluster of Day 29 hNCHAs versus NCCs before migration. Violin plots depicting cells per sample by gene set module score. Enrichment plots of pathway-related gene sets overrepresented in NCC cluster of Day 29 hNCHAs versus NCCs before migration. NES, enrichment score normalized to mean enrichment of random samples of the same size (value represents the enrichment score after normalization).

To characterize the cardiomyocyte cluster, whose cell number increased in hNCHAs due to the presence of NCCs, we analyzed the DEGs that distinguish this cluster from other cardiomyocyte populations. An abundant number of genes related to the glutamate receptor signaling pathway (GO:0007215; GO:0035249) and axon guidance (GO:0007411; GO:1902667) gene ontologies were significantly highly expressed in this cluster, which was distinctly segregated from all other cardiomyocytes in a principal component analysis (PCA) based on gene sets from these ontologies. **(Fig. 3B, C; Supp.** Fig. 3D, E). The endogenous glutamatergic transmitter system expressed in the human myocardium has been reported to play a role in the physiology and pathophysiology of vital cardiac functions^70,71^. Finally, the upregulated expression of genes specific to atrioventricular node (AVN) pacemaker cells in this cluster strongly supports the conclusion that this cardiomyocyte cluster possesses AVN identity **(Fig. 3D)**. Thus, AVN pacemaker cardiomyocytes express axon guidance genes and help to organize NCC migratory pathways and growth of NCC-generated neuronal projections. NCCs, in turn, promote an increase in the number of AVN cells and likely interact with the AVN through glutamatergic signaling, as NCCs also upregulated expression of core components of the glutamatergic system **(Supp. Fig. 3F)**.

Next, we aimed to identify the major developmental trajectories of mesenchymal cells in hHOs and hNCHAs and the potential contributions of NCCs to various populations. RNA velocity analysis revealed two primary directions of cell specification **(Fig. 3E)**, which were further supported by population subdivision based on PCA using the expression of key epicardial markers *WT1*, *TBX18*, and *TCF21* **(Fig. 3F)**. Populations with high expression of these markers were classified as epicardial-derived cells, known to be a source of cardiac fibroblasts^72–76^. Proliferating cardiac fibroblasts (PRO CF cluster) were identified based on high expression of proliferative markers such as *MKI67*, *TOP2A*, and *PCNA*, alongside cardiac fibroblast-specific genes like *FBLN5*, *VCAM1*, and *COL4A4* **(Fig. 2B).** The top markers distinguishing cardiac fibroblasts 1 (CF1 cluster) from cardiac fibroblasts 2 (CF2 cluster) are shown in **Supp.** Fig. 3G. Based on comparisons with the human embryonic heart dataset^52^, we hypothesize that CF1 may be related to subepicardial fibroblasts, while CF2 likely corresponds to myofibroblasts. However, further studies are required, as characterizing fibroblast subtypes is challenging due to their overlapping molecular signatures. A detailed analysis of populations with low to no expression of epicardial markers revealed several distinct cell types. These included cells with abundant expression of smooth muscle cell markers such as *MYH11*, *TAGLN* and *CNN1* (SMC cluster) **(Fig. 2B; Supplementary Table 3)**; cells with higher expression of genes involved in outflow tract and valve development, including *TGFBI*, *HAPLN1*, *BAMBI*, *PITX2*, *SCX*, *GLIS1*, *BMP4*, and *MSX1* (OFT cluster) ^62,77–79^ (GO:0003151) and a transitioning mesenchymal population (MESC cluster) characterized by exclusive expression of the pharyngeal mesoderm marker *LHX2*^80^, pericyte-specific genes *ABCC9*, *PDGFRB*, and *RGS5*, as well as outflow tract signatures including *HES1*, *SNAI2*, *KCND2*, *HOXB3*, *NOTCH1*, and *BMPR2* (GO:0003151) **(Fig. 2B; Supp.** Fig. 3H, I**; Supplementary Table 3)**.

Analysis of DEGs between hHOs and hNCHAs in the clusters described above revealed that presence of NCCs significantly upregulates the expression of critical genes in the OFT cluster of hNCHAs, including *BAMBI*, *COL9A3*, *SFRP2*, *COL1A1*, *COL1A2*, and *HAPLN1* **(Fig. 3G)**. The top 10 biological processes enriched in the OFT cluster of Day 29 hNCHAs included blood vessel development and morphogenesis, along with the upregulation of general pathways related to circulatory system development, anatomical structure morphogenesis, extracellular matrix organization, adhesion, and actin cytoskeleton remodeling in the MESC cluster of Day 43 hNCHAs **(Fig. 3H; Supp.** Fig. 3J, K).

Vascular SMC, pericytes and mesenchymal cells, contributing to OFT, cushion mesenchyme and aorticopulmonary septum constitute the largest lineages of the cardiac NCC derivatives^63^. We observed the part of mCherry-NCCs colocalized with MESC and OFT clusters in Day 29 hNCHAs **(Fig. 2A**; **Fig. 3E)**. Gene expression analysis revealed a gradual significant upregulation of numerous mesenchymal, pericyte, and SMC markers in NCCs from hNCHAs as compared to NCCs before migration, closely resembling cardiac NCC differentiation in vivo **(Fig. 3I)**. Moreover, NCCs from hNCHAs demonstrated a significant increase in the expression of genes belonging to the outflow tract morphogenesis gene ontology (GO:0003151) **(Fig. 3J)** and several other genes including *SOST*, *DCHS1*, *ID2*, and *ID3*, previously identified as markers of OFT development **(Fig. 3K)** ^81–86^. Finally, pathway analysis of DEGs confirmed the upregulation of processes related to mesenchyme development, mesenchymal cell differentiation, and extracellular matrix organization, along with general signaling cascades involved in heart development and morphogenesis, in NCCs from Day 29 hNCHAs compared to NCCs before migration. **(Fig. 3L; Supp.** Fig. 3L).

In summary, the scRNA-seq analysis demonstrated that human neural crest heart assembloids represent a complex multicellular model that effectively recapitulates cardiac NCC migration in the embryonic heart. These assembloids molecularly replicate how NCCs acquire multiple differentiation paths, contribute to outflow tract formation, specialize in neuronal direction, and interact with components of the cardiac conduction system.

### Neural Crest Cells promote parasympathetic innervation of hNCHAs

The cardiac autonomic nervous system regulates heart function through its extrinsic and intrinsic components. The intrinsic cardiac nervous system, composed of intracardiac neurons, is capable of independently modulating regional cardiac activity^87^. Cardiac innervation begins in the fifth week of human development and involves neural crest cell migration, differentiation into sympathetic and parasympathetic neurons, and the formation of cardiac ganglia and neuronal projections. Based on mCherry fluorescence, we observed NCCs forming neuronal-like projections in hNCHAs that exhibited synchronous contractions, suggesting their physical interaction with cardiomyocytes (**Supp. Video 6; Supp. Video 7; Supp. Video 8; Supp. Video 9**). Immunofluorescence analysis revealed that mCherry-NCCs develop an extensive neurofilament network, marked by Neurofilament-160 (*NEFM*), in Day 30 hNCHAs **(Fig. 4A)**, which continues to grow and expand as hNCHAs develop through Day 45 **(Fig. 4B; Supp.** Fig. 4A) and Day 60 **(Supp. Fig. 4B)**. We never detected NEFM^+^ neurofilaments in hHOs without NCCs **(Supp. Fig. 4D)**. High magnification images show abundance presence of additional neuronal specific cytoskeletal proteins Peripherin (*PRPH*) and class III b-tubulin (*TUBB3*) in mCherry-NCC-derived projections of Day 30 hNCHAs **(Fig. 4C, D; Supp.** Fig. 4E) and Day 45 hNCHAs **(Supp. Fig. 4F, G)**. Notably, NCC neuronal derivatives were organized into TUBB3^+^ ganglionated structures **(Fig. 4D, top; Suppl. Fig. 4E, bottom)**. Upregulation of neuronal markers visualized in hNCHAs was confirmed by the expression of corresponding genes using qRT-PCR **(Supp. Fig. 4C)**. At the same time, scRNA-seq showed increased expression of these genes specifically in NCCs **(Fig. 4E)**. Further scRNA-seq analysis demonstrated significant upregulation of numerous genes associated with neuronal development in NCCs from hNCHAs as compared to NCCs before integration **(Fig. 4F)**^88–94^.

**Figure 4.**
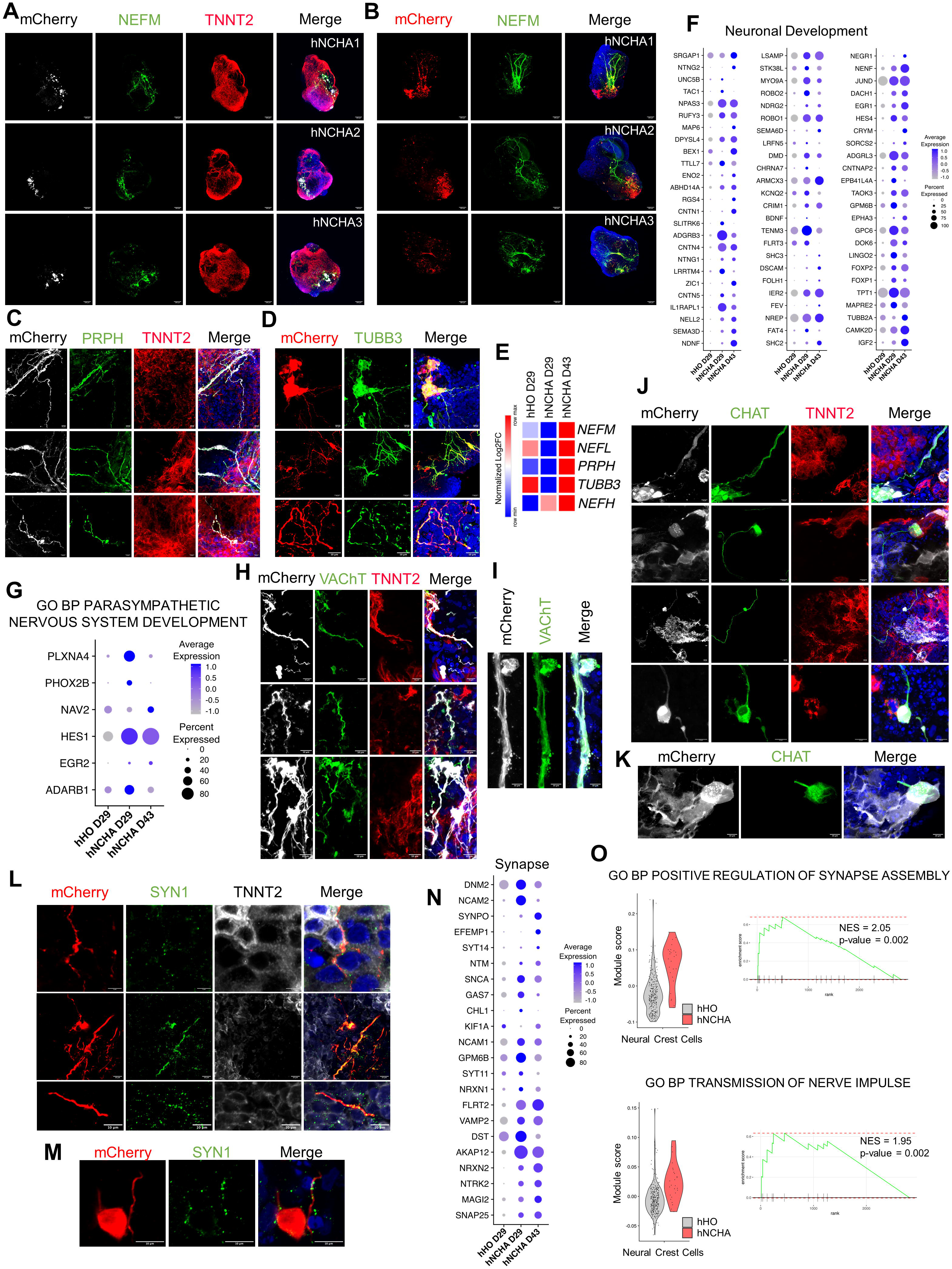
(A) Representative immunofluorescence images of three Day 30 hNCHAs displaying mCherry-NCCs (grey), neuronal marker Neurofilament-160 (*NEFM*, green), cardiomyocyte marker TNNT2 (red) and the nuclear marker DAPI (blue); Scale bar = 100 mm. (B) Representative immunofluorescence images of three Day 45 hNCHAs displaying mCherry-NCCs (grey), neuronal marker Neurofilament-160 (*NEFM*, green), cardiomyocyte marker TNNT2 (red) and the nuclear marker DAPI (blue); Scale bar = 100 mm. (C) Representative immunofluorescence images of three Day 30 hNCHAs displaying mCherry-NCCs (grey), neuronal marker Peripherin (*PRPH*, green), cardiomyocyte marker TNNT2 (red) and the nuclear marker DAPI (blue); Scale bar = 10 mm. (D) Representative immunofluorescence images of three Day 30 hNCHAs displaying mCherry-NCCs (red), neuronal marker class III b-tubulin (TUBB3, green) and the nuclear marker DAPI (blue); Scale bar = 100 mm. (E) Heatmap depicting normalized log2 fold change for neuronal filament genes in NCC cluster of Day 29 hNCHAs and Day 43 hNCHAs versus NCCs before migration. Red correlates to maximum relative expression and blue correlates to minimum relative expression (p-value<0.05). (F) Dot plot depicting genes involved in neuronal development and differentially expressed in NCC cluster of Day 29 hNCHAs and Day 43 hNCHAs versus NCCs before migration. Dot size is proportional to the percentage of cells expressing specific genes. Color intensity indicates average expression (p-value<0.05). (G) Dot plot depicting genes associated with parasympathetic nervous system development according to GO:0048486 and differentially expressed in NCC cluster of Day 29 hNCHAs and Day 43 hNCHAs versus NCCs before migration. Dot size is proportional to the percentage of cells expressing specific genes. Color intensity indicates average expression (p-value<0.05). (H) Representative immunofluorescence images of three Day 30 hNCHAs displaying mCherry-NCCs (grey), vascular acetylcholine transporter (VAChT, *SLC18A3*, green), cardiomyocyte marker TNNT2 (red) and the nuclear marker DAPI (blue); Scale bar = 10 mm. (I) High magnification image of Day 30 hNCHAs displaying mCherry-NCCs (grey), vascular acetylcholine transporter (VAChT, *SLC18A3*, green) and the nuclear marker DAPI (blue); Scale bar = 10 mm. (J) (K) Representative high magnification immunofluorescence images of Day 60 hNCHAs, Day 70 hNCHAs and Day 90 hNCHAs displaying mCherry-NCCs (grey), parasympathetic neuronal marker choline acetyltransferase (*CHAT*, green) and the nuclear marker DAPI (blue); Scale bar = 10 mm. (L) (M) Representative high magnification immunofluorescence images of Day 30 hNCHAs displaying mCherry-NCCs (grey), Synapsin I (*SYN1*, green) and the nuclear marker DAPI (blue); Scale bar = 10 mm and 5 mm (top). (N) Dot plot depicting genes involved in synapse assembly and differentially expressed in NCC cluster of Day 29 hNCHAs and Day 43 hNCHAs versus NCCs before migration. Dot size is proportional to the percentage of cells expressing specific genes. Color intensity indicates average expression (p-value<0.05). (O) Pathway enrichment analysis in NCC cluster of Day 29 hNCHAs versus NCCs before migration. Violin plots depicting cells per sample by gene set module score. Enrichment plots of pathway-related gene sets overrepresented in NCC cluster of Day 29 hNCHAs versus NCCs before migration. NES, enrichment score normalized to mean enrichment of random samples of the same size (value represents the enrichment score after normalization).

Sympathetic neurons arise from trunk NCCs, while parasympathetic innervation originates from cardiac NCCs, becoming functional before the differentiation of sympathetic neurons^95–97^. We analyzed the curated list of genes associated with parasympathetic nervous system development (GO:0048486) and found that NCCs from hNCHAs significantly upregulated the expression of several genes compared to NCCs before integration **(Fig. 4G)**. Paired-like homeobox 2b (*PHOX2B*) plays a key role in the initial differentiation of neuronal precursors to both sympathetic and parasympathetic neurons^98^. qRT-PCR confirmed upregulation of *PHOX2B* and the primary early parasympathetic marker *SLC18A3* in Day 30 hNCHAs compared to hHOs without NCCs **(Supp. Fig. 4I)**. Immunofluorescence assessment demonstrated that mCherry-NCCs differentiate into parasympathetic cholinergic neurons expressing vesicular acetylcholine transporter VAChT (*SLC18A3*) in Day 30 hNCHAs **(Fig. 4H, I)**. The key parasympathetic marker choline acetyltransferase (*CHAT*) was first detected in Day 45 hNCHAs **(Supp. Fig. 4J)**, while fully developed CHAT^+^ parasympathetic neurons forming clusters of neuronal bodies were observed in close proximity to cardiomyocytes in Day 60, Day 70, and Day 90 hNCHAs **(Fig. 4J, K)**. Weak expression of the sympathetic marker tyrosine hydroxylase (*TH*) was observed in both hNCHAs and hHOs without NCCs. Notably, TH was not localized in mCherry^+^ NCCs, hinting at the potential presence of non-neuronal intrinsic cardiac adrenergic cells known to produce TH during heart development^99,100^ **(Supp. Fig. 4H)**. NCC-derived neuronal projections extensively interlace with cardiomyocytes. To visualize potential neurocardiac junctions and synaptic varicosities, we stained Day 30 hNCHAs for the presynaptic marker Synapsin I (*SYN1*), using TNNT2 as a cardiomyocyte marker. Synapsin I was localized on the membranes of mCherry^+^ neurites at contact sites with cardiomyocytes suggesting possible neurocardiac communication **(Fig. 4L, M)**. Additionally, we investigated the expression levels of various synapse-related genes and presynaptic cell adhesion molecules (CAMs) in NCCs from hNCHAs compared to NCCs before integration and discovered significant upregulation of critical CAMs, including *NCAM1/2*, *NRXN1/2*, *FLRT2*, *CHL1*, and *NTM*; key components of the SNARE complex, such as *SNAP25* and *VAMP2*^101,102^ alongside scaffold and cytoskeletal proteins *MAGI2*, *AKAP12*, *SYT11/14*, *KIF1A*, *SNCA*, *SYNPO*, and *DNM2*, which are involved in maintaining synaptic structure, plasticity, vesicle trafficking, fusion, and neurotransmitter release **(Fig. 4N)**^96,103–107^. These data are highlighted by GO analysis, revealing terms associated with synapse assembly and nerve impulse transmission in NCCs from hNCHAs **(Fig. 4O)**. We conclude that neural crest heart assembloids develop mature NCC-derived parasympathetic neurons that establish synaptic connections with cardiomyocytes.

### NCC-derived parasympathetic neurons form functional communication with the cardiac conduction system and contribute to electrophysiological cardiomyocyte maturation

Components of the cardiac conduction system in the mammalian heart are largely innervated by NCC-derived post-ganglionic fibers^33,96,108^. Immunofluorescence analysis of hNCHAs for the sinoatrial node (SAN)-specific marker VSNL1 demonstrated mCherry^+^ NCC localization in close proximity to VSNL1^+^ cardiomyocytes in Day 30 hNCHAs **(Supp. Fig. 5A)**. By Day 45, the NCC-derived neurite network extended toward VSNL1^+^ SAN cardiomyocytes, extensively intertwining with pacemaker cells **(Fig 5A; Supp.** Fig. 5B). High magnification images show individual conductance cells of Day 45 hNCHAs physically interacting with neuronal fibers **(Fig. 5B; Supp.** Fig. 5C). Expression of crucial chemoattractant - nerve growth factor (*NGF*) was detected at varying levels across all cardiomyocyte populations, with notably higher expression in atrioventricular node cells **(Fig. 5C)**. Furthermore, we observed that SAN cells from Day 43 hNCHAs exhibited significantly higher expression of axon guidance genes compared to SAN from hHOs without NCCs. These genes included semaphorins and slit genes (*SEMA3A*, *SEMA3C*, *SLIT2*, *SLIT3*), contactins (*CNTN4*, *CNTN5*), and neuronal CAMs (*NRXN1*, *NRXN3*, *PTK2*, *NCAM1*) **(Fig. 5D)**. Notably, the expression of corresponding receptors (*NGFR*, *PLXNA2*, *PLXNA4*, *ROBO1*, *ROBO2*) by migrating NCCs **(Fig. 4F, G)** confirms the presence of a finely tuned molecular machinery in hNCHAs that regulates proper NCC migration and neurocardiac development via components of the cardiac conduction system. Consistently, pathway analysis of the SAN cluster from hNCHAs revealed enrichment in processes related to neuron projection development, synapse assembly, and postsynaptic membrane structure, further supporting the establishment of functional cross-talk between developing neurons and conductive cardiomyocytes **(Fig. 5E)**.

**Figure 5.**
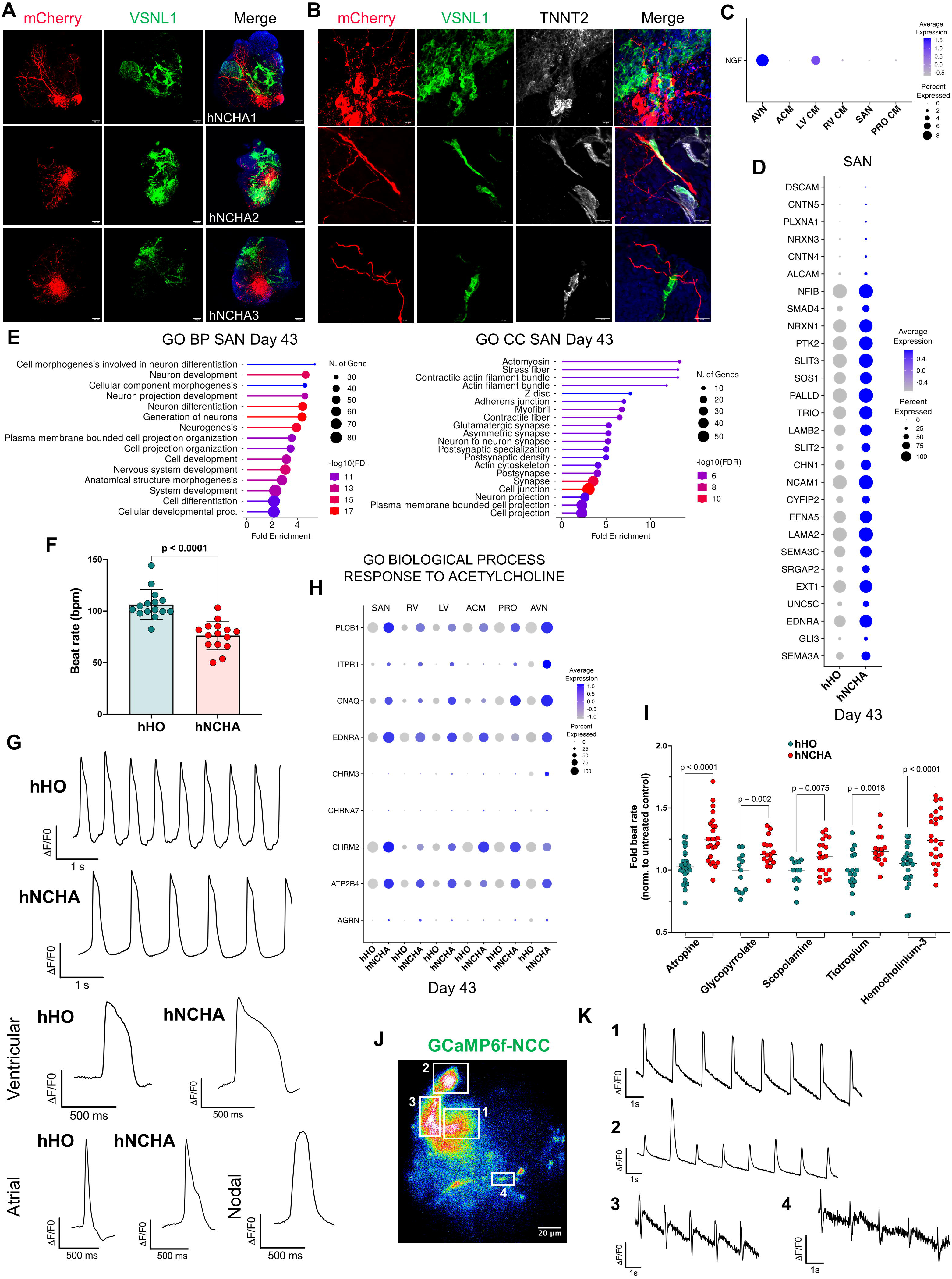
(A) Representative immunofluorescence images of three Day 45 hNCHAs displaying mCherry-NCCs (red), sinoatrial node marker *VSNL1* (green) and the nuclear marker DAPI (blue); Scale bar = 100 mm. (B) Representative high magnification immunofluorescence images of three Day 45 hNCHAs displaying mCherry-NCCs (red), sinoatrial node marker *VSNL1* (green), cardiomyocyte marker TNNT2 (grey) and the nuclear marker DAPI (blue); Scale bar = 20 mm. (C) Dot plot depicting expression on *NGF* in cardiomyocyte populations of hNCHAs. Dot size is proportional to the percentage of cells expressing specific genes. Color intensity indicates average expression. (D) Dot plot depicting axon guidance genes differentially expressed in SAN cluster of Day 43 hNCHAs versus Day 43 hHOs. Dot size is proportional to the percentage of cells expressing specific genes. Color intensity indicates average expression (p-value<0.05). (E) Lollipop plots displaying top upregulated processes in sinoatrial node cardiomyocytes of Day 43 hNCHAs versus Day 43 hHOs. GO – Gene Ontology; BP – Biological Process; CC – Cellular Component. (F) Quantification of beats per minute in Day 45 hHOs and Day 45 hNCHAs; n = 15. Values = mean ± SD, unpaired t test. (G) Representative FluoVolt traces of Day 45 hHOs and Day 45 hNCHAs depicting ventricular-, atrial-, and nodal-like potentials. Traces represent data from an individual cardiomyocyte within hHOs and hNCHAs; n = 16. (H) Dot plot depicting genes related to response to acetylcholine (according to GO:1905144) and differentially expressed in cardiomyocyte clusters of Day 43 hNCHAs versus same clusters in Day 43 hHOs. Dot size is proportional to the percentage of cells expressing specific genes. Color intensity indicates average expression (p-value<0.05). (I) Quantification of beats per minute of hHOs and hNCHAs after 1 hour treatment with 1 mM Atropine, 1 mM Glycopyrrolate, 1 mM Scopolamine, 1 mM Tiotropium, 10 mM Hemicolinium-3; n^3^ 25 (Atropine and Hemicholinnium-3) and n^3^ 15 (Glycopyrrolate, Scopolamine, Tiotropium). Beats per minute are normalized to the individual basal beating rate (before treatment). Values = mean and all individual data points; unpaired t test. (J) Live imaging of calcium transients in GCaMP6f-NCC derivatives depicting neural activity in Day 60 hNCHAs. Snapshot from a 40-sec time-lapse video is shown (see Supplementary Video 10). White frames point to fragments whose pixel intensity was measured. Numbers in snapshot correspond to numbered traces in the graphs (K). (K) The graphs representing calcium traces in hNCHAs with GCaMP6f-NCCs.

Next, we aimed to characterize the functional properties of hNCHAs. Heart assembloids with NCCs exhibited a reduced beating rate compared to hHOs without NCCs, displaying a decreasing trend in Day 30 hNCHAs that became significantly lower by Day 45 **(Fig. 5F; Supp.** Fig. 5F). An assessment of membrane potential changes in individual cardiomyocytes from Day 45 hNCHAs and hHOs, using voltage-sensitive dye (FluoVolt) and live imaging, confirmed a lower action potential frequency in hNCHAs **(Fig. 5G)**. Relied on chamber-specific action potential shapes, we found that ventricular and atrial cardiomyocytes in hNCHAs exhibited longer action potential durations compared to cardiomyocytes in hHOs, indicating a clear sign of cardiac maturation **(Fig. 5G)**. Furthermore, the presence of nodal-like cells in hNCHAs was confirmed **(Fig. 5G)**. Gene Ontology analysis of the differentially upregulated genes in ACM and VCM from Day 43 hNCHAs reflected enrichment of processes associated with cardiac maturation and affecting critical cardiac cellular components **(Supp. Fig. 5D, E)**. The upregulation of pathways related to neuronal development and synaptic structures further proved the interaction between NCC neuronal derivatives and chamber-specific cardiomyocytes. **(Supp. Fig. 5D, E)**.

Since hNCHAs contained VAChT^+^ and CHAT^+^ parasympathetic neurons and exhibited lower beating rates, we reasoned that the neurotransmitter acetylcholine (ACh) might have an impact on cardiac tissue function. Indeed, cardiomyocyte populations in Day 43 hNCHAs exhibited significantly upregulated expression of genes involved in signaling responses to acetylcholine (ACh) (GO:1905144), including the cardiac muscarinic cholinergic receptor M2 (*CHRM*2), G-protein (*GNAQ*), effector protein phospholipase C-β1 (*PLCB1*), plasma membrane calcium-transporting ATPase 4 (*ATP2B4*), and the proteoglycan agrin (*AGRN*), which is known to be involved in ACh receptor aggregation and neuromuscular junction development during embryogenesis. To assess whether the decrease in beating rate observed in hNCHAs was a result of ACh released by parasympathetic neurons, we used pharmacological agents to block muscarinic receptors and inhibit choline reuptake by its transporter, thereby suppressing the effect of ACh (1 µM Atropine, 1 µM Glycopyrrolate, 1 µM Scopolamine, 1 µM Tiotropium, and 10 µM Hemicholinium-3). hNCHAs exhibited an increase in beating rate in response to drug treatment, suggesting the effective blockade of parasympathetic neurocardiac junctions. In contrast, hHOs without NCCs showed little to no reaction to the same treatment **(Fig. 5I)**. Overall, these data provide experimental evidence of the close communication between NCC-derived parasympathetic neurons and the myocardium via the acetylcholine neurotransmitter, highlighting their significant impact on cardiac function.

We then validated the functionality of NCC-derived neurons in hNCHAs using live imaging with the Ca²⁺ sensor GCaMP6f^109^. To achieve this, we generated NCCs from an iPSC line expressing GCaMP6f and integrated them into hHOs lacking a reporter. Treatment of NCC cultures with 5 uM Ionomycin induced spontaneous calcium activity, indicating the high responsiveness of this cellular system **(Supp. Fig 5G)**. Live imaging of Day 60 heart assembloids with GCaMP6f-NCCs revealed periodical rapid calcium transients in GCaMP6f-NCC derivatives, providing evidence of functional nerve impulses generated by developing parasympathetic neurons **(Fig. 5J, K; Supplementary Video 10)**.

### Neural Crest Cells differentiate into glial cells, supporting developing parasympathetic innervation and contributing to outflow tract development

Glial cells are developmentally and functionally essential for the nervous system, including the intracardiac nervous system. While the differentiation of glial cell populations during heart development remains poorly understood, previous studies have established that most peripheral glial populations are neural crest-derived, and cardiac glia is generated by cardiac NCCs^110,111^. We stained hNCHAs for the primary glial marker S100 calcium-binding protein B (*S100B*) and observed that a subset of mCherry^+^ NCCs differentiated into glial cells of a spindle shape with thin processes at each end, which were colocalized with mCherry^+^ projections interlacing cardiomyocytes **(Fig. 6A; Supp.** Fig. 6A). High-magnification imaging clearly showed S100B^+^ cells enveloping mCherry^+^ neuronal fibers **(Fig. 6B; Supp.** Fig. 6B). Single-cell RNA-seq data and gene expression analysis by qRT-PCR provided additional evidence for the presence of NCC-derived glial cells in assembloids. Glial-associated genes were gradually upregulated in NCCs of Day 29 and Day 43 hNCHAs. Pathway analysis of differentially upregulated genes revealed enrichment in processes related to gliogenesis and glial cell differentiation **(Fig. 6C, D; Supp.** Fig. 6C). To demonstrate that differentiating mCherry^+^ glial cells are part of the developing cardiac parasympathetic innervation and provide support for growing neuronal fibers, hNCHAs were co-stained with the glial marker S100B and the neuronal markers TUBB3 and PRPH. This revealed a close association between S100B-positive glial cells and neurites labeled with neuronal markers, consistent with the characteristics of neuroglial cells **(Fig. 6E, F)**. Both glial cells and neuronal fibers expressed mCherry, confirming their NCC origin and highlighting the multipotency of NCCs migrating within hNCHAs.

**Figure 6.**
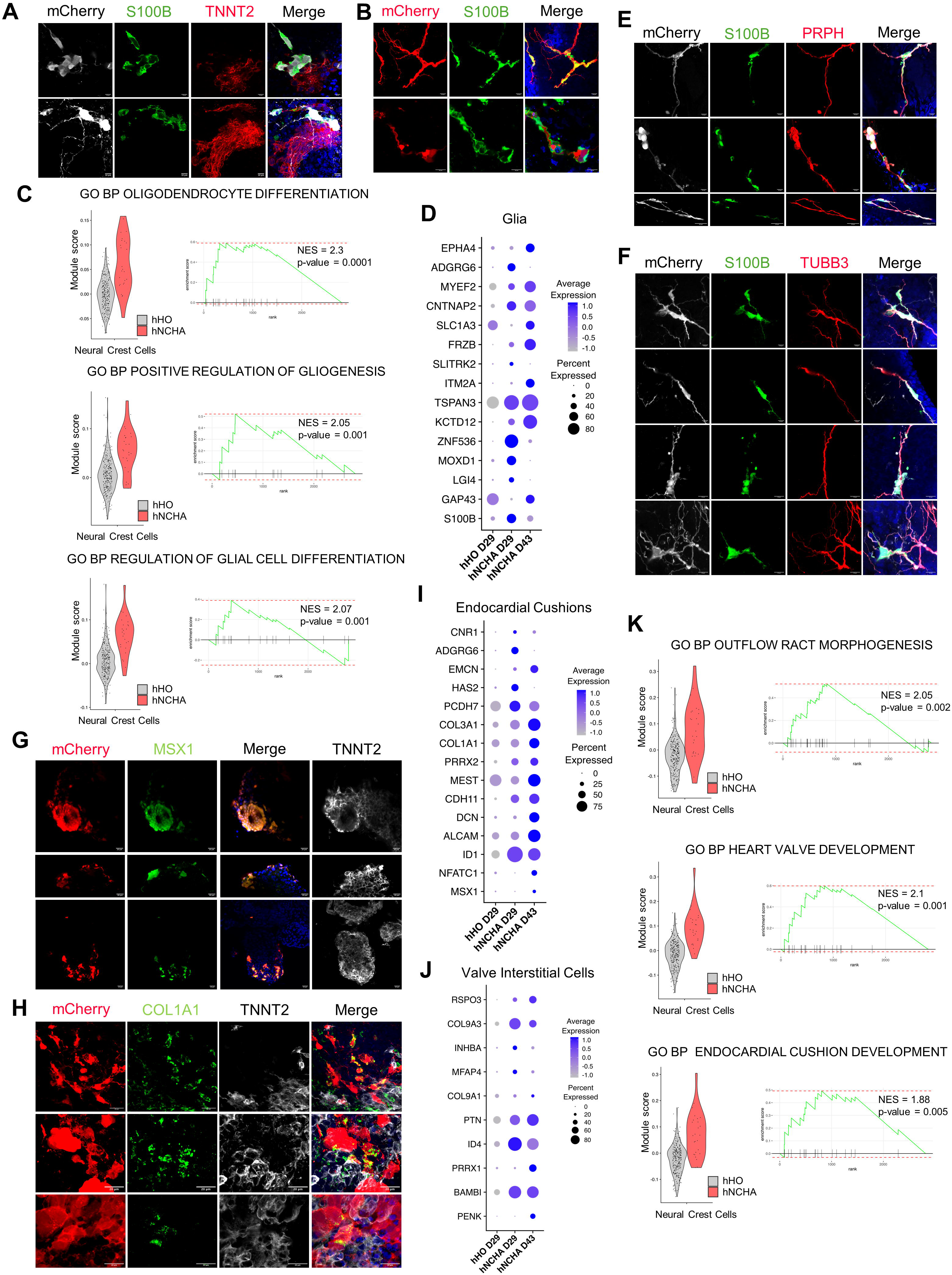
(A) Representative immunofluorescence images of Day 30 hNCHAs displaying mCherry-NCCs (grey), glial marker (*S100B*, green), cardiomyocyte marker TNNT2 (red) and the nuclear marker DAPI (blue); Scale bar = 10 mm. (B) Representative high magnification immunofluorescence images of Day 30 hNCHAs displaying mCherry-NCCs (red), glial marker (*S100B*, green) and the nuclear marker DAPI (blue); Scale bar = 10 mm. (C) Pathway enrichment analysis in NCC cluster of Day 29 hNCHAs versus NCCs before migration. Violin plots depicting cells per sample by gene set module score. Enrichment plots of pathway-related gene sets overrepresented in NCC cluster of Day 29 hNCHAs versus NCCs before migration. NES, enrichment score normalized to mean enrichment of random samples of the same size (value represents the enrichment score after normalization). (D) Dot plot depicting genes involved in glial cells development and differentially expressed in NCC cluster of Day 29 hNCHAs and Day 43 hNCHAs versus NCCs before migration. Dot size is proportional to the percentage of cells expressing specific genes. Color intensity indicates average expression (p-value<0.05). (E) Representative immunofluorescence images of Day 30 hNCHAs displaying mCherry-NCCs (grey), glial marker (*S100B*, green), neuronal marker Peripherin (*PRPH*, red) and the nuclear marker DAPI (blue); Scale bar = 10 mm and 5 mm (bottom). (F) Representative immunofluorescence images of Day 30 hNCHAs displaying mCherry-NCCs (grey), glial marker (*S100B*, green), neuronal marker class III b-tubulin (*TUBB3*, red) and the nuclear marker DAPI (blue); Scale bar = 10 mm and 5 mm (bottom). (G) Representative immunofluorescence images of Day 30 hNCHAs displaying mCherry-NCCs (red), valve marker Msh Homeobox 1 (*MSX1*, green) and the nuclear marker DAPI (blue); Scale bar = 20 mm. (H) Representative immunofluorescence images of Day 30 hNCHAs displaying mCherry-NCCs (red), valve marker Collagen 1A1 (*COL1A1*, green) and the nuclear marker DAPI (blue); Scale bar = 20 mm. (I) Dot plot depicting genes involved in valvular cuchions development and differentially expressed in NCC cluster of Day 29 hNCHAs and Day 43 hNCHAs versus NCCs before migration. Dot size is proportional to the percentage of cells expressing specific genes. Color intensity indicates average expression (p-value<0.05). (J) Dot plot depicting genes involved in valve interstitial cells development and differentially expressed in NCC cluster of Day 29 hNCHAs and Day 43 hNCHAs versus NCCs before migration. Dot size is proportional to the percentage of cells expressing specific genes. Color intensity indicates average expression (p-value<0.05). (K) Pathway enrichment analysis in NCC cluster of Day 29 hNCHAs versus NCCs before migration. Violin plots depicting cells per sample by gene set module score. Enrichment plots of pathway-related gene sets overrepresented in NCC cluster of Day 29 hNCHAs versus NCCs before migration. NES, enrichment score normalized to mean enrichment of random samples of the same size (value represents the enrichment score after normalization).

Mesenchymal NCC derivatives migrate into the developing outflow tract (OFT), where they form the aorticopulmonary septal complex and populate the OFT cushion mesenchyme, which undergoes remodeling to form the valvular leaflets^63,112^. Given that NCCs migrating in hNCHAs exhibited upregulated expression of mesenchymal genes associated with OFT formation **(Fig. 2I, J, K)**, we next visualized mCherry-NCCs with potential valvular commitment by staining for established valve formation markers transcriptional factor Msh Homeobox 1 (MSX1) and Collagen 1A1 (*COL1A1*)^113^. Double-positive mCherry^+^/MSX1^+^ cell aggregates were identified on the periphery of hNCHAs, closely associated with cardiomyocytes **(Fig. 6G; Supp.** Fig. 6D). The extracellular matrix protein Collagen 1A1 (*COL1A1*) is abundantly expressed in the developing heart, where it plays a critical role in extracellular matrix remodeling and maintaining structural integrity, as well as serving as a primary marker for valve interstitial cells (VICs)^114–118^. Consistently, we found COL1A1 expression throughout the hNCHAs **(Supp. Fig. 6E)**. High-magnification images revealed abundant COL1A1 localization in a subset of mCherry^+^ NCCs, suggesting their potential early differentiation into VICs **(Fig. 6H; Supp.** Fig. 6F). Notably, COL1A1 was detected exclusively in large, condensed mCherry^+^ NCCs lacking neuronal projections, whereas mCherry^+^ neurites were devoid of COL1A1 expression **(Supp. Fig. 6G)**. This finding highlights the divergence of the neuronal and mesenchymal multipotent NCC fates in hNCHAs. Neural crest cells from hNCHAs demonstrated a significantly increased expression of genes previously associated with valvular cushions, including *HAS2*, *PRRX2*, *MEST*, *CDH11*, and *ID1*, along with genes more specifically linked to valve interstitial cell differentiation, such as *PENK*, *BAMBI*, *PRRX1*, and *ID4* **(Fig. 6I, J)**^52,63,79,84,86,117,119–121^. Finally, signaling pathways associated with outflow tract morphogenesis, endocardial cushion formation, and heart valve development, among others, were identified as significantly upregulated based on the differentially expressed genes in NCC cluster of Day 29 hNCHAs **(Fig. 6K; Supp.** Fig. 6H). In summary, detailed immunofluorescence and gene expression analyses revealed distinct cell lineages and transcriptomic states of cardiac NCC derivatives in human heart assembloids, aligning with in vivo and in vitro studies of NCC migration during heart development.

### Early exposure of Neural Crest Cells to Selective Serotonin Reuptake Inhibitors disrupts the proper development of NCC derivatives in hNCHAs

The creation of highly complex, stem cell-derived human organ-like 3D organoids *in vitro* has opened unprecedented opportunities to study previously inaccessible developmental processes and disease states, particularly at stages where human tissue samples cannot be obtained^27^. Managing depression during pregnancy is challenging^36^, as untreated maternal depression poses risks to maternal health, while the use of Selective Serotonin Reuptake Inhibitors (SSRIs), commonly prescribed antidepressants, have been linked to fetal heart defects^41–43^ and increased risk of spontaneous abortion^122–124^. Epidemiological studies associate prenatal SSRI exposure with an increased incidence of septal defects^125,126^, transposition of the great arteries, and outflow tract malformations^38^. However, the precise mechanisms underlying these associations remain unclear^38,127^, data are controversial^128–130^, and the potential risks may be underestimated^131,132^. Disruption of NCC migration, survival, and differentiation results in defective patterning of outflow tract and great arteries, impaired looping and lengthening of heart tube, and abnormal early myocardial function. In this context, we hypothesized that human neural crest heart assembloids could serve as a suitable model for studying the effects of prenatal SSRI exposure on neural crest cell migration and the development of their derivatives.

The effects of seven commonly prescribed SSRIs - Paroxetine, Venlafaxine, Sertraline, Citalopram, Fluoxetine, Escitalopram, and Fluvoxamine - were investigated. Neural crest cells were exposed to these SSRIs throughout differentiation, beginning on Day 0 **(Figure 7A)**. Two concentrations were selected for each drug, reflecting maternal plasma levels for the specific SSRIs. Subsequently, SSRI-treated NCCs were incorporated into untreated heart organoids to create hNCHAs for further analysis **(Figure 7A)**. First, we analyzed SSRI-exposed NCCs before integration and observed a significant dose-dependent decrease in *PHOX2B* expression, with the strongest effects for the three most used SSRIs - Paroxetine, Venlafaxine, and Sertraline **(Figure 7B)**. *PHOX2B* is a gene essential for the development of autonomic neural crest derivatives and the generation of parasympathetic cholinergic and serotonergic neurons ^98,133,134^. Further gene expression analysis of Day 30 assembloids with treated NCCs revealed significant downregulation of key markers for NCC derivatives, including neuronal markers *PHOX2B*, *NEFM*, and *PRPH*, the parasympathetic marker *SLC18A3*, and glial-specific genes *S100B* and *GAP43* **(Figure 7C)**. Functional characterization demonstrated an increased beating rate in Day 45 hNCHAs containing NCCs exposed to Paroxetine and Sertraline, compared to assembloids with untreated NCCs, suggesting a potential disruption in the development of cholinergic neurons and their acetylcholine-mediated effects on cardiac function **(Figure 7D).** Interestingly, waterborne exposure to antidepressants during early development increased the heart rate at 48 hpf of zebrafish larvae^135^. Immunostaining with Neurofilament-160 (NEFM) antibodies in Day 30 hNCHAs confirmed the gene expression findings. hNCHAs exposed to Paroxetine and Sertraline exhibited significantly less developed NEFM^+^ NCC-derived neurites **(Figure 7E, F, G)**. Interestingly, the mCherry^+^ area was slightly reduced in hNCHAs with Paroxetine-treated NCCs but not in those with Sertraline-treated NCCs **(Figure 7G, top)**. This suggests that early SSRI exposure affects NCC differentiation into neuronal derivatives rather than their migration. Consistently, the ratio of NEFM^+^ area to mCherry^+^ area was significantly lower in assembloids with NCCs exposed to Paroxetine and Sertraline **(Figure 7G, bottom)**. In contrast, Venlafaxine, a serotonin-norepinephrine reuptake inhibitor, may exert a different effect on NCCs and hNCHAs, potentially due to its distinct mechanism of action. Collectively, these findings provide the first experimental human evidence of the potential deleterious effects of antidepressant use during early pregnancy, a critical period for heart formation. Our study raises important clinical questions regarding the safety of medications during pregnancy, particularly in relation to congenital heart defects. It also lays the groundwork for further research aimed at identifying alternative drugs with similar therapeutic efficacy for depressive disorders but with reduced adverse effects on embryonic heart development. Furthermore, we demonstrated the robust functional integration of NCCs into developing heart tissue, highlighting their significant impact on heart assembloids development, as only the NCCs were exposed to SSRIs. Finally, we validated the functionality of our novel human heart assembloids containing neural crest cells, showing that they respond molecularly, morphologically, and functionally to drug treatments. These findings further support the utility of these assembloids as a model for studying drug effects during early heart development.

**Figure 7.**
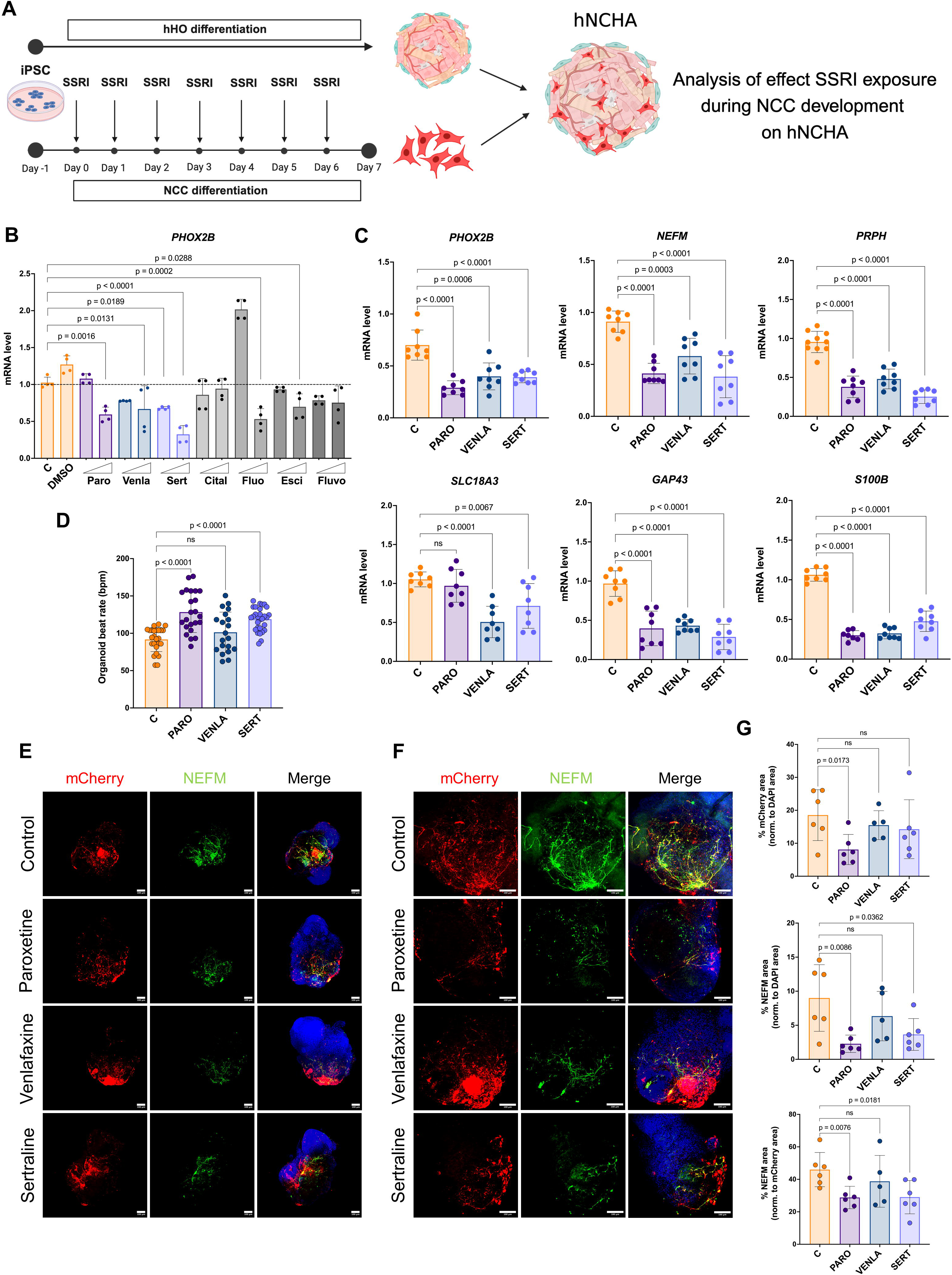
(A) A schematic representation of the experimental design illustrating the use of hNCHAs to investigate the effects of early SSRI exposure on cardiac NCCs during heart development. (B) qRT-PCR gene expression analysis of *PHOX2B* in NCCs differentiated under exposure to SSRIs; n = 4. Values = mean ± SD. The triangle represents the low and high doses of a specific SSRI. (C) qRT-PCR gene expression analysis of markers of main NCC derivatives in Day 30 hNCHAs with healthy NCCs (C) and NCCs exposed to Paroxetine (PARO), Venlafaxine (VENLA) and Sertraline (SERT) during differentiation; n = 6. Values = mean ± SD. (D) Quantification of beats per minute in Day 30 hNCHAs with healthy NCCs (C) and NCCs exposed to SSRIs during differentiation; n ≥ 20. Values = mean ± SD, unpaired t test. (E) Representative immunofluorescence images of Day 30 hNCHAs with healthy NCCs (C) and NCCs exposed to SSRIs during differentiation displaying mCherry-NCCs (red), neuronal marker *NEFM* (green) and the nuclear marker DAPI (blue); Scale bar = 100 mm. (F) Representative high magnification immunofluorescence images of Day 30 hNCHAs with healthy NCCs (C) and NCCs exposed to SSRIs during differentiation displaying mCherry-NCCs (red), neuronal marker *NEFM* (green) and the nuclear marker DAPI (blue); Scale bar = 100 mm. (G) Quantification of mCherry^+^ area and NEFM^+^ area in Day 30 hNCHAs with healthy NCCs (C) and NCCs exposed to SSRIs during differentiation measured by image analysis; n = 6. Values are presented as a percentage of DAPI^+^ area (top, middle) and mCherry^+^ are (bottom). Values = mean ± SD, unpaired t test.

## DISCUSSION

Neural crest cells (NCCs) are a highly fascinating embryonic cell population that delaminates from the neural tube and begins migrating to the developing heart in humans during the fourth week of gestation. Unlike other cell types, NCCs do not fully form cardiac structures themselves but instead contribute to and orchestrate the development of several critical structures, both morphologically and functionally. Consequently, NCCs are key players in complex tissue interactions and serve as sensitive signaling pathway modulators, making them highly vulnerable to perturbations caused by or affecting neighboring cells in these regions^112^. Despite the significant role of NCCs in cardiac development, current stem cell-derived human heart organoid models have not incorporated this indispensable cell population^21,22,136,137^. This omission restricts these models’ morphological complexity, functionality, and overall applicability in studying human heart development and disease. We leveraged the human-relevant cellular complexity of our previously published heart organoids (hHO)^15^, applied maturation strategies from our recent work^19^ and successfully integrated neural crest cells to advance the modeling of the earliest stages of human heart development in vitro. Here, we present an unprecedented human heart assembloid model containing migrating cardiac NCCs, which, during development, give rise to relevant cardiovascular derivatives of NCC origin.

We successfully differentiated NCCs exhibiting key transcriptional features and migratory behaviors characteristic of cardiac NCCs. Building on insights from animal studies about the molecular drivers of NCC migration during heart development, particularly the critical roles of *SEMA3C* and Plexin-A2 in the semaphorin-plexin signaling pathway, which regulates the migration and function of cardiac NCCs^45,47,138,139^, ensuring precise integration, we selected a developmental stage of hHOs that aligns with the signaling networks driving NCC migration in vivo. Upon integration, NCCs readily migrated upon contact with the cardiac organoids, indicating the appropriateness of the selected day for NCC addition. The NCCs exhibited a robust response to the spatiotemporal expression of developmental signals from the hHOs, closely mirroring embryonic processes observed during human heart development in vivo.

scRNA-seq data revealed the developmental trajectories of NCCs within human neural crest heart assembloids (hNCHAs). The analysis demonstrated how NCCs transition from an undifferentiated state to neuronal or mesenchymal fates—the two primary lineages of NCC derivatives in the embryonic heart. A comparison with human embryonic heart data underscored the remarkable resemblance between NCC- integrated hNCHAs and the developing human heart. NCCs in hNCHAs recapitulated the majority of transcriptional features reported for NCC development and their derivatives, as described in numerous transcriptomic studies^63,79,86,140–143^. Detailed characterization of the neuronal specification of NCCs in hNCHAs revealed their progressive alignment with the developmental trajectory of neuron formation. Initially, NCCs expressed common neuronal markers such as *NEFM*, *TUBB3*, and *PRPH*, followed by gradual parasympathetic specification, culminating in the formation of functional cholinergic neurons capable of generating nerve impulses. While sympathetic and parasympathetic neurons both derive from NCCs, they originate from distinct regions of the neural tube^144^. Notably, cardiac NCCs give rise exclusively to parasympathetic neurons of the intrinsic cardiac nervous system^97,108^. In our system, we observed the formation of functional parasympathetic neurons but not sympathetic neurons, further validating that NCCs in hNCHAs closely resemble the characteristics of cardiac NCCs in vivo. Several studies have attempted to achieve in vitro cardiac tissue innervation^145,146^. These exciting models utilized pre-differentiated neurons or neuronal progenitors for integration into cardiac tissues. In contrast, we leveraged the developmentally driven self-organization capability of multipotent NCCs, allowing them to differentiate and integrate within forming embryonic-like cardiac tissue. This approach maximized the recapitulation of early developmental events, offering a powerful model to study NCC-driven heart development and its associated processes.

Neural crest cells are a multipotent population contributing to the development of various tissues and organs, and their incorporation into organoid models has been explored in several studies. The feasibility of NCC incorporation and supporting their development within appropriate tissue contexts was previously demonstrated in human intestinal organoids, where NCCs contributed to the formation of an enteric nervous system^147^. Another study integrated NCCs derived from neuroepithelial progenitors within the neuro-mesodermal interface, further illustrating their developmental potential and interaction with surrounding tissues^148^. Both studies highlighted the critical interactions between migrating, differentiating NCCs and the tissues they inhabit. In the human neural crest heart assembloids presented here, NCCs predominantly colocalized with cardiomyocytes, forming electromechanical couplings. Cardiac function relies on the coordination between neuronal cells and cardiac pacemaker cells, a relationship established during embryonic development^96,108,149,150^. Numerous in vitro and in vivo models have demonstrated the formation of neuro-cardiac junctions, where neurons regulate cardiomyocyte function, including beating rate and cardiac maturation^151–159^. In the hNCHAs described here, neurites from neuronal NCC derivatives closely associated with sinoatrial node cardiomyocytes formed functional synaptic neuro-cardiac junctions and also increased the number of atrioventricular node cells, emphasizing the functional relevance of NCC integration in modeling early human cardiac development.

It has been shown that NCCs promote the maturation of the cardiac conduction system, while NCC ablation results in delayed maturation^160^. A detailed analysis of gene expression and activated processes revealed a highly fine-tuned and molecularly interactive system, wherein the presence of NCCs, pacemaker cardiomyocytes activate axon guidance gene programs, facilitating the development of a growing neuronal network. Simultaneously, neuronal derivatives of NCCs express both receptors and ligands involved in axon guidance pathways, thereby promoting further neuronal network expansion in close collaboration with conductive cardiomyocytes. Acetylcholine, the primary neurotransmitter of the intrinsic cardiac nervous system, acts through the muscarinic M2 receptor, which is specific to the cardiovascular system^161^. hNCHAs exhibited a decreased beating rate, indicating acetylcholine’s action on cardiac function. This was accompanied by increased expression of the M2 receptor (*CHRM2*) and other genes regulating the transcriptional response to acetylcholine. Additionally, the presence of NCC-derived neurons significantly influenced ventricular cardiomyocyte function, including an increase in action potential duration. This observation aligns with in vivo data showing depressed left ventricular function in mice with reduced cardiac acetylcholine release^162^. Conversely, chronic stimulation of parasympathetic activity has been shown to improve left ventricular function in patients with heart failure^163,164^. Notably, while NCC derivatives closely interacted with cardiomyocytes, we did not observe any direct contribution of NCCs to the cardiomyocyte population. This finding suggests possible interspecies differences in NCC specification and lineage potential^165–167^.

Consistent with previous studies^110,111,149,168^, we identified NCCs with gliagenic characteristics in hNCHAs. These cells were characterized by a neurite-enveloping morphology and the expression of the major glial marker *S100B* at both transcriptional and protein levels. While little is known about the supporting cells of the cardiac parasympathetic nervous system, existing evidence suggests that glial progenitors migrate alongside growing nerves. Neuronal-glial interactions are critical for glial cell survival, and, in turn, glial cells promote axonal growth and guidance^169^. They achieve this by releasing trophic factors, contributing to the extracellular matrix, bundling axons, and playing a pivotal role in response to cardiac tissue damage^149,170,171^. In hNCHAs, we observed the activation of transcriptional programs and processes associated with early gliagenesis, which persisted throughout development. These glial-like cells likely play an active role in regulating the growth and distribution of cholinergic neurons, mimicking their developmental functions in the embryonic heart. Interestingly, NCCs migrating to form the enteric nervous system in engineered intestinal tissue have also been shown to differentiate into both neuronal and glial structures^147^. This suggests that NCCs in hNCHAs exhibit conserved behaviors across distinct tissues, reflecting their roles in vivo.

The outflow tract (OFT) develops from the heart tube, with NCCs serving as a major extracardiac source of mesenchyme. NCCs contribute to the formation of the outflow septum and valvular cushions, playing a critical role in orchestrating proper OFT patterning^29,172^. Migrating NCCs populate the developing OFT, which is formed by the second heart field, and give rise to two condensed columns of mesenchymal cells that elongate to form the aorta and pulmonary trunk^33^. In Day 29 hNCHAs, after 12 days of NCC migration through the heart assembloids, transcriptional analysis revealed two distinct NCC populations. One of these populations colocalized with OFT mesenchymal progenitors **(Figure 2A; E)**. NCCs exhibited a strong commitment to the mesenchymal lineage, characterized by the expression of markers associated with mesenchymal specification, early OFT development, and valve interstitial cells. Gene ontology analysis highlighted the activation of processes related to aortic valve morphogenesis. Additionally, NCCs in hNCHAs expressed markers of smooth muscle cells, consistent with their known contribution to the formation of the tunica media in great arteries, including the aorta and aortic arch^173^. These findings underscore the essential role of NCCs in OFT development and the structural organization of the embryonic heart. The NCC origin of all described cell populations was confirmed by constitutive mCherry expression. The human heart assembloids with neural crest cells presented here closely replicate findings from in vivo studies, demonstrating that parasympathetic neurons, glial cells, valves, and the mesenchymal components of the OFT are all derived from the same NCC population^60,63,159^. However, the mechanisms regulating NCC specification toward neuronal or mesenchymal fates remain unclear.

The safety of selective serotonin reuptake inhibitor (SSRIs) use during pregnancy is a critical clinical concern^40^. SSRIs modulate serotonin levels, which play a key role in various aspects of cardiac development, including the patterning of cardiac progenitor cells^174^, the lengthening of the outflow tract mediated by signals from neural crest cells^175–177^, endocardial cushion formation^39,174^, and myocardial cell development^38,178,179^. Congenital heart defects associated with prenatal SSRI exposure often involve cardiac structures that rely on NCC contributions^41,126^. Characterization of heart assembloids with NCCs differentiated in the presence of SSRIs revealed early disruptions in the key transcriptional regulator *PHOX2B*, leading to impaired NCC development along neuronal and glial lineages. This disruption had functional consequences, as heart assembloids with NCCs exposed to Paroxetine and Sertraline – two widely used antidepressants – exhibited an increased beating rate, indicating abnormal differentiation and functionality of parasympathetic neurons and glial cells. These findings provide critical evidence of the potential deleterious effects of SSRI use during the first trimester, a period when the fetal heart undergoes crucial developmental processes.

In conclusion, this study developed the first molecularly and functionally relevant in vitro model that accurately recapitulates neural crest contributions to early human heart development. We demonstrated a remarkable similarity between the developmental trajectories of NCCs in heart assembloids and their counterparts in vivo. Notably, we are the first to report the formation of NCC-derived functional parasympathetic innervation, supported by mature glial cells, in a 3D model of human heart development. New NCC-derived mesenchymal populations add significant cellular complexity to the in vitro heart tissue, advancing the development of a model that closely mimics a fully formed outflow tract. Finally, the complexity and physiological relevance of the established model enabled us to replicate the abnormal development of neural crest-derived cardiac structures triggered by SSRIs in the maternal environment. These findings underscore the exceptional utility of the hNCHA-based platform for drug screening and investigating the mechanisms underlying NCC-related congenital heart diseases.

### Limitations of the study

Despite their sophisticated complexity and functionality, hNCHAs have several limitations. At their current stage of development, hNCHAs do not replicate the heart looping and complete maturation of the outflow tract. While cardiac NCCs, as a subset of vagal NCCs, contribute to parasympathetic innervation, trunk NCCs – which are essential for forming cardiac sympathetic innervation – are absent. Additionally, crucial physiological factors such as blood flow and mechanical strain are lacking. Furthermore, interactions with other cell types and tissues, such as those of endodermal origin, are necessary to provide critical signals for proper cardiac morphogenesis. Nonetheless, hNCHAs offer a promising platform to enhance complexity by integrating essential cell types. Future advancements could involve the incorporation of immune cells^27^ and endodermal progenitors^17,22,180^ into heart assembloids with neural crest cells. At this stage, hNCHAs hold significant promise for advancing our understanding of early NCC migration and specification under both healthy and disease conditions.

## METHODS

### Stem cell culture

The following human induced pluripotent stem cell (hiPSC) lines were used in this study: hiPSC-L1 (male, iPSCORE), hiPSC-L1-mCherry, and hiPSC-L1-GCaMP6f. Cells were maintained in Essential 8 Flex medium supplemented with 1% penicillin-streptomycin (Gibco) on growth factor-reduced Matrigel (Corning) in 6-well plates, incubated at 37°C with 5% CO₂. Routine analyses were performed to confirm pluripotency, karyotype integrity, and the absence of mycoplasma contamination. hiPSCs were regularly passaged using ReLeSR passaging reagent (STEMCELL Technologies) when they reached 60–80% confluency.

### Neural crest cells differentiation

hiPSC-L1-mCherry cells were differentiated into neural crest cells as previously described^44^. Briefly, hiPSC-L1- mCherry cells were seeded at a density of 15,000 cells per cm² on Matrigel-coated 6-well plates in Essential 6 medium supplemented with 1% penicillin-streptomycin and containing 2 μM ROCK inhibitor (Thiazovivin, Millipore Sigma). After 24 hours, the medium was replaced with NCC differentiation medium containing Essential 6 medium, 1% penicillin-streptomycin, N-2 supplement (Gibco), 1 μM CHIR99021, and 10 μM SB431542. The differentiation medium was changed daily. On Day 7, the cultures were detached using Accutase (Innovative Cell Technologies) and used for integration into human heart organoids.

To analyze migratory capacity, neural crest cells were seeded at a density of 10,000 cells per well in round-bottom, ultra-low attachment 96-well plates (Costar) in a volume of 100 μL. The plate was centrifuged at 100 g for 3 minutes and incubated at 37°C with 5% CO₂. After 24 hours, neural crest cells aggregates were transferred to Matrigel-coated 6-well plates using wide-bore tips at a density of 10 aggregates per well in maintenance medium consisting of DMEM/F12, N-2 supplement, and 1% penicillin-streptomycin. The plates were incubated at 37°C with 5% CO₂ and monitored daily for evidence of migrating neural crest cells.

### Human heart organoid differentiation

Self-assembling human heart organoids were generated as detailed in previous protocols^15,181^. Briefly, hiPSC- L1 cells were detached using Accutase and dissociated into a single-cell suspension. The cells were collected and centrifuged at 300 g for 5 minutes, and the pellet was resuspended in 1 mL of Essential 8 Flex medium supplemented with 2 μM ROCK inhibitor (Thiazovivin). The hiPSCs were counted using a Moxi cell counter (Orflo Technologies) and seeded at a density of 10,000 cells per well in round-bottom, ultra-low attachment 96- well plates (Costar) in a volume of 100 μL. The plate was centrifuged at 100 g for 3 minutes and incubated undisturbed at 37°C with 5% CO₂. After 24 hours, 50 μL of medium in each well was replaced with 200 μL of fresh Essential 8 Flex medium, and the plate was returned to the incubator. After another 24 hours, 166 μL of medium in each well was replaced with differentiation medium consisting of RPMI medium with B27 supplement minus insulin (Gibco) (hereafter termed RPMI/B27 minus insulin), 1% penicillin-streptomycin (Gibco), 6 μM CHIR99021, 1.875 μg/mL BMP4, and 1.5 μg/mL Activin A. The plate was incubated at 37°C with 5% CO₂. Exactly 24 hours later, 166 μL of medium in each well was replaced with fresh RPMI/B27 minus insulin. After another 24 hours, 166 μL of medium was replaced with RPMI/B27 minus insulin containing 3 μM Wnt-C59 (Selleck). After 48 hours, 166 μL of medium was replaced with fresh RPMI/B27 minus insulin. Another 48 hours later, 166 μL of medium was replaced with RPMI medium with B27 supplement with insulin (hereafter termed RPMI/B27) and 1% penicillin-streptomycin. After an additional 24 hours, 166 μL of medium in each well was replaced with RPMI/B27 containing 3 μM CHIR99021. The plate was incubated for 1 hour, followed by replacement of 166 μL of medium with fresh RPMI/B27. From Day 9 to Day 17, human heart organoids were maintained in RPMI/B27, with 166 μL of medium replaced with fresh RPMI/B27 every 48 hours.

### Generation of hNCHAs

Two days prior to integration (Day -2), NCCs were resuspended in maintenance medium composed of DMEM/F12 (Gibco), N-2 supplement, 1% penicillin-streptomycin, and 2 µM ROCK inhibitor (Thiazovivin). A total of 2,000 cells per well were seeded into a v-bottom ultra-low attachment 96-well plate at a volume of 100 µL per well. The plate was centrifuged at 100 g for 3 minutes and incubated at 37°C with 5% CO₂. After 24 hours (Day -1), 50 µL of medium was carefully removed from each well, and 100 µL of fresh maintenance medium was added, bringing the final volume to 150 µL per well. The plate was then returned to the incubator for an additional 24 hours. On Day 0, 120 µL of medium was removed from each well and replaced with 120 µL of RPMI/B27 medium. hHOs at Day 17 were transferred into the v-bottom plate containing NCCs using wide-bore pipette tips to minimize disruption of the organoids. The plate was returned to the incubator and cultured undisturbed for 48 hours. Subsequently, 120 µL of RPMI/B27 medium was refreshed every 48 hours. Starting on Day 20 (three days after integration), hNCHAs were cultured in enhanced maturation medium 2/1, as previously described^19^. After Day 30, the hNCHAs were maintained in EMM1 medium until they were ready for analysis.

### hHO and hNCHA dissociation

hHOs and hNCHAs were collected individually into separate tubes and washed with PBS. Cardiomyocyte Isolation Enzyme 2 (Thermo) was diluted at a 1:1000 ratio into Cardiomyocyte Dissociation Medium (STEMCELL Technologies), and the mixture was pre-warmed to 37°C before use. A total of 200 µL of TrypLE (Gibco) was added to each hHO/hNCHA and incubated on a thermocycler at 300 RPM and 37°C for 7 minutes. After incubation, the TrypLE was collected and transferred to RPMI/B27 medium containing 2% BSA (w/v). Next, 200 µL of the pre-warmed Cardiomyocyte Dissociation Medium with Enzyme 2 was added to the hHO/hNCHA and incubated under the same conditions (300 RPM, 37°C, for 7 minutes). This step was repeated 5–6 times or until the hHO/hNCHA was fully dissociated and no longer visible. The dissociated cells in RPMI/B27 + 2% BSA were filtered through a 50 µm filter and centrifuged at 300 g for 7 minutes. The resulting cell pellet was resuspended in a smaller volume of RPMI/B27 + 2% BSA, filtered again through a 50 µm filter, and proceeded with scRNA-seq library preparation. Day 29 and Day 43 hHOs/hNCHAs were analyzed by scRNA-seq. Control NCCs (NCCs before integration) were dissociated into a single-cell suspension using Accutase, filtered through a 50 µm filter and pooled with Day 29 hHOs sample at 10%. Subsequently, this population was isolated from Day 29 hHOs sample based on mCherry presence during bioinformatic analysis.

### Single-cell RNA sequencing

Libraries were prepared using the 10x Chromium Next GEM Single Cell 3’ Kit, v3.1 and associated components. Libraries were pooled in equimolar proportions, then loaded onto 1 lane of an Illumina NovaSeq 6000 S4 flow cell (v1.5). The sequencing was performed in a 2x150bp paired end format using 300 NovaSeq v1.5 reagent cartridge. The counts matrix was generated using 10x Cell Ranger (v7.1.1) with the Ensemble Homo sapiens GRCh38.110 genome as the reference. To maximize read mapping for the mCherry marker transgene, a 3030 bp sequence, spanning from the Kozak sequence upstream of mCherry to the SV40 early pA, was excised from the pLV[Exp]-Puro-EF1A>mCherry plasmid sequence (https://en.vectorbuilder.com/vector/VB010000-9485zvg.html). This custom sequence was incorporated into the reference genome following 10x Genomics guidelines. As a result, 30% more reads were mapped to the transgene compared to using the mCherry CDS alone. The data were then converted into Seurat (v5.0.2) objects by sample^182^. To retain control NCCs (before integration) from Day 29 hHOs sample the cells containing the mCherry transgene sequence were first isolated into a separate object. The quality filtering was performed using the following criteria: nFeature_RNA: 2000–7500, nCount_RNA: 1500–40,000, percent_mito: <15%. Subsequently, Seurat objects were merged and integrated by sample using the RPCA method within Seurat. The Louvain algorithm (Seurat-integrated) was used to generate the UMAP. For integration with data from Farah et al.^55^, SCT transform and Harmony (v0.1, Broad) were employed^183^. Throughout the workflow, auxiliary functions from the scCustomise package (doi:10.5281/zenodo.5706430) were utilized at various stages. Differential expression gene analysis and pathway enrichment analysis were performed as previously described^16^. Log₂fold change values and p-values are provided in **Supplementary Table 4**. Function AddModuleScore from Seurat was used to calculate module scores for enriched pathways. For mRNA velocity analysis, Velocyto^184^ was used to generate initial outputs, which were then converted into Scanpy AnnData objects. Secondary analyses and visualizations were performed using scvelo^185^. Permutation testing to determine the significance of proportional difference in cell populations was calculated using the tool scProportionTest^186^.

### Total RNA extraction and qRT-PCR

hHOs and hNCHAs were collected in Eppendorf tubes and washed with PBS. Total RNA was extracted using the Qiagen RNeasy Mini Kit, following the manufacturer’s instructions. RNA concentration was measured using the Qubit 4 Fluorometer with the RNA High Sensitivity Assay Kit (Thermo Fisher Scientific). cDNA was synthesized using the Qiagen QuantiTect Reverse Transcription Kit. Primers for qRT-PCR were designed using the Primer-BLAST tool. qRT-PCR reactions were performed using the Qiagen QuantiTect SYBR Green PCR Kit on a QuantStudio 5 Real-Time PCR System (Applied Biosystems). Gene expression levels were normalized to *HPRT1*, and fold change values were calculated using the 2^−ΔΔCT method. Primer sequences are provided in Supplementary Table 1.

### Immunofluorescence staining

hHOs and hNCHAs were prepared for confocal microscopy as previously described^187^. Briefly, the samples were fixed in 4% paraformaldehyde, then washed in PBS containing 20 mM glycine. Following this, the samples were incubated overnight at 4°C in a blocking and permeabilization solution on a thermal mixer. The next day, hHOs/hNCHAs were incubated with primary antibodies in antibody solution on a thermal mixer at 4°C for 24 hours. Subsequently, the samples were incubated with secondary antibodies under the same conditions for 24 hours in the dark. Finally, the hHOs/hNCHAs were mounted on glass slides using a mounting-clearing medium composed of 60% (vol/vol) glycerol and 2.5 M fructose. Antibodies are provided in Supplementary Table 2.

### Confocal microscopy and image analysis

Immunofluorescence images were acquired using a Nikon Instruments A1 Confocal Laser Microscope, and image analysis was performed with Fiji (https://imagej.net/Fiji). Several images in the merged channels do not include TNNT2 staining to improve visual clarity. To quantify mCherry+, NEFM+, and DAPI+ areas, the auto-threshold function in Fiji was applied, and the areas were measured and expressed as a percentage of the total DAPI+ area in the corresponding graphs. For colocalization analysis, Pearson’s correlation coefficient was calculated using the JaCOP colocalization plugin in Fiji.

### Live imaging for membrane potential and calcium transients analysis

Changes in membrane potential were visualized using the FluoVolt Membrane Potential Kit (Thermo). Day 45 hHOs/hNCHAs were loaded with a voltage-sensitive fluorescent probe according to the manufacturer’s instructions. Live-cell imaging videos were recorded for 20 seconds at a frame rate of 104 frames per second using a super-resolution microscope (Olympus CellVivo). NCCs were differentiated from hiPSC-L1-GCaMP6f. To rapidly elevate intracellular calcium levels, the cells were treated with 5 µM Ionomycin (STEMCELL Technologies). hHOs were generated from hiPSC-L1, and NCC-GCaMP6f cells were integrated into the hHOs to create hNCHAs. Calcium transients were observed in Day 60 hNCHAs through live imaging using Olympus CellVivo. Recordings were captured over 40 seconds at a frame rate of 104 frames per second. Membrane potential and calcium transients data were analyzed using Fiji, and fluorescence values were expressed as ΔF/F0.

### Exposure to SSRIs

NCCs were differentiated for 7 days following the described protocol. SSRIs were added daily, starting on Day 0, during each differentiation media change. The concentrations of SSRIs used were as follows: Paroxetine (0.055 µM; 0.2 µM), Venlafaxine (0.35 µM; 1.5 µM), Sertraline (0.1 µM; 0.45 µM), Citalopram (0.245 µM; 1 µM), Fluoxetine (0.3 µM; 0.9 µM), Escitalopram (0.05 µM; 0.1 µM), Fluvoxamine (0.25 µM; 1.5 µM). The higher concentrations of SSRIs were used for analyzing beating rate, gene expression in hNCHAs, and immunofluorescence.

### Analysis of acetylcholine effect

Day 45 hHOs/hNCHAs were treated with muscarinic acetylcholine receptor antagonists (1 µM Atropine, 1 µM Glycopyrrolate, 1 µM Scopolamine, and 1 µM Tiotropium) or a competitive inhibitor of the high-affinity choline transporter, 10 µM Hemicholinium 3, for 1 hour. Live videos were recorded before and after treatment to assess changes in the beating rate.

### Live imaging and beating rate analysis

Fluorescence and phase-contrast live images of hNCHAs were acquired using the CellVoyager CQ1 Benchtop High-Content Analysis System (Yokogawa). Phase-contrast live videos were recorded using an inverted Thunder Microscope (Leica) at a frame rate of at least 15 frames per second. The beat rate was calculated by counting the total number of beats observed in each 1-minute video.

### Statistical analysis

All analyses were performed using GraphPad Prism 10 software. Statistical significance was assessed using either an unpaired Student’s t-test or one-way ANOVA, depending on the experimental design. Differences were considered statistically significant at p < 0.05. Data are presented as the mean ± SD and are derived from a minimum of three independent experiments.

### Data availability

scRNA-seq data has been deposited in the National Center for Biotechnology Information Gene Expression Omnibus repository under accession number GSE280777. The organoid Bulk RNA-Sequencing dataset used in this study is available under accession code GSE153185. The organoid scRNA-seq datasets used in this study are available under accession codes GSE218582 and GSE201343. scRNA-seq dataset of human embryonic heart used in this study for integration is available from dbGAP under accession number phs002031. All data generated or analyzed in this study are provided in the published article and its supplementary information files or can be obtained from the corresponding author upon request.

## Supporting information

Supplementary video descriptions

## ACKNOWLEDGEMENTS

We thank the IQ and MSU Advanced Microscopy Cores for access to confocal microscopes and the MSU Genomics and Stem Cell Cores for sequencing and cell culture services. We also want to thank Dr. Brian Johnson and his lab for helping with live imaging. We also want to thank all members of the Aguirre Lab for their valuable comments and advice. Work in Dr. Aguirre’s laboratory was supported by startup funds from MSU, the National Institutes of Health (NIH) under award numbers K01HL135464, R01HL151505, by the American Heart Association under award numbers 19IPLOI34660342, 23IPA1053441, by the Corewell-MSU Foundation, Corewell Health, and the Alternatives Research and Development Foundation (ARDF). Work in Dr. Park’s lab was supported by startup funds from MSU and the NIH under award number R01AR083086.

## AUTHOR CONTRIBUTIONS

A.K. and A.A. designed all experiments, conceptualized all the work, assembled figures, and wrote the manuscript. A.K., A.H., H.L., S.C., B.V., A.J-F., and C.O. performed cell differentiation, cell and organoid culture experiments. A.K., H.L. and S.C. performed PCR experiments. A.K. performed confocal microscopy and image analysis. A.Ki. and S.P. integrated, processed, and analyzed scRNA-seq data. A.J., Y.L., and Z.Q. performed and analyzed live-imaging. A.K. performed all other experiments not explicitly mentioned in this section.

## REFERENCES

1. Di Cesare, M., Perel, P., Taylor, S., Kabudula, C., Bixby, H., Gaziano, T.A., McGhie, D.V., Mwangi, J., Pervan, B., Narula, J., et al. (2024). The Heart of the World. Glob Heart 19, 11. 10.5334/gh.1288.

2. Hoffman, J.I., and Kaplan, S. (2002). The incidence of congenital heart disease. J Am Coll Cardiol 39, 1890–1900. 10.1016/s0735-1097(02)01886-7.

3. National Center for Health Statistics. Multiple Cause of Death 2018–2022 on CDC WONDER Database. .

4. Costantini, D.L., Arruda, E.P., Agarwal, P., Kim, K.H., Zhu, Y., Zhu, W., Lebel, M., Cheng, C.W., Park, C.Y., Pierce, S.A., et al. (2005). The homeodomain transcription factor Irx5 establishes the mouse cardiac ventricular repolarization gradient. Cell 123, 347–358. 10.1016/j.cell.2005.08.004.

5. Xu, H., Morishima, M., Wylie, J.N., Schwartz, R.J., Bruneau, B.G., Lindsay, E.A., and Baldini, A. (2004). Tbx1 has a dual role in the morphogenesis of the cardiac outflow tract. Development 131, 3217–3227. 10.1242/dev.01174.

6. Grego-Bessa, J., Luna-Zurita, L., del Monte, G., Bolos, V., Melgar, P., Arandilla, A., Garratt, A.N., Zang, H., Mukouyama, Y.S., Chen, H., et al. (2007). Notch signaling is essential for ventricular chamber development. Dev Cell 12, 415–429. 10.1016/j.devcel.2006.12.011.

7. Timmerman, L.A., Grego-Bessa, J., Raya, A., Bertran, E., Perez-Pomares, J.M., Diez, J., Aranda, S., Palomo, S., McCormick, F., Izpisua-Belmonte, J.C., and de la Pompa, J.L. (2004). Notch promotes epithelial-mesenchymal transition during cardiac development and oncogenic transformation. Genes & development 18, 99–115. 10.1101/gad.276304.

8. Theodoris, C.V., Zhou, P., Liu, L., Zhang, Y., Nishino, T., Huang, Y., Kostina, A., Ranade, S.S., Gifford, C.A., Uspenskiy, V., et al. (2021). Network-based screen in iPSC-derived cells reveals therapeutic candidate for heart valve disease. Science 371. 10.1126/science.abd0724.

9. Wessels, A., and Sedmera, D. (2003). Developmental anatomy of the heart: a tale of mice and man. Physiol Genomics 15, 165–176. 10.1152/physiolgenomics.00033.2003.

10. Shah, A., Goerlich, C.E., Pasrija, C., Hirsch, J., Fisher, S., Odonkor, P., Strauss, E., Ayares, D., Mohiuddin, M.M., and Griffith, B.P. (2022). Anatomical Differences Between Human and Pig Hearts and Their Relevance for Cardiac Xenotransplantation Surgical Technique. JACC Case Rep 4, 1049–1052. 10.1016/j.jaccas.2022.06.011.

11. Liu, C., and Shao, N.Y. (2024). The Differences in the Developmental Stages of the Cardiomyocytes and Endothelial Cells in Human and Mouse Embryos at the Single-Cell Level. International journal of molecular sciences 25. 10.3390/ijms25063240.

12. Hamlin, R.L., and Keene, B.W. (2020). Species differences in cardiovascular physiology that affect pharmacology and toxicology. Current Opinion in Toxicology 23-24, 106-113. 10.1016/j.cotox.2020.07.004.

13. Krishnan, A., Samtani, R., Dhanantwari, P., Lee, E., Yamada, S., Shiota, K., Donofrio, M.T., Leatherbury, L., and Lo, C.W. (2014). A detailed comparison of mouse and human cardiac development. Pediatr Res 76, 500–507. 10.1038/pr.2014.128.

14. Ruiz-Orera, J., Miller, D.C., Greiner, J., Genehr, C., Grammatikaki, A., Blachut, S., Mbebi, J., Patone, G., Myronova, A., Adami, E., et al. (2024). Evolution of translational control and the emergence of genes and open reading frames in human and non-human primate hearts. Nat Cardiovasc Res 3, 1217–1235. 10.1038/s44161-024-00544-7.

15. Lewis-Israeli, Y.R., Wasserman, A.H., Gabalski, M.A., Volmert, B.D., Ming, Y., Ball, K.A., Yang, W., Zou, J., Ni, G., Pajares, N., et al. (2021). Self-assembling human heart organoids for the modeling of cardiac development and congenital heart disease. Nature communications 12, 5142. 10.1038/s41467-021-25329-5.

16. Kostina, A., Lewis-Israeli, Y.R., Abdelhamid, M., Gabalski, M.A., Kiselev, A., Volmert, B.D., Lankerd, H., Huang, A.R., Wasserman, A.H., Lydic, T., et al. (2024). ER stress and lipid imbalance drive diabetic embryonic cardiomyopathy in an organoid model of human heart development. Stem cell reports 19, 317–330. 10.1016/j.stemcr.2024.01.003.

17. Drakhlis, L., Biswanath, S., Farr, C.M., Lupanow, V., Teske, J., Ritzenhoff, K., Franke, A., Manstein, F., Bolesani, E., Kempf, H., et al. (2021). Human heart-forming organoids recapitulate early heart and foregut development. Nat Biotechnol 39, 737–746. 10.1038/s41587-021-00815-9.

18. Hofbauer, P., Jahnel, S.M., Papai, N., Giesshammer, M., Deyett, A., Schmidt, C., Penc, M., Tavernini, K., Grdseloff, N., Meledeth, C., et al. (2021). Cardioids reveal self-organizing principles of human cardiogenesis. Cell 184, 3299–3317 e3222. 10.1016/j.cell.2021.04.034.

19. Volmert, B., Kiselev, A., Juhong, A., Wang, F., Riggs, A., Kostina, A., O’Hern, C., Muniyandi, P., Wasserman, A., Huang, A., et al. (2023). A patterned human primitive heart organoid model generated by pluripotent stem cell self-organization. Nature communications 14, 8245. 10.1038/s41467-023-43999-1.

20. Gu, M., and Zorn, A.M. (2021). Follow your heart and trust your gut: Co-development of heart and gut tissue in organoids. Cell stem cell 28, 2037–2038. 10.1016/j.stem.2021.09.003.

21. Schmidt, C., Deyett, A., Ilmer, T., Haendeler, S., Torres Caballero, A., Novatchkova, M., Netzer, M.A., Ceci Ginistrelli, L., Mancheno Juncosa, E., Bhattacharya, T., et al. (2023). Multi-chamber cardioids unravel human heart development and cardiac defects. Cell 186, 5587–5605 e5527. 10.1016/j.cell.2023.10.030.

22. Silva, A.C., Matthys, O.B., Joy, D.A., Kauss, M.A., Natarajan, V., Lai, M.H., Turaga, D., Blair, A.P., Alexanian, M., Bruneau, B.G., and McDevitt, T.C. (2021). Co-emergence of cardiac and gut tissues promotes cardiomyocyte maturation within human iPSC-derived organoids. Cell stem cell 28, 2137–2152 e2136. 10.1016/j.stem.2021.11.007.

23. Ng, W.H., Varghese, B., Jia, H., and Ren, X. (2023). Alliance of Heart and Endoderm: Multilineage Organoids to Model Co-development. Circulation research 132, 511–518. 10.1161/CIRCRESAHA.122.321769.

24. Dardano, M., Kleemiss, F., Kosanke, M., Lang, D., Wilson, L., Franke, A., Teske, J., Shivaraj, A., de la Roche, J., Fischer, M., et al. (2024). Blood-generating heart-forming organoids recapitulate co-development of the human haematopoietic system and the embryonic heart. Nature cell biology 26, 1984–1996. 10.1038/s41556-024-01526-4.

25. Kostina, A., Volmert, B., and Aguirre, A. (2024). Human heart organoids: current applications and future perspectives. Eur Heart J 45, 751–753 10.1093/eurheartj/ehad841.

26. Litvinukova, M., Talavera-Lopez, C., Maatz, H., Reichart, D., Worth, C.L., Lindberg, E.L., Kanda, M., Polanski, K., Heinig, M., Lee, M., et al. (2020). Cells of the adult human heart. Nature 588, 466–472. 10.1038/s41586-020-2797-4.

27. O’Hern, C., Caywood, S., Aminova, S., Kiselev, A., Volmert, B., Wang, F., Sewavi, M.-L., Cao, W., Dionise, M., Muniyandi, P., et al. (2024). 10.1101/2024.11.13.623051.

28. Szabo, A., and Mayor, R. (2018). Mechanisms of Neural Crest Migration. Annu Rev Genet 52, 43–63. 10.1146/annurev-genet-120417-031559.

29. Creazzo, T.L., Godt, R.E., Leatherbury, L., Conway, S.J., and Kirby, M.L. (1998). Role of cardiac neural crest cells in cardiovascular development. Annu Rev Physiol 60, 267–286. 10.1146/annurev.physiol.60.1.267.

30. Keyte, A., and Hutson, M.R. (2012). The neural crest in cardiac congenital anomalies. Differentiation 84, 25–40. 10.1016/j.diff.2012.04.005.

31. Erhardt, S., Zheng, M., Zhao, X., Le, T.P., Findley, T.O., and Wang, J. (2021). The Cardiac Neural Crest Cells in Heart Development and Congenital Heart Defects. J Cardiovasc Dev Dis 8. 10.3390/jcdd8080089.

32. Kirby, M.L., Gale, T.F., and Stewart, D.E. (1983). Neural crest cells contribute to normal aorticopulmonary septation. Science 220, 1059–1061. 10.1126/science.6844926.

33. Yamagishi, H. (2021). Cardiac Neural Crest. Cold Spring Harb Perspect Biol 13. 10.1101/cshperspect.a036715.

34. Waldo, K., Miyagawa-Tomita, S., Kumiski, D., and Kirby, M.L. (1998). Cardiac neural crest cells provide new insight into septation of the cardiac outflow tract: aortic sac to ventricular septal closure. Dev Biol 196, 129–144. 10.1006/dbio.1998.8860.

35. Waldo, K.L., Hutson, M.R., Stadt, H.A., Zdanowicz, M., Zdanowicz, J., and Kirby, M.L. (2005). Cardiac neural crest is necessary for normal addition of the myocardium to the arterial pole from the secondary heart field. Dev Biol 281, 66–77. 10.1016/j.ydbio.2005.02.011.

36. Pearlstein, T. (2015). Depression during Pregnancy. Best Pract Res Clin Obstet Gynaecol 29, 754–764. 10.1016/j.bpobgyn.2015.04.004.

37. Loughhead, A.M., Fisher, A.D., Newport, D.J., Ritchie, J.C., Owens, M.J., DeVane, C.L., and Stowe, Z.N. (2006). Antidepressants in amniotic fluid: another route of fetal exposure. Am J Psychiatry 163, 145–147. 10.1176/appi.ajp.163.1.145.

38. Sadler, T.W. (2011). Selective serotonin reuptake inhibitors (SSRIs) and heart defects: potential mechanisms for the observed associations. Reprod Toxicol 32, 484–489. 10.1016/j.reprotox.2011.09.004.

39. Yavarone, M.S., Shuey, D.L., Tamir, H., Sadler, T.W., and Lauder, J.M. (1993). Serotonin and cardiac morphogenesis in the mouse embryo. Teratology 47, 573–584. 10.1002/tera.1420470609.

40. Berard, A. (2010). Paroxetine exposure during pregnancy and the risk of cardiac malformations: what is the evidence? Birth Defects Res A Clin Mol Teratol 88, 171–174. 10.1002/bdra.20643.

41. Berard, A., Iessa, N., Chaabane, S., Muanda, F.T., Boukhris, T., and Zhao, J.P. (2016). The risk of major cardiac malformations associated with paroxetine use during the first trimester of pregnancy: a systematic review and meta-analysis. Br J Clin Pharmacol 81, 589–604. 10.1111/bcp.12849.

42. Gao, S.Y., Wu, Q.J., Zhang, T.N., Shen, Z.Q., Liu, C.X., Xu, X., Ji, C., and Zhao, Y.H. (2017). Fluoxetine and congenital malformations: a systematic review and meta-analysis of cohort studies. Br J Clin Pharmacol 83, 2134–2147. 10.1111/bcp.13321.

43. Shen, Z.Q., Gao, S.Y., Li, S.X., Zhang, T.N., Liu, C.X., Lv, H.C., Zhang, Y., Gong, T.T., Xu, X., Ji, C., et al. (2017). Sertraline use in the first trimester and risk of congenital anomalies: a systemic review and meta-analysis of cohort studies. Br J Clin Pharmacol 83, 909–922. 10.1111/bcp.13161.

44. Hackland, J.O.S., Frith, T.J.R., Thompson, O., Marin Navarro, A., Garcia-Castro, M.I., Unger, C., and Andrews, P.W. (2017). Top-Down Inhibition of BMP Signaling Enables Robust Induction of hPSCs Into Neural Crest in Fully Defined, Xeno-free Conditions. Stem cell reports 9, 1043–1052. 10.1016/j.stemcr.2017.08.008.

45. Feiner, L., Webber, A.L., Brown, C.B., Lu, M.M., Jia, L., Feinstein, P., Mombaerts, P., Epstein, J.A., and Raper, J.A. (2001). Targeted disruption of semaphorin 3C leads to persistent truncus arteriosus and aortic arch interruption. Development 128, 3061–3070. 10.1242/dev.128.16.3061.

46. Epstein, J.A., Aghajanian, H., and Singh, M.K. (2015). Semaphorin signaling in cardiovascular development. Cell Metab 21, 163–173. 10.1016/j.cmet.2014.12.015.

47. Toyofuku, T., Yoshida, J., Sugimoto, T., Yamamoto, M., Makino, N., Takamatsu, H., Takegahara, N., Suto, F., Hori, M., Fujisawa, H., et al. (2008). Repulsive and attractive semaphorins cooperate to direct the navigation of cardiac neural crest cells. Dev Biol 321, 251–262. 10.1016/j.ydbio.2008.06.028.

48. Fan, X., Masamsetti, V.P., Sun, J.Q., Engholm-Keller, K., Osteil, P., Studdert, J., Graham, M.E., Fossat, N., and Tam, P.P. (2021). TWIST1 and chromatin regulatory proteins interact to guide neural crest cell differentiation. Elife 10. 10.7554/eLife.62873.

49. Vincentz, J.W., Firulli, B.A., Lin, A., Spicer, D.B., Howard, M.J., and Firulli, A.B. (2013). Twist1 controls a cell-specification switch governing cell fate decisions within the cardiac neural crest. PLoS Genet 9, e1003405. 10.1371/journal.pgen.1003405.

50. Bertol, J.W., Johnston, S., Ahmed, R., Xie, V.K., Hubka, K.M., Cruz, L., Nitschke, L., Stetsiv, M., Goering, J.P., Nistor, P., et al. (2022). TWIST1 interacts with beta/delta-catenins during neural tube development and regulates fate transition in cranial neural crest cells. Development 149. 10.1242/dev.200068.

51. Cranley, J., Kanemaru, K., Bayraktar, S., Knight-Schrijver, V., Pett, J.P., Polanski, K., Dabrowska, M., Mulas, I., Richardson, L., Semprich, C.I., et al. (2024). 10.1101/2024.04.29.591736.

52. Bayraktar, S., Cranley, J., Kanemaru, K., Knight-Schrijver, V., Colzani, M., Davaapil, H., Lee, J.C.M., Polanski, K., Richardson, L., Semprich, C.I., et al. (2024). 10.1101/2024.04.27.591127.

53. Kanemaru, K., Cranley, J., Muraro, D., Miranda, A.M.A., Ho, S.Y., Wilbrey-Clark, A., Patrick Pett, J., Polanski, K., Richardson, L., Litvinukova, M., et al. (2023). Spatially resolved multiomics of human cardiac niches. Nature 619, 801–810. 10.1038/s41586-023-06311-1.

54. Li, G., Xu, A., Sim, S., Priest, J.R., Tian, X., Khan, T., Quertermous, T., Zhou, B., Tsao, P.S., Quake, S.R., and Wu, S.M. (2016). Transcriptomic Profiling Maps Anatomically Patterned Subpopulations among Single Embryonic Cardiac Cells. Dev Cell 39, 491–507. 10.1016/j.devcel.2016.10.014.

55. Farah, E.N., Hu, R.K., Kern, C., Zhang, Q., Lu, T.Y., Ma, Q., Tran, S., Zhang, B., Carlin, D., Monell, A., et al. (2024). Spatially organized cellular communities form the developing human heart. Nature 627, 854–864. 10.1038/s41586-024-07171-z.

56. Cui, Y., Zheng, Y., Liu, X., Yan, L., Fan, X., Yong, J., Hu, Y., Dong, J., Li, Q., Wu, X., et al. (2019). Single-Cell Transcriptome Analysis Maps the Developmental Track of the Human Heart. Cell Rep 26, 1934–1950 e1935. 10.1016/j.celrep.2019.01.079.

57. Asp, M., Giacomello, S., Larsson, L., Wu, C., Furth, D., Qian, X., Wardell, E., Custodio, J., Reimegard, J., Salmen, F., et al. (2019). A Spatiotemporal Organ-Wide Gene Expression and Cell Atlas of the Developing Human Heart. Cell 179, 1647–1660 e1619. 10.1016/j.cell.2019.11.025.

58. Spurlock, B., and Qian, L. (2023). Tracing the history of a heart. Elife 12. 10.7554/eLife.89988.

59. Forte, E., Furtado, M.B., and Rosenthal, N. (2018). The interstitium in cardiac repair: role of the immune-stromal cell interplay. Nature reviews. Cardiology 15, 601–616. 10.1038/s41569-018-0077-x.

60. Kirby, M.L., and Hutson, M.R. (2010). Factors controlling cardiac neural crest cell migration. Cell Adh Migr 4, 609–621. 10.4161/cam.4.4.13489.

61. Gandhi, S., Ezin, M., and Bronner, M.E. (2020). Reprogramming Axial Level Identity to Rescue Neural-Crest-Related Congenital Heart Defects. Dev Cell 53, 300–315 e304. 10.1016/j.devcel.2020.04.005.

62. De Bono, C., Liu, Y., Ferrena, A., Valentine, A., Zheng, D., and Morrow, B.E. (2023). Single-cell transcriptomics uncovers a non-autonomous Tbx1-dependent genetic program controlling cardiac neural crest cell development. Nature communications 14, 1551. 10.1038/s41467-023-37015-9.

63. Chen, W., Liu, X., Li, W., Shen, H., Zeng, Z., Yin, K., Priest, J.R., and Zhou, Z. (2021). Single-cell transcriptomic landscape of cardiac neural crest cell derivatives during development. EMBO Rep 22, e52389. 10.15252/embr.202152389.

64. Hoover, L.L., Burton, E.G., Brooks, B.A., and Kubalak, S.W. (2008). The expanding role for retinoid signaling in heart development. ScientificWorldJournal 8, 194–211. 10.1100/tsw.2008.39.

65. Tomarev, S.I., and Nakaya, N. (2009). Olfactomedin domain-containing proteins: possible mechanisms of action and functions in normal development and pathology. Mol Neurobiol 40, 122–138. 10.1007/s12035-009-8076-x.

66. Soldatov, R., Kaucka, M., Kastriti, M.E., Petersen, J., Chontorotzea, T., Englmaier, L., Akkuratova, N., Yang, Y., Haring, M., Dyachuk, V., et al. (2019). Spatiotemporal structure of cell fate decisions in murine neural crest. Science 364. 10.1126/science.aas9536.

67. Guo, S., Wang, Y., and Wang, A. (2020). Identity and lineage fate of proteolipid protein 1 gene (Plp1)- expressing cells in the embryonic murine spinal cord. Developmental dynamics : an official publication of the American Association of Anatomists 249, 946–960. 10.1002/dvdy.184.

68. Baynash, A.G., Hosoda, K., Giaid, A., Richardson, J.A., Emoto, N., Hammer, R.E., and Yanagisawa, M. (1994). Interaction of endothelin-3 with endothelin-B receptor is essential for development of epidermal melanocytes and enteric neurons. Cell 79, 1277–1285. 10.1016/0092-8674(94)90018-3.

69. Shin, M.K., Levorse, J.M., Ingram, R.S., and Tilghman, S.M. (1999). The temporal requirement for endothelin receptor-B signalling during neural crest development. Nature 402, 496–501. 10.1038/990040.

70. Xie, D., Xiong, K., Su, X., Wang, G., Ji, Q., Zou, Q., Wang, L., Liu, Y., Liang, D., Xue, J., et al. (2021). Identification of an endogenous glutamatergic transmitter system controlling excitability and conductivity of atrial cardiomyocytes. Cell Res 31, 951–964. 10.1038/s41422-021-00499-5.

71. Xie, D., Xiong, K., Su, X., Wang, G., Zou, Q., Wang, L., Zhang, C., Cao, Y., Shao, B., Zhang, Y., et al. (2022). Glutamate drives ’local Ca(2+) release’ in cardiac pacemaker cells. Cell Res 32, 843–854. 10.1038/s41422-022-00693-z.

72. Acharya, A., Baek, S.T., Huang, G., Eskiocak, B., Goetsch, S., Sung, C.Y., Banfi, S., Sauer, M.F., Olsen, G.S., Duffield, J.S., et al. (2012). The bHLH transcription factor Tcf21 is required for lineage-specific EMT of cardiac fibroblast progenitors. Development 139, 2139–2149. 10.1242/dev.079970.

73. Dettman, R.W., Denetclaw, W., Jr., Ordahl, C.P., and Bristow, J. (1998). Common epicardial origin of coronary vascular smooth muscle, perivascular fibroblasts, and intermyocardial fibroblasts in the avian heart. Dev Biol 193, 169–181. 10.1006/dbio.1997.8801.

74. Gittenberger-de Groot, A.C., Vrancken Peeters, M.P., Mentink, M.M., Gourdie, R.G., and Poelmann, R.E. (1998). Epicardium-derived cells contribute a novel population to the myocardial wall and the atrioventricular cushions. Circulation research 82, 1043–1052. 10.1161/01.res.82.10.1043.

75. Tallquist, M.D. (2020). Developmental Pathways of Cardiac Fibroblasts. Cold Spring Harb Perspect Biol 12. 10.1101/cshperspect.a037184.

76. Mikawa, T., and Gourdie, R.G. (1996). Pericardial mesoderm generates a population of coronary smooth muscle cells migrating into the heart along with ingrowth of the epicardial organ. Dev Biol 174, 221–232. 10.1006/dbio.1996.0068.

77. Neeb, Z., Lajiness, J.D., Bolanis, E., and Conway, S.J. (2013). Cardiac outflow tract anomalies. Wiley Interdiscip Rev Dev Biol 2, 499–530. 10.1002/wdev.98.

78. Derrick, C.J., and Noel, E.S. (2021). The ECM as a driver of heart development and repair. Development 148. 10.1242/dev.191320.

79. Leshem, R., Baker, S.M., Mallen, J., Wang, L., Dark, J., Sharrocks, A.D., Hanley, K.P., Hanley, N.A., Rattray, M., Bamforth, S.D., and Bobola, N. (2024). 10.1101/2023.04.05.535627.

80. Harel, I., Maezawa, Y., Avraham, R., Rinon, A., Ma, H.Y., Cross, J.W., Leviatan, N., Hegesh, J., Roy, A., Jacob-Hirsch, J., et al. (2012). Pharyngeal mesoderm regulatory network controls cardiac and head muscle morphogenesis. Proc Natl Acad Sci U S A 109, 18839–18844. 10.1073/pnas.1208690109.

81. van Bezooijen, R.L., Deruiter, M.C., Vilain, N., Monteiro, R.M., Visser, A., van der Wee-Pals, L., van Munsteren, C.J., Hogendoorn, P.C., Aguet, M., Mummery, C.L., et al. (2007). SOST expression is restricted to the great arteries during embryonic and neonatal cardiovascular development. Developmental dynamics : an official publication of the American Association of Anatomists 236, 606–612. 10.1002/dvdy.21054.

82. Delwarde, C., Capoulade, R., Merot, J., Le Scouarnec, S., Bouatia-Naji, N., Yu, M., Huttin, O., Selton-Suty, C., Sellal, J.M., Piriou, N., et al. (2023). Genetics and pathophysiology of mitral valve prolapse. Front Cardiovasc Med 10, 1077788. 10.3389/fcvm.2023.1077788.

83. Martinsen, B.J., Frasier, A.J., Baker, C.V., and Lohr, J.L. (2004). Cardiac neural crest ablation alters Id2 gene expression in the developing heart. Dev Biol 272, 176–190. 10.1016/j.ydbio.2004.04.030.

84. Hu, W., Xin, Y., Hu, J., Sun, Y., and Zhao, Y. (2019). Inhibitor of DNA binding in heart development and cardiovascular diseases. Cell Commun Signal 17, 51. 10.1186/s12964-019-0365-z.

85. Jongbloed, M.R., Vicente-Steijn, R., Douglas, Y.L., Wisse, L.J., Mori, K., Yokota, Y., Bartelings, M.M., Schalij, M.J., Mahtab, E.A., Poelmann, R.E., and Gittenberger-De Groot, A.C. (2011). Expression of Id2 in the second heart field and cardiac defects in Id2 knock-out mice. Developmental dynamics : an official publication of the American Association of Anatomists 240, 2561–2577. 10.1002/dvdy.22762.

86. Liu, X., Chen, W., Li, W., Li, Y., Priest, J.R., Zhou, B., Wang, J., and Zhou, Z. (2019). Single-Cell RNA-Seq of the Developing Cardiac Outflow Tract Reveals Convergent Development of the Vascular Smooth Muscle Cells. Cell Rep 28, 1346–1361 e1344. 10.1016/j.celrep.2019.06.092.

87. Vegh, A.M.D., Duim, S.N., Smits, A.M., Poelmann, R.E., Ten Harkel, A.D.J., DeRuiter, M.C., Goumans, M.J., and Jongbloed, M.R.M. (2016). Part and Parcel of the Cardiac Autonomic Nerve System: Unravelling Its Cellular Building Blocks during Development. J Cardiovasc Dev Dis 3. 10.3390/jcdd3030028.

88. Kuang, X.L., Zhao, X.M., Xu, H.F., Shi, Y.Y., Deng, J.B., and Sun, G.T. (2010). Spatio-temporal expression of a novel neuron-derived neurotrophic factor (NDNF) in mouse brains during development. BMC Neurosci 11, 137. 10.1186/1471-2202-11-137.

89. Haenisch, C., Diekmann, H., Klinger, M., Gennarini, G., Kuwada, J.Y., and Stuermer, C.A. (2005). The neuronal growth and regeneration associated Cntn1 (F3/F11/Contactin) gene is duplicated in fish: expression during development and retinal axon regeneration. Mol Cell Neurosci 28, 361–374. 10.1016/j.mcn.2004.04.013.

90. Montani, C., Ramos-Brossier, M., Ponzoni, L., Gritti, L., Cwetsch, A.W., Braida, D., Saillour, Y., Terragni, B., Mantegazza, M., Sala, M., et al. (2017). The X-Linked Intellectual Disability Protein IL1RAPL1 Regulates Dendrite Complexity. J Neurosci 37, 6606–6627. 10.1523/JNEUROSCI.3775-16.2017.

91. Morimoto, Y., Tokumitsu, A., Sone, T., Hirota, Y., Tamura, R., Sakamoto, A., Nakajima, K., Toda, M., Kawakami, Y., Okano, H., and Ohta, S. (2022). TPT1 Supports Proliferation of Neural Stem/Progenitor Cells and Brain Tumor Initiating Cells Regulated by Macrophage Migration Inhibitory Factor (MIF). Neurochem Res 47, 2741–2756. 10.1007/s11064-022-03629-6.

92. Vidal, O.M., Velez, J.I., and Arcos-Burgos, M. (2022). ADGRL3 genomic variation implicated in neurogenesis and ADHD links functional effects to the incretin polypeptide GIP. Scientific reports 12, 15922. 10.1038/s41598-022-20343-z.

93. Rigter, P.M.F., de Konink, C., Dunn, M.J., Proietti Onori, M., Humberson, J.B., Thomas, M., Barnes, C., Prada, C.E., Weaver, K.N., Ryan, T.D., et al. (2024). Role of CAMK2D in neurodevelopment and associated conditions. Am J Hum Genet 111, 364–382. 10.1016/j.ajhg.2023.12.016.

94. Allen, N.J., Bennett, M.L., Foo, L.C., Wang, G.X., Chakraborty, C., Smith, S.J., and Barres, B.A. (2012). Astrocyte glypicans 4 and 6 promote formation of excitatory synapses via GluA1 AMPA receptors. Nature 486, 410–414. 10.1038/nature11059.

95. Kirby, M.L., and Stewart, D.E. (1983). Neural crest origin of cardiac ganglion cells in the chick embryo: identification and extirpation. Dev Biol 97, 433–443. 10.1016/0012-1606(83)90100-8.

96. Fedele, L., and Brand, T. (2020). The Intrinsic Cardiac Nervous System and Its Role in Cardiac Pacemaking and Conduction. J Cardiovasc Dev Dis 7. 10.3390/jcdd7040054.

97. Hildreth, V., Webb, S., Bradshaw, L., Brown, N.A., Anderson, R.H., and Henderson, D.J. (2008). Cells migrating from the neural crest contribute to the innervation of the venous pole of the heart. J Anat 212, 1–11. 10.1111/j.1469-7580.2007.00833.x.

98. Pattyn, A., Morin, X., Cremer, H., Goridis, C., and Brunet, J.F. (1999). The homeobox gene Phox2b is essential for the development of autonomic neural crest derivatives. Nature 399, 366–370. 10.1038/20700.

99. Huang, M.H., Friend, D.S., Sunday, M.E., Singh, K., Haley, K., Austen, K.F., Kelly, R.A., and Smith, T.W. (1996). An intrinsic adrenergic system in mammalian heart. J Clin Invest 98, 1298–1303. 10.1172/JCI118916.

100. Giannino, G., Braia, V., Griffith Brookles, C., Giacobbe, F., D’Ascenzo, F., Angelini, F., Saglietto, A., De Ferrari, G.M., and Dusi, V. (2024). The Intrinsic Cardiac Nervous System: From Pathophysiology to Therapeutic Implications. Biology (Basel) 13. 10.3390/biology13020105.

101. Madrigal, M.P., Portales, A., SanJuan, M.P., and Jurado, S. (2019). Postsynaptic SNARE Proteins: Role in Synaptic Transmission and Plasticity. Neuroscience 420, 12–21. 10.1016/j.neuroscience.2018.11.012.

102. Sauvola, C.W., and Littleton, J.T. (2021). SNARE Regulatory Proteins in Synaptic Vesicle Fusion and Recycling. Front Mol Neurosci 14, 733138. 10.3389/fnmol.2021.733138.

103. Danielson, E., Zhang, N., Metallo, J., Kaleka, K., Shin, S.M., Gerges, N., and Lee, S.H. (2012). S-SCAM/MAGI-2 is an essential synaptic scaffolding molecule for the GluA2-containing maintenance pool of AMPA receptors. J Neurosci 32, 6967–6980. 10.1523/JNEUROSCI.0025-12.2012.

104. Kimura, S., Lok, J., Gelman, I.H., Lo, E.H., and Arai, K. (2023). Role of A-Kinase Anchoring Protein 12 in the Central Nervous System. J Clin Neurol 19, 329–337. 10.3988/jcn.2023.0095.

105. Chiba, K., Kita, T., Anazawa, Y., and Niwa, S. (2023). Insight into the regulation of axonal transport from the study of KIF1A-associated neurological disorder. J Cell Sci 136. 10.1242/jcs.260742.

106. Zhao, M., Maani, N., and Dowling, J.J. (2018). Dynamin 2 (DNM2) as Cause of, and Modifier for, Human Neuromuscular Disease. Neurotherapeutics 15, 966–975. 10.1007/s13311-018-00686-0.

107. Biederer, T., Kaeser, P.S., and Blanpied, T.A. (2017). Transcellular Nanoalignment of Synaptic Function. Neuron 96, 680–696. 10.1016/j.neuron.2017.10.006.

108. Hasan, W. (2013). Autonomic cardiac innervation: development and adult plasticity. Organogenesis 9, 176–193. 10.4161/org.24892.

109. Chen, T.W., Wardill, T.J., Sun, Y., Pulver, S.R., Renninger, S.L., Baohan, A., Schreiter, E.R., Kerr, R.A., Orger, M.B., Jayaraman, V., et al. (2013). Ultrasensitive fluorescent proteins for imaging neuronal activity. Nature 499, 295–300. 10.1038/nature12354.

110. Kikel-Coury, N.L., Brandt, J.P., Correia, I.A., O’Dea, M.R., DeSantis, D.F., Sterling, F., Vaughan, K., Ozcebe, G., Zorlutuna, P., and Smith, C.J. (2021). Identification of astroglia-like cardiac nexus glia that are critical regulators of cardiac development and function. PLoS Biol 19, e3001444. 10.1371/journal.pbio.3001444.

111. Jessen, K.R., and Mirsky, R. (2005). The origin and development of glial cells in peripheral nerves. Nat Rev Neurosci 6, 671–682. 10.1038/nrn1746.

112. Hutson, M.R., and Kirby, M.L. (2007). Model systems for the study of heart development and disease. Cardiac neural crest and conotruncal malformations. Semin Cell Dev Biol 18, 101–110. 10.1016/j.semcdb.2006.12.004.

113. Chen, Y.H., Ishii, M., Sun, J., Sucov, H.M., and Maxson, R.E., Jr. (2007). Msx1 and Msx2 regulate survival of secondary heart field precursors and post-migratory proliferation of cardiac neural crest in the outflow tract. Dev Biol 308, 421–437. 10.1016/j.ydbio.2007.05.037.

114. Xu, K., Xie, S., Huang, Y., Zhou, T., Liu, M., Zhu, P., Wang, C., Shi, J., Li, F., Sellke, F.W., and Dong, N. (2020). Cell-Type Transcriptome Atlas of Human Aortic Valves Reveal Cell Heterogeneity and Endothelial to Mesenchymal Transition Involved in Calcific Aortic Valve Disease. Arteriosclerosis, thrombosis, and vascular biology 40, 2910–2921. 10.1161/ATVBAHA.120.314789.

115. Fu, M., and Song, J. (2023). Single-cell RNA sequencing reveals the diversity and biology of valve cells in cardiac valve disease. J Cardiol 81, 49–56. 10.1016/j.jjcc.2022.03.012.

116. Peacock, J.D., Lu, Y., Koch, M., Kadler, K.E., and Lincoln, J. (2008). Temporal and spatial expression of collagens during murine atrioventricular heart valve development and maintenance. Developmental dynamics : an official publication of the American Association of Anatomists 237, 3051–3058. 10.1002/dvdy.21719.

117. Cai, Z., Zhu, M., Xu, L., Wang, Y., Xu, Y., Yim, W.Y., Cao, H., Guo, R., Qiu, X., He, X., et al. (2024). Directed Differentiation of Human Induced Pluripotent Stem Cells to Heart Valve Cells. Circulation 149, 1435–1456. 10.1161/CIRCULATIONAHA.123.065143.

118. Hulin, A., Hortells, L., Gomez-Stallons, M.V., O’Donnell, A., Chetal, K., Adam, M., Lancellotti, P., Oury, C., Potter, S.S., Salomonis, N., and Yutzey, K.E. (2019). Maturation of heart valve cell populations during postnatal remodeling. Development 146. 10.1242/dev.173047.

119. Camenisch, T.D., Spicer, A.P., Brehm-Gibson, T., Biesterfeldt, J., Augustine, M.L., Calabro, A., Jr., Kubalak, S., Klewer, S.E., and McDonald, J.A. (2000). Disruption of hyaluronan synthase-2 abrogates normal cardiac morphogenesis and hyaluronan-mediated transformation of epithelium to mesenchyme. J Clin Invest 106, 349–360. 10.1172/JCI10272.

120. Jia, Q., McDill, B.W., Li, S.Z., Deng, C., Chang, C.P., and Chen, F. (2007). Smad signaling in the neural crest regulates cardiac outflow tract remodeling through cell autonomous and non-cell autonomous effects. Dev Biol 311, 172–184. 10.1016/j.ydbio.2007.08.044.

121. Bowen, C.J., Zhou, J., Sung, D.C., and Butcher, J.T. (2015). Cadherin-11 coordinates cellular migration and extracellular matrix remodeling during aortic valve maturation. Dev Biol 407, 145–157. 10.1016/j.ydbio.2015.07.012.

122. Nakhai-Pour, H.R., Broy, P., and Berard, A. (2010). Use of antidepressants during pregnancy and the risk of spontaneous abortion. CMAJ 182, 1031–1037. 10.1503/cmaj.091208.

123. Einarson, A., Choi, J., Einarson, T.R., and Koren, G. (2009). Rates of spontaneous and therapeutic abortions following use of antidepressants in pregnancy: results from a large prospective database. J Obstet Gynaecol Can 31, 452–456. 10.1016/s1701-2163(16)34177-9.

124. Broy, P., and Berard, A. (2010). Gestational exposure to antidepressants and the risk of spontaneous abortion: a review. Curr Drug Deliv 7, 76–92. 10.2174/156720110790396508.

125. Berard, A., Zhao, J.P., and Sheehy, O. (2015). Sertraline use during pregnancy and the risk of major malformations. Am J Obstet Gynecol 212, 795 e791–795 e712. 10.1016/j.ajog.2015.01.034.

126. Desai, P.H., Yagnik, P.J., Ross Ascuitto, N., Prajapati, P., and Sernich, S. (2019). Risk of Congenital Heart Disease in Newborns with Prenatal Exposure to Anti-depressant Medications. Cureus 11, e4673. 10.7759/cureus.4673.

127. El Marroun, H., Jaddoe, V.W., Hudziak, J.J., Roza, S.J., Steegers, E.A., Hofman, A., Verhulst, F.C., White, T.J., Stricker, B.H., and Tiemeier, H. (2012). Maternal use of selective serotonin reuptake inhibitors, fetal growth, and risk of adverse birth outcomes. Arch Gen Psychiatry 69, 706–714. 10.1001/archgenpsychiatry.2011.2333.

128. Jimenez-Solem, E., Andersen, J.T., Petersen, M., Broedbaek, K., Jensen, J.K., Afzal, S., Gislason, G.H., Torp-Pedersen, C., and Poulsen, H.E. (2012). Exposure to selective serotonin reuptake inhibitors and the risk of congenital malformations: a nationwide cohort study. BMJ Open 2. 10.1136/bmjopen-2012-001148.

129. Gao, S.Y., Wu, Q.J., Sun, C., Zhang, T.N., Shen, Z.Q., Liu, C.X., Gong, T.T., Xu, X., Ji, C., Huang, D.H., et al. (2018). Selective serotonin reuptake inhibitor use during early pregnancy and congenital malformations: a systematic review and meta-analysis of cohort studies of more than 9 million births. BMC Med 16, 205. 10.1186/s12916-018-1193-5.

130. Huybrechts, K.F., Palmsten, K., Avorn, J., Cohen, L.S., Holmes, L.B., Franklin, J.M., Mogun, H., Levin, R., Kowal, M., Setoguchi, S., and Hernandez-Diaz, S. (2014). Antidepressant use in pregnancy and the risk of cardiac defects. N Engl J Med 370, 2397–2407. 10.1056/NEJMoa1312828.

131. Lou, Z.Q., Zhou, Y.Y., Zhang, X., and Jiang, H.Y. (2022). Exposure to selective noradrenalin reuptake inhibitors during the first trimester of pregnancy and risk of congenital malformations: A meta-analysis of cohort studies. Psychiatry Res 316, 114756. 10.1016/j.psychres.2022.114756.

132. Riggin, L., Frankel, Z., Moretti, M., Pupco, A., and Koren, G. (2013). The fetal safety of fluoxetine: a systematic review and meta-analysis. J Obstet Gynaecol Can 35, 362–369. 10.1016/S1701-2163(15)30965-8.

133. Gaultier, C., Trang, H., Dauger, S., and Gallego, J. (2005). Pediatric disorders with autonomic dysfunction: what role for PHOX2B? Pediatr Res 58, 1–6. 10.1203/01.PDR.0000166755.29277.C4.

134. Muller, F., and Rohrer, H. (2002). Molecular control of ciliary neuron development: BMPs and downstream transcriptional control in the parasympathetic lineage. Development 129, 5707–5717. 10.1242/dev.00165.

135. Thompson, W.A., Shvartsburd, Z., and Vijayan, M.M. (2022). The antidepressant venlafaxine perturbs cardiac development and function in larval zebrafish. Aquat Toxicol 242, 106041. 10.1016/j.aquatox.2021.106041.

136. Meier, A.B., Zawada, D., De Angelis, M.T., Martens, L.D., Santamaria, G., Zengerle, S., Nowak-Imialek, M., Kornherr, J., Zhang, F., Tian, Q., et al. (2023). Epicardioid single-cell genomics uncovers principles of human epicardium biology in heart development and disease. Nat Biotechnol 41, 1787–1800. 10.1038/s41587-023-01718-7.

137. Feng, W., Schriever, H., Jiang, S., Bais, A., Wu, H., Kostka, D., and Li, G. (2022). Computational profiling of hiPSC-derived heart organoids reveals chamber defects associated with NKX2-5 deficiency. Commun Biol 5, 399. 10.1038/s42003-022-03346-4.

138. Brown, C.B., Feiner, L., Lu, M.M., Li, J., Ma, X., Webber, A.L., Jia, L., Raper, J.A., and Epstein, J.A. (2001). PlexinA2 and semaphorin signaling during cardiac neural crest development. Development 128, 3071–3080. 10.1242/dev.128.16.3071.

139. Kodo, K., Shibata, S., Miyagawa-Tomita, S., Ong, S.G., Takahashi, H., Kume, T., Okano, H., Matsuoka, R., and Yamagishi, H. (2017). Regulation of Sema3c and the Interaction between Cardiac Neural Crest and Second Heart Field during Outflow Tract Development. Scientific reports 7, 6771. 10.1038/s41598-017-06964-9.

140. Zhao, R., Moore, E.L., Gogol, M.M., Unruh, J.R., Yu, Z., Scott, A., Wang, Y., Rajendran, N.K., and Trainor, P.A. (2024). Identification and characterization of intermediate states in mammalian neural crest cell epithelial to mesenchymal transition and delamination. bioRxiv. 10.1101/2023.10.26.564204.

141. Alexander, B.E., Zhao, H., and Astrof, S. (2023). SMAD4: A Critical Regulator of Cardiac Neural Crest Cell Fate and Vascular Smooth Muscle Differentiation. bioRxiv. 10.1101/2023.03.14.532676.

142. De Bono, C., Liu, Y., Ferrena, A., Valentine, A., Zheng, D., and Morrow, B.E. (2022). 10.1101/2022.08.01.502391.

143. Iwase, A., Uchijima, Y., Seya, D., Kida, M., Higashiyama, H., Matsui, K., Taguchi, A., Yamamoto, S., Fukuda, S., Nomura, S., et al. (2022). 10.1101/2022.06.23.497419.

144. Moss, A., Robbins, S., Achanta, S., Kuttippurathu, L., Turick, S., Nieves, S., Hanna, P., Smith, E.H., Hoover, D.B., Chen, J., et al. (2021). A single cell transcriptomics map of paracrine networks in the intrinsic cardiac nervous system. iScience 24, 102713. 10.1016/j.isci.2021.102713.

145. Schneider, L.V., Guobin, B., Methi, A., Jensen, O., Schmoll, K.A., Setya, M.G., Sakib, S., Fahud, A.L., Brockmöller, J., Fischer, A., et al. (2023). 10.1101/2023.08.18.552653.

146. Zeltner, N., Wu, H.F., Saito-Diaz, K., Sun, X., Song, M., Saini, T., Grant, C., James, C., Thomas, K., Abate, Y., et al. (2024). A modular platform to generate functional sympathetic neuron-innervated heart assembloids. Res Sq. 10.21203/rs.3.rs-3894397/v1.

147. Workman, M.J., Mahe, M.M., Trisno, S., Poling, H.M., Watson, C.L., Sundaram, N., Chang, C.F., Schiesser, J., Aubert, P., Stanley, E.G., et al. (2017). Engineered human pluripotent-stem-cell-derived intestinal tissues with a functional enteric nervous system. Nat Med 23, 49–59. 10.1038/nm.4233.

148. Rockel, A.F., Wagner, N., Spenger, P., Ergun, S., and Worsdorfer, P. (2023). Neuro-mesodermal assembloids recapitulate aspects of peripheral nervous system development in vitro. Stem cell reports 18, 1155–1165. 10.1016/j.stemcr.2023.03.012.

149. Fregoso, S.P., and Hoover, D.B. (2012). Development of cardiac parasympathetic neurons, glial cells, and regional cholinergic innervation of the mouse heart. Neuroscience 221, 28–36. 10.1016/j.neuroscience.2012.06.061.

150. Gittenberger-de Groot, A.C., Blom, N.M., Aoyama, N., Sucov, H., Wenink, A.C., and Poelmann, R.E. (2003). The role of neural crest and epicardium-derived cells in conduction system formation. Novartis Found Symp 250, 125–134; discussion 134-141, 276-129. 10.1002/0470868066.ch8.

151. Takeuchi, A., Shimba, K., Mori, M., Takayama, Y., Moriguchi, H., Kotani, K., Lee, J.K., Noshiro, M., and Jimbo, Y. (2012). Sympathetic neurons modulate the beat rate of pluripotent cell-derived cardiomyocytes in vitro. Integr Biol (Camb) 4, 1532–1539. 10.1039/c2ib20060k.

152. Bernardin, A.A., Colombani, S., Rousselot, A., Andry, V., Goumon, Y., Delanoe-Ayari, H., Pasqualin, C., Brugg, B., Jacotot, E.D., Pasquie, J.L., et al. (2022). Impact of Neurons on Patient-Derived Cardiomyocytes Using Organ-On-A-Chip and iPSC Biotechnologies. Cells 11. 10.3390/cells11233764.

153. Winbo, A., Ramanan, S., Eugster, E., Jovinge, S., Skinner, J.R., and Montgomery, J.M. (2020). Functional coculture of sympathetic neurons and cardiomyocytes derived from human-induced pluripotent stem cells. American journal of physiology. Heart and circulatory physiology 319, H927–H937. 10.1152/ajpheart.00546.2020.

154. Dokshokova, L., Franzoso, M., Di Bona, A., Moro, N., Sanchez Alonso, J.L., Prando, V., Sandre, M., Basso, C., Faggian, G., Abriel, H., et al. (2022). Nerve growth factor transfer from cardiomyocytes to innervating sympathetic neurons activates TrkA receptors at the neuro-cardiac junction. J Physiol 600, 2853–2875. 10.1113/JP282828.

155. Takayama, Y., Kushige, H., Akagi, Y., Suzuki, Y., Kumagai, Y., and Kida, Y.S. (2020). Selective Induction of Human Autonomic Neurons Enables Precise Control of Cardiomyocyte Beating. Scientific reports 10, 9464. 10.1038/s41598-020-66303-3.

156. Oh, Y., Cho, G.S., Li, Z., Hong, I., Zhu, R., Kim, M.J., Kim, Y.J., Tampakakis, E., Tung, L., Huganir, R., et al. (2016). Functional Coupling with Cardiac Muscle Promotes Maturation of hPSC-Derived Sympathetic Neurons. Cell stem cell 19, 95–106. 10.1016/j.stem.2016.05.002.

157. Takeuchi, A., Shimba, K., Takayama, Y., Kotani, K., Lee, J.K., Noshiro, M., and Jimbo, Y. (2013). Microfabricated device for co-culture of sympathetic neuron and iPS-derived cardiomyocytes. Annu Int Conf IEEE Eng Med Biol Soc 2013, 3817–3820. 10.1109/EMBC.2013.6610376.

158. Hakli, M., Jantti, S., Joki, T., Sukki, L., Tornberg, K., Aalto-Setala, K., Kallio, P., Pekkanen-Mattila, M., and Narkilahti, S. (2022). Human Neurons Form Axon-Mediated Functional Connections with Human Cardiomyocytes in Compartmentalized Microfluidic Chip. International journal of molecular sciences 23. 10.3390/ijms23063148.

159. Nakamura, T., Colbert, M.C., and Robbins, J. (2006). Neural crest cells retain multipotential characteristics in the developing valves and label the cardiac conduction system. Circulation research 98, 1547–1554. 10.1161/01.RES.0000227505.19472.69.

160. Gurjarpadhye, A., Hewett, K.W., Justus, C., Wen, X., Stadt, H., Kirby, M.L., Sedmera, D., and Gourdie, R.G. (2007). Cardiac neural crest ablation inhibits compaction and electrical function of conduction system bundles. American journal of physiology. Heart and circulatory physiology 292, H1291–1300. 10.1152/ajpheart.01017.2006.

161. Dolejsi, E., Janouskova, A., and Jakubik, J. (2024). Muscarinic Receptors in Cardioprotection and Vascular Tone Regulation. Physiol Res 73, S389–S400. 10.33549/physiolres.935270.

162. Lara, A., Damasceno, D.D., Pires, R., Gros, R., Gomes, E.R., Gavioli, M., Lima, R.F., Guimaraes, D., Lima, P., Bueno, C.R., Jr., et al. (2010). Dysautonomia due to reduced cholinergic neurotransmission causes cardiac remodeling and heart failure. Molecular and cellular biology 30, 1746–1756. 10.1128/MCB.00996-09.

163. Sabbah, H.N. (2011). Electrical vagus nerve stimulation for the treatment of chronic heart failure. Cleve Clin J Med 78 *Suppl 1*, S24–29. 10.3949/ccjm.78.s1.04.

164. De Ferrari, G.M., Crijns, H.J., Borggrefe, M., Milasinovic, G., Smid, J., Zabel, M., Gavazzi, A., Sanzo, A., Dennert, R., Kuschyk, J., et al. (2011). Chronic vagus nerve stimulation: a new and promising therapeutic approach for chronic heart failure. Eur Heart J 32, 847–855. 10.1093/eurheartj/ehq391.

165. Tang, W., Martik, M.L., Li, Y., and Bronner, M.E. (2019). Cardiac neural crest contributes to cardiomyocytes in amniotes and heart regeneration in zebrafish. Elife 8. 10.7554/eLife.47929.

166. Cavanaugh, A.M., Huang, J., and Chen, J.N. (2015). Two developmentally distinct populations of neural crest cells contribute to the zebrafish heart. Dev Biol 404, 103–112. 10.1016/j.ydbio.2015.06.002.

167. Morrison, J.A., Pushel, I., McLennan, R., McKinney, M.C., Gogol, M.M., Scott, A., Krumlauf, R., and Kulesa, P.M. (2024). Comparative Analysis of Neural Crest Development in the Chick and Mouse. bioRxiv. 10.1101/2024.11.06.622355.

168. Hortells, L., Meyer, E.C., Thomas, Z.M., and Yutzey, K.E. (2021). Periostin-expressing Schwann cells and endoneurial cardiac fibroblasts contribute to sympathetic nerve fasciculation after birth. Journal of molecular and cellular cardiology 154, 124–136. 10.1016/j.yjmcc.2021.02.001.

169. Learte, A.R., and Hidalgo, A. (2007). The role of glial cells in axon guidance, fasciculation and targeting. Adv Exp Med Biol 621, 156–166. 10.1007/978-0-387-76715-4_12.

170. Scherschel, K., Hedenus, K., Jungen, C., Lemoine, M.D., Rubsamen, N., Veldkamp, M.W., Klatt, N., Lindner, D., Westermann, D., Casini, S., et al. (2019). Cardiac glial cells release neurotrophic S100B upon catheter-based treatment of atrial fibrillation. Sci Transl Med 11. 10.1126/scitranslmed.aav7770.

171. Watabe, K., Fukuda, T., Tanaka, J., Honda, H., Toyohara, K., and Sakai, O. (1995). Spontaneously immortalized adult mouse Schwann cells secrete autocrine and paracrine growth-promoting activities. J Neurosci Res 41, 279–290. 10.1002/jnr.490410215.

172. Kirby, M.L., and Waldo, K.L. (1995). Neural crest and cardiovascular patterning. Circulation research 77, 211–215. 10.1161/01.res.77.2.211.

173. Jain, R., Engleka, K.A., Rentschler, S.L., Manderfield, L.J., Li, L., Yuan, L., and Epstein, J.A. (2011). Cardiac neural crest orchestrates remodeling and functional maturation of mouse semilunar valves. J Clin Invest 121, 422–430. 10.1172/JCI44244.

174. Mekontso-Dessap, A., Brouri, F., Pascal, O., Lechat, P., Hanoun, N., Lanfumey, L., Seif, I., Benhaiem-Sigaux, N., Kirsch, M., Hamon, M., et al. (2006). Deficiency of the 5-hydroxytryptamine transporter gene leads to cardiac fibrosis and valvulopathy in mice. Circulation 113, 81–89. 10.1161/CIRCULATIONAHA.105.554667.

175. Choi, D.S., Ward, S.J., Messaddeq, N., Launay, J.M., and Maroteaux, L. (1997). 5-HT2B receptor-mediated serotonin morphogenetic functions in mouse cranial neural crest and myocardiac cells. Development 124, 1745–1755. 10.1242/dev.124.9.1745.

176. Moiseiwitsch, J.R., and Lauder, J.M. (1995). Serotonin regulates mouse cranial neural crest migration. Proc Natl Acad Sci U S A 92, 7182–7186. 10.1073/pnas.92.16.7182.

177. Hansson, S.R., Mezey, E., and Hoffman, B.J. (1999). Serotonin transporter messenger RNA expression in neural crest-derived structures and sensory pathways of the developing rat embryo. Neuroscience 89, 243–265. 10.1016/s0306-4522(98)00281-4.

178. Nebigil, C.G., Hickel, P., Messaddeq, N., Vonesch, J.L., Douchet, M.P., Monassier, L., Gyorgy, K., Matz, R., Andriantsitohaina, R., Manivet, P., et al. (2001). Ablation of serotonin 5-HT(2B) receptors in mice leads to abnormal cardiac structure and function. Circulation 103, 2973–2979. 10.1161/01.cir.103.24.2973.

179. Sari, Y., and Zhou, F.C. (2003). Serotonin and its transporter on proliferation of fetal heart cells. Int J Dev Neurosci 21, 417–424. 10.1016/j.ijdevneu.2003.10.002.

180. Ng, W.H., Johnston, E.K., Tan, J.J., Bliley, J.M., Feinberg, A.W., Stolz, D.B., Sun, M., Wijesekara, P., Hawkins, F., Kotton, D.N., and Ren, X. (2022). Recapitulating human cardio-pulmonary co-development using simultaneous multilineage differentiation of pluripotent stem cells. Elife 11. 10.7554/eLife.67872.

181. Lewis-Israeli, Y.R., Volmert, B.D., Gabalski, M.A., Huang, A.R., and Aguirre, A. (2021). Generating Self-Assembling Human Heart Organoids Derived from Pluripotent Stem Cells. J Vis Exp. 10.3791/63097.

182. Hao, Y., Stuart, T., Kowalski, M.H., Choudhary, S., Hoffman, P., Hartman, A., Srivastava, A., Molla, G., Madad, S., Fernandez-Granda, C., and Satija, R. (2024). Dictionary learning for integrative, multimodal and scalable single-cell analysis. Nat Biotechnol 42, 293–304. 10.1038/s41587-023-01767-y.

183. Korsunsky, I., Millard, N., Fan, J., Slowikowski, K., Zhang, F., Wei, K., Baglaenko, Y., Brenner, M., Loh, P.R., and Raychaudhuri, S. (2019). Fast, sensitive and accurate integration of single-cell data with Harmony. Nat Methods 16, 1289–1296. 10.1038/s41592-019-0619-0.

184. La Manno, G., Soldatov, R., Zeisel, A., Braun, E., Hochgerner, H., Petukhov, V., Lidschreiber, K., Kastriti, M.E., Lonnerberg, P., Furlan, A., et al. (2018). RNA velocity of single cells. Nature 560, 494–498. 10.1038/s41586-018-0414-6.

185. Bergen, V., Lange, M., Peidli, S., Wolf, F.A., and Theis, F.J. (2020). Generalizing RNA velocity to transient cell states through dynamical modeling. Nat Biotechnol 38, 1408–1414. 10.1038/s41587-020-0591-3.

186. Miller, S.A., Policastro, R.A., Sriramkumar, S., Lai, T., Huntington, T.D., Ladaika, C.A., Kim, D., Hao, C., Zentner, G.E., and O’Hagan, H.M. (2021). LSD1 and Aberrant DNA Methylation Mediate Persistence of Enteroendocrine Progenitors That Support BRAF-Mutant Colorectal Cancer. Cancer Res 81, 3791–3805. 10.1158/0008-5472.CAN-20-3562.

187. Aguirre, A., Huang, A.R., Gabalski, M.A., Volmert, B.D., and Lewis-Israeli, Y.R. (2021). Generating Self-Assembling Human Heart Organoids Derived from Pluripotent Stem Cells. Journal of Visualized Experiments. 10.3791/63097-v.

