## Supplementary video descriptions for "Self-organizing human heart assembloids with autologous and developmentally relevant cardiac neural crest-derived tissues"

**SUPPLEMENTARY VIDEO LEGENDS**

**Supplementary Video 1.**

A representative fluorescence video of a Day 19 hNCHA (2 days after mCherry-NCC integration) shows migrating NCCs with stable mCherry expression.

**Supplementary Video 2.**

A representative fluorescence video of a Day 19 hNCHA (2 days after mCherry-NCC integration) shows migrating NCCs with stable mCherry expression.

**Supplementary Video 3.**

A representative fluorescence video of a Day 19 hNCHA (2 days after mCherry-NCC integration) shows migrating NCCs with stable mCherry expression.

**Supplementary Video 4.**

A representative fluorescence video of a Day 21 hNCHA (4 days after mCherry-NCC integration) shows migrating NCCs with stable mCherry expression.

**Supplementary Video 5.**

A representative fluorescence video of a Day 23 hNCHA (6 days after mCherry-NCC integration) shows migrating NCCs with stable mCherry expression.

**Supplementary Video 6.**

A representative fluorescence video of a Day 30 hNCHA (13 days after mCherry-NCC integration) shows migrating NCCs with stable mCherry expression are closely associated with cardiomyocytes and form neuronal projections. Scale bar = 100 μm.

**Supplementary Video 7.**

A representative high magnification fluorescence video of a Day 30 hNCHA (13 days after mCherry-NCC integration) shows migrating NCCs with stable mCherry expression are closely associated with cardiomyocytes and form neuronal projections. Scale bar = 20 μm.

**Supplementary Video 8.**

A representative high magnification fluorescence video of a Day 30 hNCHA (13 days after mCherry-NCC integration) shows mCherry-NCCs-derived neuronal projections closely associated with cardiomyocytes. Scale bar = 20 μm.

**Supplementary Video 9.**

A representative high magnification fluorescence video of a Day 30 hNCHA (13 days after mCherry-NCC integration) shows mCherry-NCCs-derived neuronal projections closely associated with cardiomyocytes. Scale bar = 20 μm.

**Supplementary Video 10.**

Live imaging of calcium transients in GCaMP6f-NCC parasympathetic derivatives depicting neural activity in Day 60 hNCHAs. NCCs were derived from iPSC line containing GCaMP6f reporter. Video corresponds to Figure 5J, K.
