## Supplementary table legends for "Self-organizing human heart assembloids with autologous and developmentally relevant cardiac neural crest-derived tissues"

**Supplementary Table 1. List of Primers used in the study.**

**Supplementary Table 2. List of Antibodies used in the study.**

**Supplementary Table 3. Expression level and p-value of genes represented in Figure 2B.**

**Supplementary Table 4. Expression level and p-value of genes represented in Dot plots throughout the study.**
