## Supplementary figures for "Self-organizing human heart assembloids with autologous and developmentally relevant cardiac neural crest-derived tissues"

***IN BRIEF***

Human neural crest heart assembloids resembling the major directions of neural crest differentiation in the human embryonic heart, including parasympathetic innervation and the mesenchymal component of the outflow tract, provide a human-relevant embryonic platform for studying congenital heart defects and drug safety.

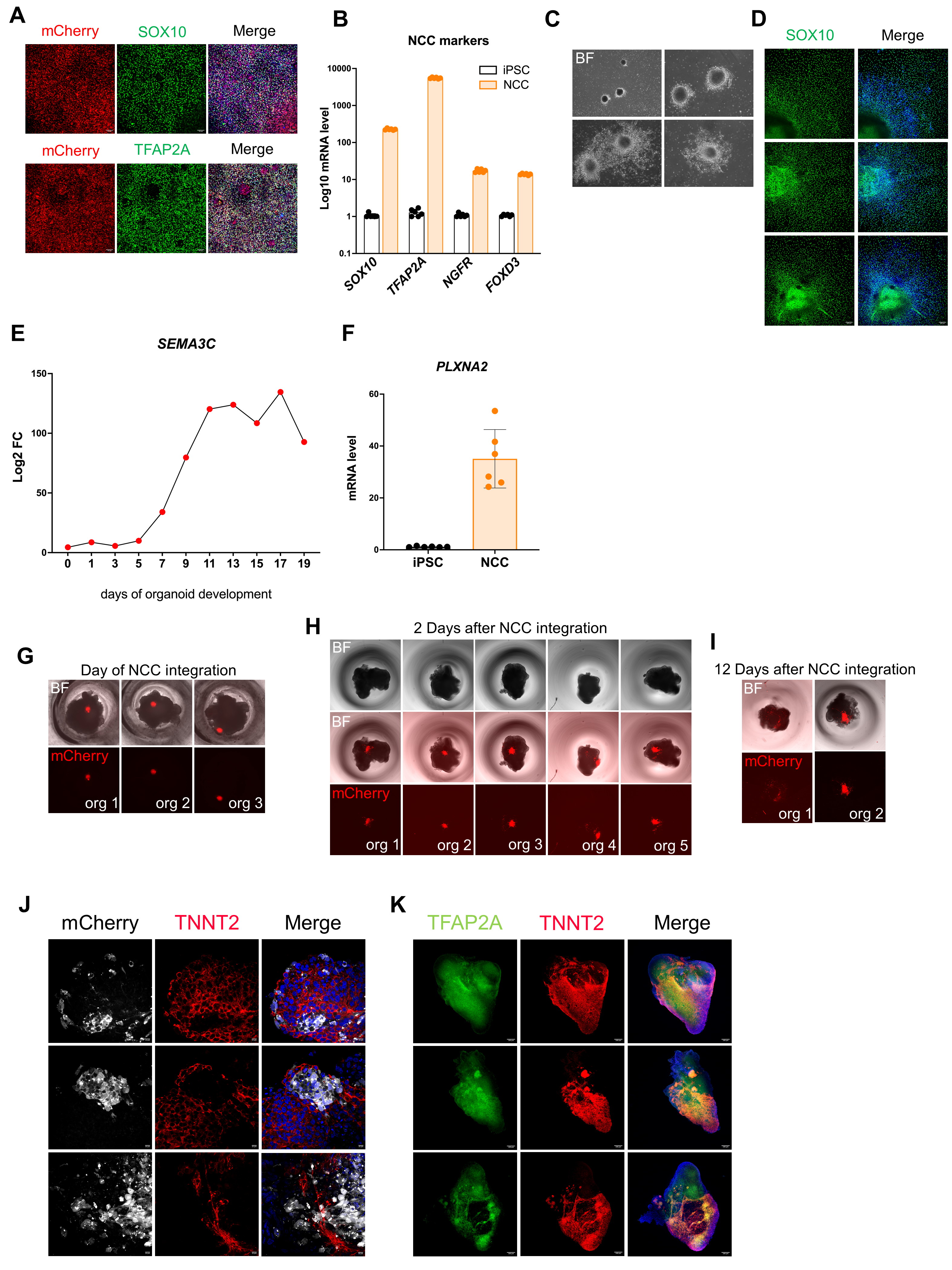

**Supplementary Figure 1**

(A) Representative immunofluorescence images iPSC-derived mCherry-NCCs (red) stained of SOX10 (green, top), TFAP2A (green, bottom) and the nuclear marker DAPI (blue). Scale bar = 50 mm.

(B) qRT-PCR gene expression analysis of key NCC markers in iPSC-derived NCCs; n = 6. Values = mean ± SD.

(C) Brightfield images of attached to Matrigel neurospheres with migrating NCCs.

(D) Representative immunofluorescence images of SOX10 (green) and the nuclear marker DAPI (blue) in NCCs migrating from neurospheres.

(E) Time-course gene expression analysis of SEMA3C between days 0 and 19 of heart organoid differentiation. Data presented as log2 fold-change normalized to day 0.

(F) qRT-PCR gene expression analysis of PLXNA2 in iPSC-derived NCCs; n = 6. Values = mean ± SD.

(G-I) Brightfield and fluorescent images of three hNCHAs on Day of NCC integration (G), five hNCHAs two days after NCC integration (I) and two hNCHAs twelve days after NCC integration (I).

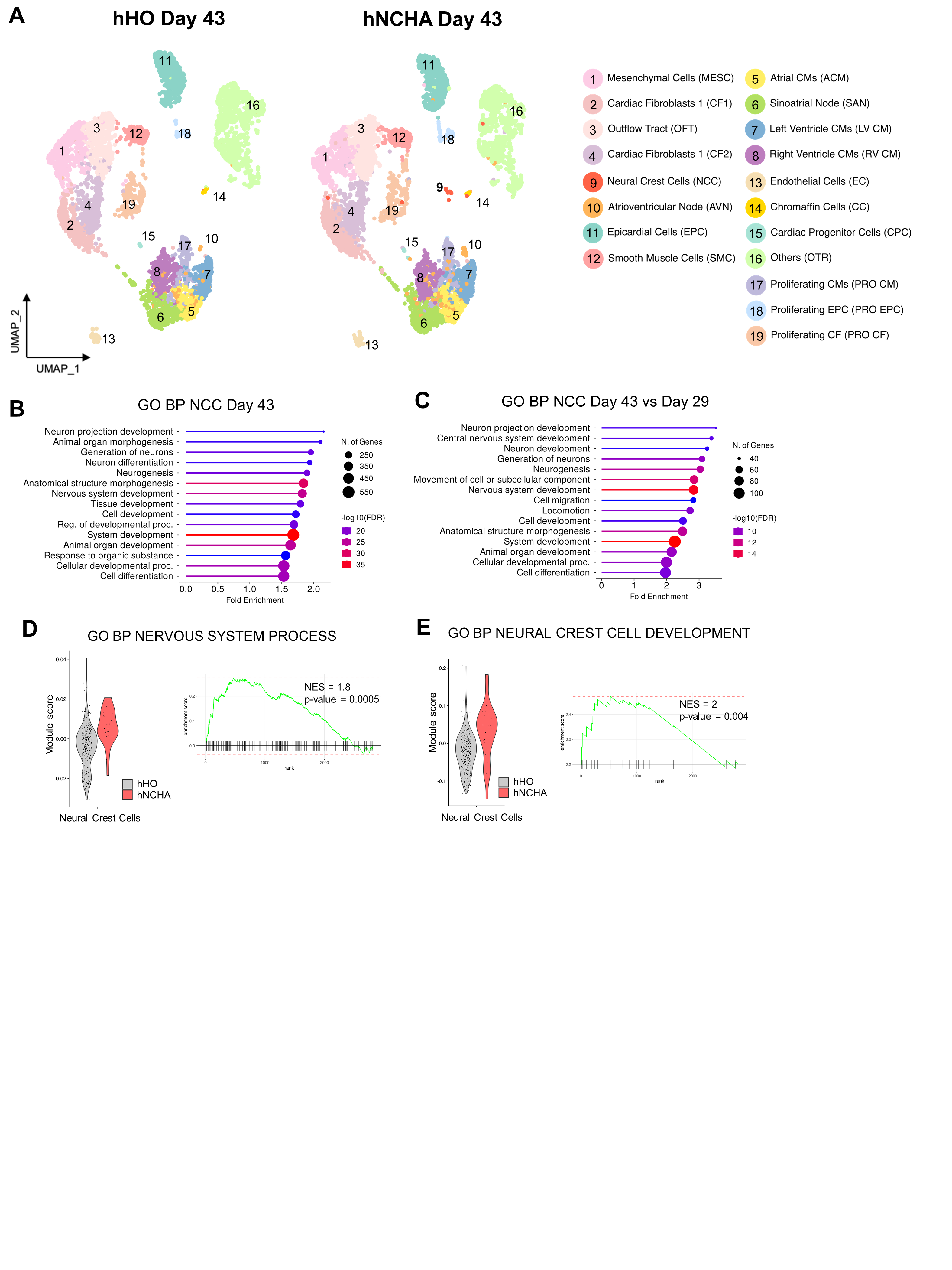

**Supplementary Figure 2**

(A) Uniform manifold approximation and projection (UMAP) plots of single-cell RNA sequencing data for Day 43 hHOs and Day 43 hNCHAs. Cluster identities are in the legend (right).

(B) Lollipop plots displaying top upregulated processes in NCCs of Day 43 hNCHAs versus NCCs before migration. GO – Gene Ontology; BP – Biological Process.

(C) Lollipop plots displaying top upregulated processes in NCCs of Day 43 hNCHAs versus Day 29 hNCHAs. GO – Gene Ontology; BP – Biological Process.

(D) (E) Pathway enrichment analysis in NCC cluster of Day 29 hNCHAs versus NCCs before migration. Violin plots depicting cells per sample by gene set module score. Enrichment plots of pathway-related gene sets overrepresented in NCC cluster of Day 29 hNCHAs versus NCCs before migration. NES, enrichment score normalized to mean enrichment of random samples of the same size (value represents the enrichment score after normalization).

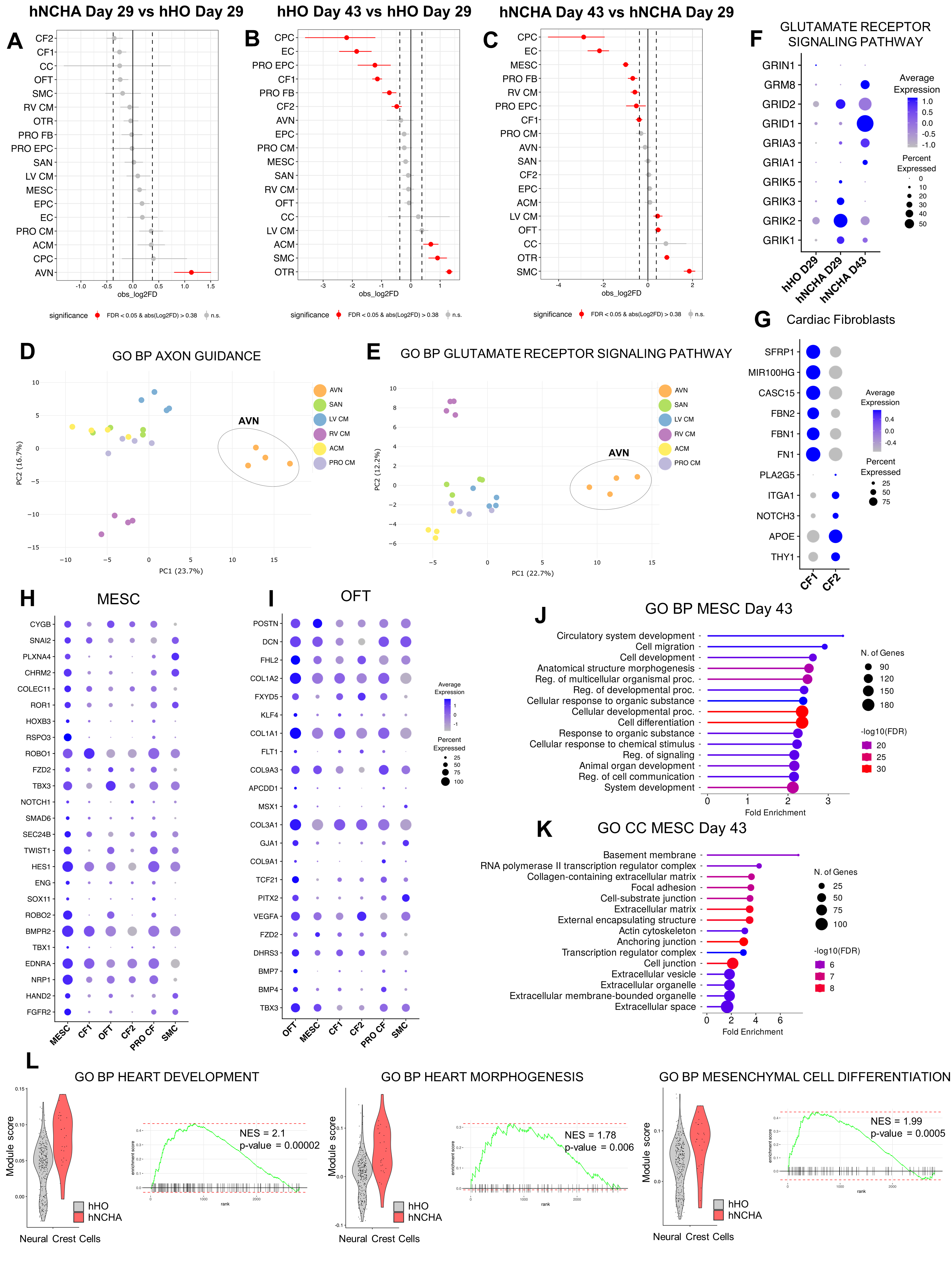

**Supplementary Figure 3**

(A) Relative differences in cell proportion for each cluster in Day 29 hNCHAs versus Day 29 hHOs. Clusters colored red have a false discovery rate <0.05 and mean |log2 fold enrichment| > 0.38 (permutation test; n = 10,000).

(B) Relative differences in cell proportion for each cluster in Day 43 hHOs versus Day 29 hHOs. Clusters colored red have a false discovery rate <0.05 and mean |log2 fold enrichment| > 0.38 (permutation test; n = 10,000).

(C) Relative differences in cell proportion for each cluster in Day 43 hNCHAs versus Day 29 hNCHAs. Clusters colored red have a false discovery rate <0.05 and mean |log2 fold enrichment| > 0.38 (permutation test; n = 10,000).

(D) A principal component analysis (PCA) plot illustrates the distinct clustering of the AVN cardiomyocyte population based on the expression of genes linked to axon guidance (GO:0007411; GO:1902667).

(E) A principal component analysis (PCA) plot illustrates the distinct clustering of the AVN cardiomyocyte population based on the expression of genes linked to glutamatergic receptor signaling pathway (GO:0007215; GO:0035249).

(F) Dot plot depicting genes associated with glutamatergic receptor signaling pathway (according to GO:0007215; GO:0035249) and differentially expressed in NCC cluster of Day 29 hNCHAs and Day 43 hNCHAs versus NCCs before migration. Dot size is proportional to the percentage of cells expressing specific genes. Color intensity indicates average expression (p-value<0.05).

(G) Dot plots depicting genes distinguishing CF1 and CF2 clusters. Dot size is proportional to the percentage of cells expressing specific genes. Color intensity indicates average expression (p-value<0.05).

(H) Dot plots depicting genes distinguishing MESC cluster from other mesenchymal clusters. Dot size is proportional to the percentage of cells expressing specific genes. Color intensity indicates average expression (p-value<0.05).

(I) Dot plots depicting genes distinguishing OFT cluster from other mesenchymal clusters. Dot size is proportional to the percentage of cells expressing specific genes. Color intensity indicates average expression (p-value<0.05).

(J) (K) Lollipop plots displaying top upregulated processes in MESC cluster of Day 43 hNCHAs versus Day 43 hHOs. GO – Gene Ontology; BP – Biological Process; CC – Cellular component.

(L) Pathway enrichment analysis in NCC cluster of Day 29 hNCHAs versus NCCs before migration. Violin plots depicting cells per sample by gene set module score. Enrichment plots of pathway-related gene sets overrepresented in NCC cluster of Day 29 hNCHAs versus NCCs before migration. NES, enrichment score normalized to mean enrichment of random samples of the same size (value represents the enrichment score after normalization).

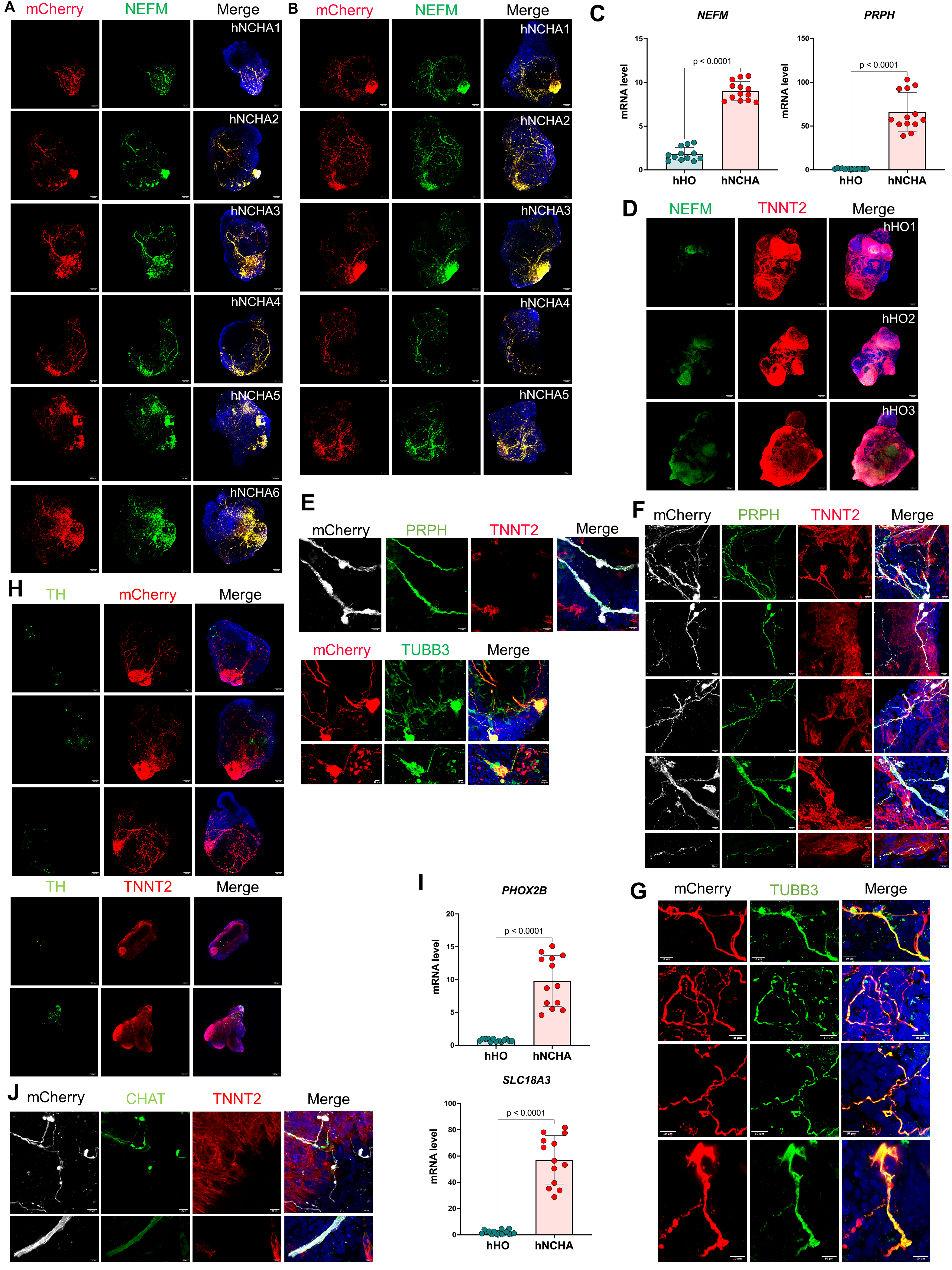

**Supplementary Figure 4**

(A) Representative immunofluorescence images of six Day 45 hNCHAs displaying mCherry-NCCs (grey), neuronal marker Neurofilament-160 (NEFM, green) and the nuclear marker DAPI (blue); Scale bar = 100 mm.

(B) Representative immunofluorescence images of five Day 60 hNCHAs displaying mCherry-NCCs (grey), neuronal marker Neurofilament-160 (NEFM, green) and the nuclear marker DAPI (blue); Scale bar = 100 mm.

(C) qRT-PCR gene expression analysis of NEFM and PRPH in Day 30 hHOs and Day 30 hNCHAs; n = 13. Values = mean ± SD.

(D) Representative immunofluorescence images of three Day 30 hHOs displaying neuronal marker Neurofilament-160 (NEFM, green), cardiomyocyte marker TNNT2 (red) and the nuclear marker DAPI (blue); Scale bar = 100 mm.

(E, top) Representative immunofluorescence image of Day 30 hNCHA displaying mCherry-NCCs (grey), neuronal marker Peripherin (PRPH green) and the nuclear marker DAPI (blue); Scale bar = 10 mm.

(E, bottom) Representative immunofluorescence images of two Day 30 hNCHAs displaying mCherry-NCCs (red), neuronal marker class III b-tubulin in ganglionated structures (TUBB3, green) and the nuclear marker DAPI (blue); Scale bar = 10 mm.

(F) Representative immunofluorescence images of four Day 45 hNCHAs displaying mCherry-NCCs (grey), neuronal marker Peripherin (PRPH, green), cardiomyocyte marker TNNT2 (red) and the nuclear marker DAPI (blue); Scale bar = 10 mm.

(G) Representative immunofluorescence images of four Day 45 hNCHAs displaying mCherry-NCCs (red), neuronal marker class III b-tubulin (TUBB3, green) and the nuclear marker DAPI (blue); Scale bar = 10 mm.

(H) Representative immunofluorescence images of three Day 30 hNCHAs and two Day 30 hHOs displaying sympathetic neuronal marker tyrosine hydrolase (TH, green), mCherry-NCCs (red) in hNHCAs or cardiomyocyte marker TNNT2 (red) in hHOs, and the nuclear marker DAPI (blue); Scale bar = 100 mm.

(I) qRT-PCR gene expression analysis of PHOX2B and SLC18A3 in Day 30 hHOs and Day 30 hNCHAs; n = 13. Values = mean ± SD.

(J) Representative immunofluorescence images of two Day 45 hNCHAs displaying mCherry-NCCs (grey), parasympathetic neuronal marker choline acetyltransferase (CHAT, green), cardiomyocyte marker TNNT2 (red) and the nuclear marker DAPI (blue); Scale bar = 10 mm.

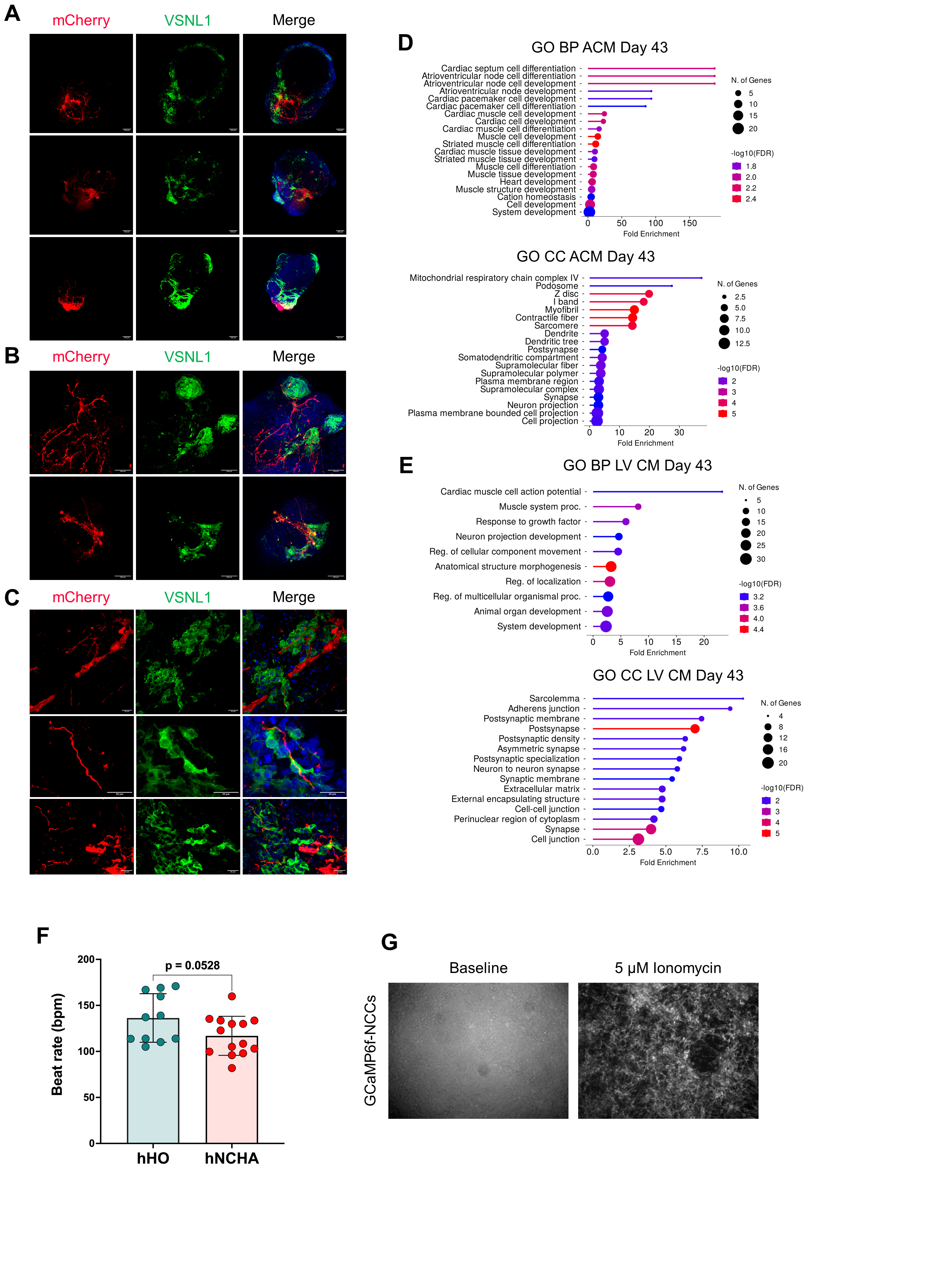

**Supplementary Figure 5**

(A) Representative immunofluorescence images of three Day 30 hNCHAs displaying mCherry-NCCs (red), sinoatrial node marker VSNL1 (green) and the nuclear marker DAPI (blue); Scale bar = 100 mm.

(B) Representative immunofluorescence images of three Day 45 hNCHAs displaying mCherry-NCCs (red), sinoatrial node marker VSNL1 (green) and the nuclear marker DAPI (blue); Scale bar = 100 mm.

(C) Representative high magnification immunofluorescence images of Day 45 hNCHAs displaying mCherry-NCCs (red), sinoatrial node marker VSNL1 (green) and the nuclear marker DAPI (blue); Scale bar = 20 mm.

(D) Lollipop plots displaying top upregulated processes in atrial cardiomyocytes of Day 43 hNCHAs versus Day 43 hHOs. GO – Gene Ontology; BP – Biological Process; CC – Cellular Component.

(E) Lollipop plots displaying top upregulated processes in left ventricle cardiomyocytes of Day 43 hNCHAs versus Day 43 hHOs. GO – Gene Ontology; BP – Biological Process; CC – Cellular Component.

(F) Quantification of beats per minute in Day 30 hHOs and Day 30 hNCHAs; n = 14. Values = mean ± SD, unpaired t test.

(G) Live imaging of calcium transients in GCaMP6f-NCCs before (left) and after (right) Ionomycin treatment, which induces spontaneous calcium activity.

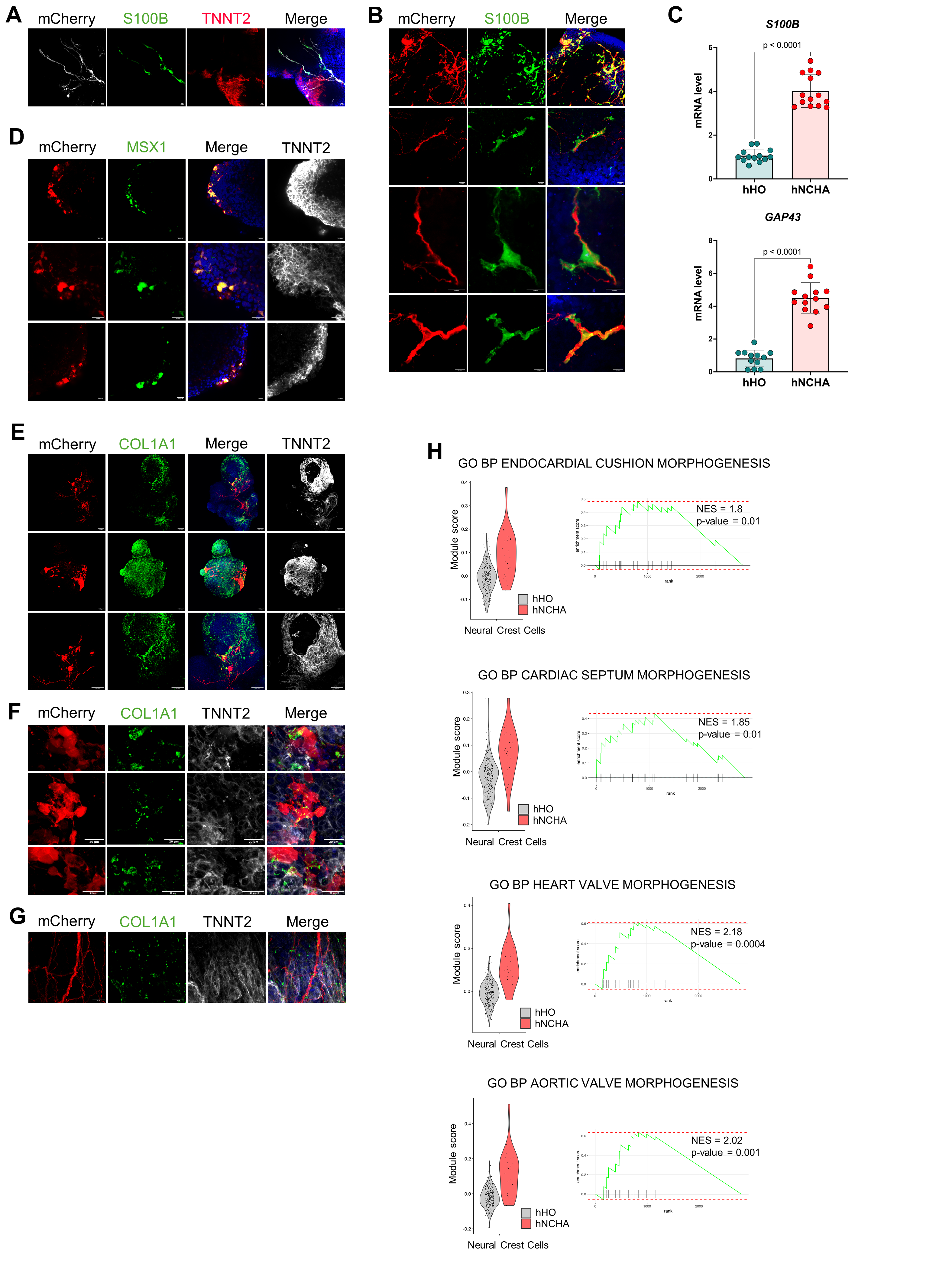

**Supplementary Figure 6**

(A) Representative immunofluorescence image of Day 30 hNCHAs displaying mCherry-NCCs (grey), glial marker (S100B, green), cardiomyocyte marker TNNT2 (red) and the nuclear marker DAPI (blue); Scale bar = 10 mm.

(B) Representative immunofluorescence images of Day 30 hNCHAs displaying mCherry-NCCs (red), glial marker (S100B, green) and the nuclear marker DAPI (blue); Scale bar = 10 mm and 20 mm (top).

(C) qRT-PCR gene expression analysis of S100B and GAP43 in Day 30 hHOs and Day 30 hNCHAs; n = 13. Values = mean ± SD.

(D) Representative immunofluorescence images of Day 30 hNCHAs displaying mCherry-NCCs (red), valve marker Msh Homeobox 1 (MSX1, green) and the nuclear marker DAPI (blue); Scale bar = 20 mm.

(E) Representative immunofluorescence images of Day 30 hNCHAs displaying mCherry-NCCs (red), valve marker Collagen 1A1 (COL1A1, green) and the nuclear marker DAPI (blue); Scale bar = 100 mm.

(F) Representative high magnification immunofluorescence images of Day 30 hNCHAs displaying mCherry-NCCs (red), valve marker Collagen 1A1 (COL1A1, green) and the nuclear marker DAPI (blue); Scale bar = 20 mm.

(G) Representative high magnification immunofluorescence image of Day 30 hNCHAs displaying mCherry-NCCs (red), valve marker Collagen 1A1 (COL1A1, green), cardiomyocyte marker TNNT2 (grey) and the nuclear marker DAPI (blue); Scale bar = 20 mm.

(H) Pathway enrichment analysis in NCC cluster of Day 29 hNCHAs versus NCCs before migration. Violin plots depicting cells per sample by gene set module score. Enrichment plots of pathway-related gene sets overrepresented in NCC cluster of Day 29 hNCHAs versus NCCs before migration. NES, enrichment score normalized to mean enrichment of random samples of the same size (value represents the enrichment score after normalization).

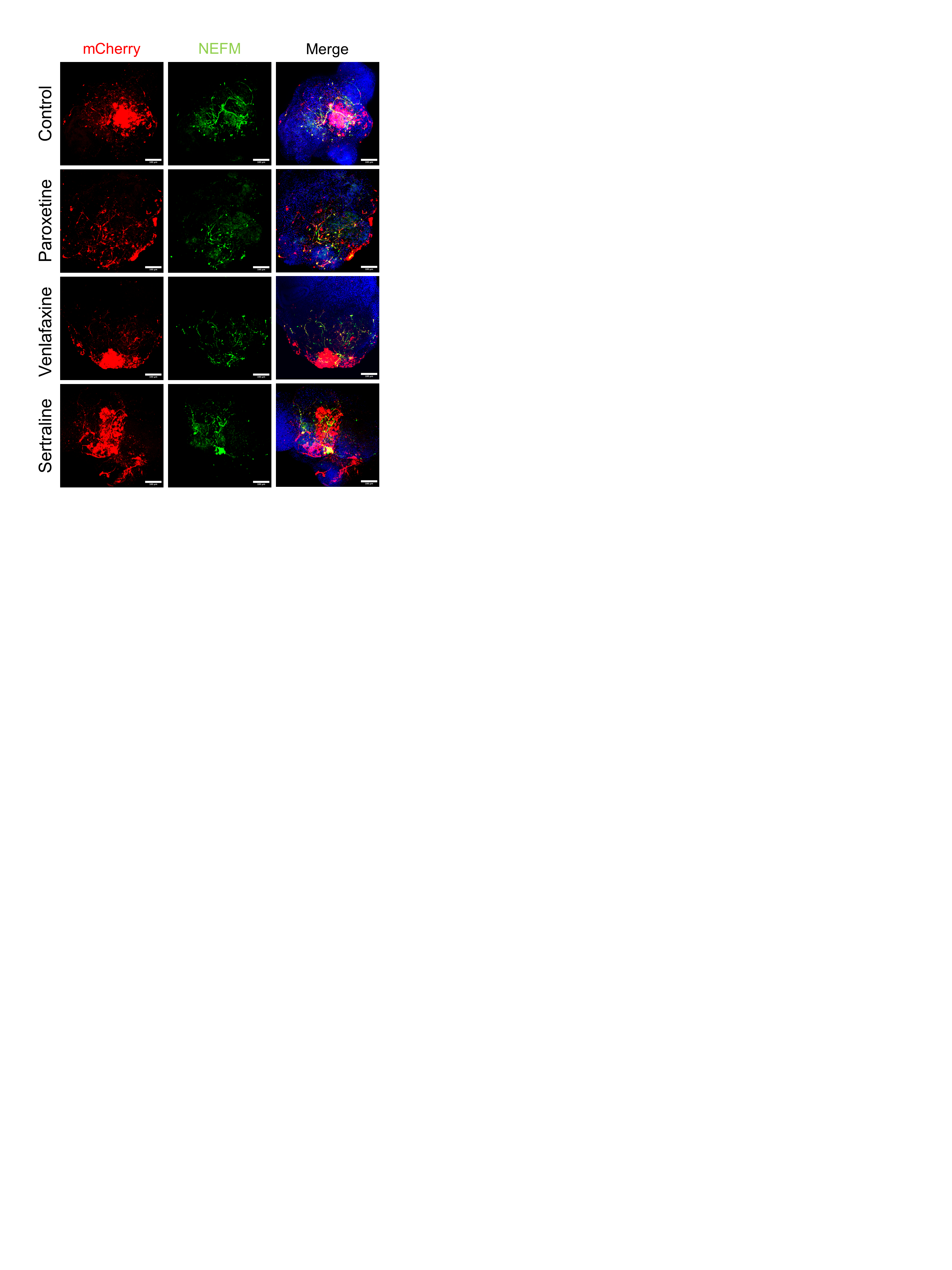

**Supplementary Figure 7**

Representative immunofluorescence images of Day 30 hNCHAs with healthy NCCs (C) and NCCs exposed to SSRIs during differentiation displaying mCherry-NCCs (red), neuronal marker NEFM (green) and the nuclear marker DAPI (blue); Scale bar = 100 mm.
